# A Premotor Excitation-Inhibition Motif Shapes Protopodium Dynamics During Drosophila Larval Forward Crawling

**DOI:** 10.64898/2026.01.23.701396

**Authors:** Ankura Sitaula, Lizzy Olsen, Dulanjana M. Perera, Yuhan Huang, Vashisht Errabolu Meher, Parinita Mitchelle Mandhyan, Nathan Byrd, Sara Arredondo, Lauren Carlisle, Yufeng Wan, Isuru S. Godage, Aref Zarin

## Abstract

Animals use shared muscles and motor neurons to generate distinct movements, but how premotor circuits differentially recruit these neuromuscular components remains unclear. *Drosophila* larvae crawl forward and backward using overlapping motor pools, yet the timing of ventral oblique (VO) muscle activity differs between these behaviors. Here, we show that VO muscles are major contributors to the movement of protopodia, limb-like structures that help larvae interact with the crawling surface. Muscle imaging, perturbation, and modeling indicate that VO activity is particularly important for protopodium folding, whereas contractions of both VO and ventral longitudinal (VL) muscles contribute to protopodium stride length and crawling efficiency. We identify a tri-segmental premotor motif—comprising excitatory (A27h, A18b3) and inhibitory (A06c) neurons—that shapes VO timing during forward crawling. A06c exhibits two activity peaks during forward fictive locomotion, defining a constrained window of VO MN activation, whereas during backward locomotion the early A06c peak is absent and the excitatory PMNs A27h and A18b3 are inactive. Disrupting this motif shifts VO activity earlier, reduces intersegmental timing delays, increases temporal overlap of VO activity, and impairs forward protopodium folding, stride, and crawling efficiency. These findings show how premotor excitation and inhibition coordinate muscle timing to shape locomotor mechanics.

## Introduction

Animals often generate multiple locomotor behaviors using the same sets of muscles and motor neurons (MNs). This flexibility raises a fundamental question in motor control: how do shared neuromuscular systems produce distinct motor outputs that satisfy different biomechanical demands? Addressing this problem requires understanding the biomechanical demands of a given motor behavior, determining which muscles and MNs are recruited and when, and identifying how upstream premotor circuits coordinate their timing across body regions.

The *Drosophila* larval locomotor system provides a powerful model for investigating this question. Larvae perform forward and backward crawling using largely overlapping MN pools and muscles, yet these behaviors differ in their kinematics and interactions with the substrate. Our previous work showed that ventral oblique (VO) muscles exhibit behavior-specific activation timing, activating later during forward crawling and earlier during backward crawling [1]. However, the functional significance of this timing difference and the premotor mechanisms that shape VO activity have remained unclear.

Larval crawling depends on repeated interactions between the body wall and the substrate, mediated in part by segmentally repeated, limb-like structures termed protopodia [2]. During both forward and backward crawling, protopodia undergo folding, swing, and stance-like phases, but the direction of folding and swing differs between behaviors [2]. Because muscle contraction pulls on both ends of a body-wall segment, productive protopodium movement likely requires a biomechanical mechanism that counteracts force transmission on one side of the contracting segment, allowing centerward muscle contraction to generate movement of only one of the two flanking protopodia. Thus, crawling direction imposes distinct biomechanical demands on how ventral muscle activity is timed within and across segments. However, how specific muscle groups across adjacent segments are activated and mechanically coordinated so that their forces generate direction-specific protopodium movement remains poorly understood.

VO muscles are well positioned to influence protopodium mechanics because they occupy the ventral body wall near the protopodia and exhibit behavior-specific timing during crawling [1, 3]. Yet their functional role in protopodium movement has not been directly tested using causal manipulations. Moreover, although multiple premotor neurons (PMNs) involved in larval crawling have been identified [1, 4–13], the temporal relationships among premotor excitation, inhibition, and downstream VO MN activity have not been fully characterized. In particular, it remains unclear how premotor activity is coordinated across adjacent segments to impose the forward-specific delay in VO activation, and how this premotor activity pattern differs during backward locomotion.

Here, we investigate how motor and premotor circuits control protopodium dynamics during larval crawling. We first combine muscle calcium imaging, optogenetic manipulation, and biomechanical modeling to test how VO and ventral longitudinal (VL) muscles contribute to protopodium folding, stride, and crawling efficiency. We then use connectome-based circuit analysis and functional perturbation to identify a tri-segmental premotor excitation-inhibition motif composed of the excitatory PMNs A27h and A18b3 and the inhibitory PMN A06c that shapes VO timing during forward crawling. During forward locomotion, A27h and A18b3 are active and A06c exhibits two activity peaks that bracket the VO activation window. Disrupting this motif shifts VO activity earlier, reduces intersegmental timing delays, increases temporal overlap of VO activity across segments, and impairs forward protopodium folding, stride, and crawling efficiency. During backward locomotion, A27h and A18b3 are inactive and the early A06c activity peak is absent, while A06c remains involved in VO coordination. Together, this approach links muscle mechanics, MN function, and premotor circuit organization to explain how premotor excitation and inhibition coordinate muscle timing to shape locomotor mechanics.

## Results

### Larval protopodia undergo walking-like swing and stance phases during forward and backward locomotion

The Drosophila larval body consists of three thoracic (T1–T3) and nine abdominal (A1–A9) segments, innervated by a correspondingly segmented ventral nerve cord (VNC) comprising 12 segments. Each abdominal segment contains ~30 bilaterally symmetric muscle pairs **(Figure 1A, Figure S1)** [3], innervated by ~30 glutamatergic excitatory MNs [1, 14–18]. During crawling, peristaltic waves of motor activity propagate posterior-to-anterior during forward locomotion and anterior-to-posterior during backward locomotion **(Figure 1)**. Both behaviors are bilaterally synchronous, with left and right muscles contracting simultaneously within each segment.

**Figure 1.**
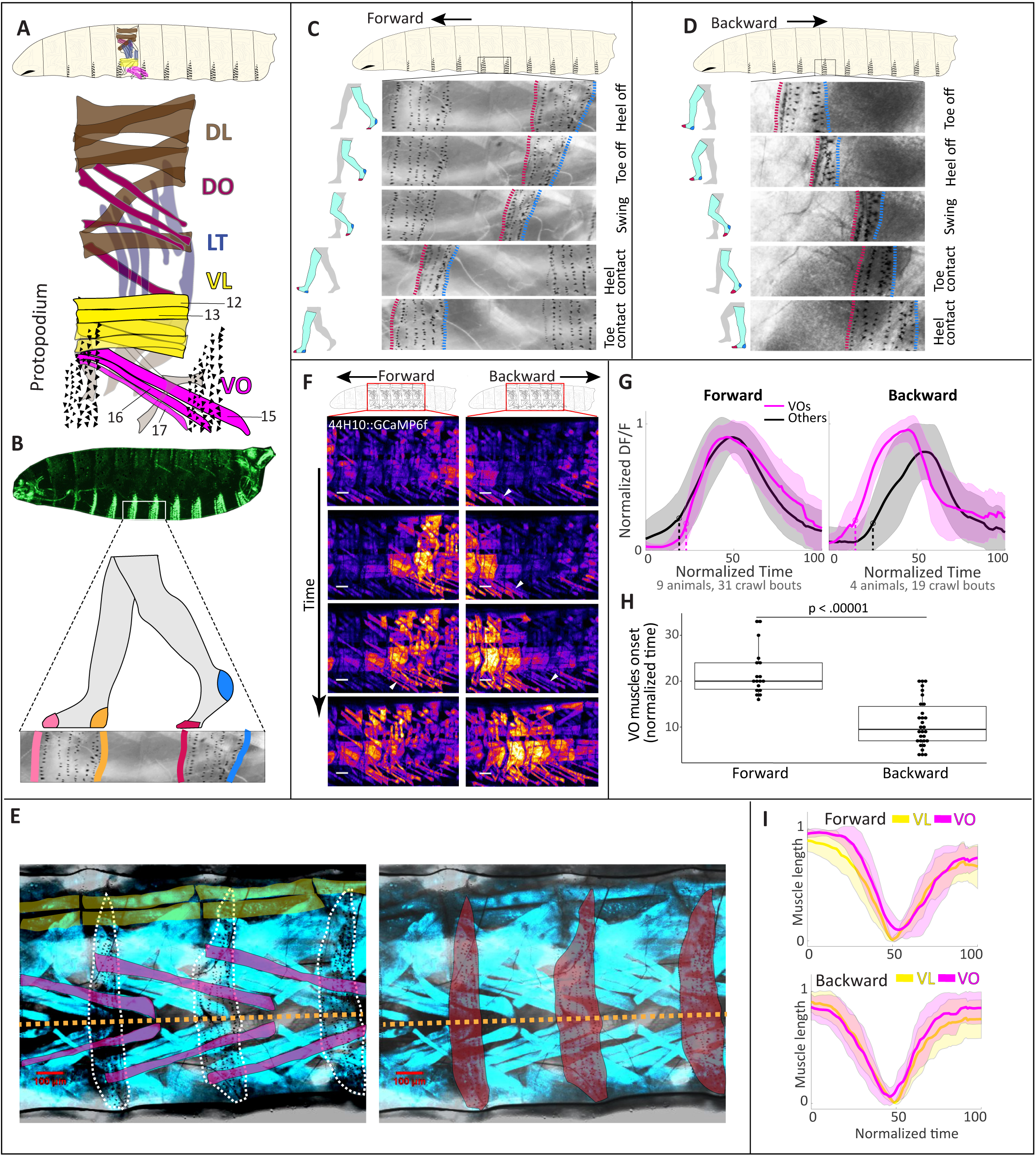
Segmental protopodia and VO muscles show direction-specific movement and activation during crawling. **(A)** Schematic of the Drosophila larval body plan showing 3 thoracic and 9 abdominal segments. Each abdominal segment contains a ventrally positioned protopodium. Muscle layout of segments A2–A6 highlights ventral oblique (VO) muscles 15–17, which span adjacent segments and anatomically lie adjacent to the protopodia. **(B)** The protopodium is a protruding, foot-like ventral structure composed of 6–8 rows of denticle bands. The posterior-most and anterior-most bands are referred to as the “heel” and “toe,” respectively. Whole-larva image adapted from the FlyMove [67]. **(C)** During forward crawling, the protopodium of the actively contracting segment folds in a heel-to-toe sequence, detaches from the substrate, and enters the swing phase translocating forward. After the swing phase, it recontacts the substrate and unfolds in a heel-to-toe order, analogous to human forward walking. **(D)** During backward crawling, the sequence of protopodium folding and unfolding is reversed (toe-to-heel), resembling backward walking. **(E)** Ventral view of larval body wall showing muscle and protopodium anatomy. VO muscle 15 (magenta) and ventral longitudinal (VL) muscles 12 and 13 (yellow) are labeled. Protopodia are outlined (left) or shaded red (right). The dashed orange line marks the ventral midline. *Genotype: 44H10-lexA, AOP-mCherry.* **(F)** Time-lapse images of VO muscle calcium activity visualized with GCaMP6f during forward (left) and backward (right) crawling. Arrowheads indicate active VO muscles. Scale bar: 50 µm. *Genotype: 44H10-GCaMP6f.* **(G)** GCaMP6f traces show earlier VO muscle activation during backward crawling. Magenta traces indicate VO muscle activity; black traces indicate combined activity of other muscles in the same segment. Time is normalized to a full peristaltic cycle (0–100%) using a PCA-based alignment method [1]. Sample sizes: forward crawling, 9 animals and 31 crawl bouts; backward crawling, 4 animals and 19 crawl bouts. **(H)** VO onset timing is significantly earlier during backward crawling. Each dot represents one VO muscle. Student’s *t*-test, *p* < 0.00001. **(I)** Normalized muscle length measurements show that VL muscles contract before VO muscles during forward crawling, whereas both contract nearly simultaneously during backward crawling.

Larvae interact with the substrate via segmentally repeated, foot-like structures termed protopodia, which are composed of 6–8 rows of actin-rich denticles located on the ventroanterior surface of each segment **(Figure 1B–D, Figure S1)** [2]. To describe their ground-contact dynamics, we use a functional analogy to the human foot, referring to posterior and anterior denticle rows as “heel” and “toe,” respectively; this analogy reflects shared biomechanical principles rather than anatomical homology.

Consistent with recent kinematic analyses by Booth et al. [2], during both forward and backward crawling the protopodium of the actively contracting segment folds and then enters a swing phase, lifting off the substrate and translocating in the direction of movement, while protopodia in adjacent, non-contracting segments remain in stance **(Figure 1B–D, Figure S1, Video S1)**. During forward crawling, protopodium folding proceeds in a heel-to-toe sequence **(Figure 1C, Video S1)**, whereas during backward crawling this sequence is reversed, proceeding toe-to-heel **(Figure 1D, Video S1)**. Following the swing phase, the protopodium recontacts the substrate and unfolds in a heel-to-toe sequence during forward crawling and toe-to-heel during backward crawling. Together, these observations demonstrate that larval crawling exhibits direction-specific protopodial kinematics analogous to forward and backward walking in humans [19–24].

### Ventral oblique muscle activity correlates with behavior-specific protopodium movement

We next sought to identify the muscles that contribute to protopodium movement. Forward and backward crawling exhibit distinct patterns of protopodium folding, suggesting that the muscles driving protopodial progression are differentially timed across behaviors. The protopodium occupies the ventral plane of the larval body wall, directly overlying the VO muscles 15–17 and ventral acute (VA) muscle 29. Its lateral edges partially extend into the ventrolateral region containing ventral longitudinal (VL) and VA muscles 26–27 **(Figure 1E)**. Consistent with our prior work [1], calcium imaging revealed that VO muscles are among the earliest activated muscle groups during backward crawling, whereas during forward crawling they activate later in the cycle, following VL and VA muscles **(Figure 1F–H, Figure S2, Video S2)**. Muscle length measurements further showed that VL muscles shorten before VO muscles during forward crawling, while they contract nearly synchronously during backward crawling **(Figure 1I)**.

VO muscles in a given segment **n** are positioned between two adjacent protopodia, n and n+1. Thus, based on their anatomical proximity to the protopodia and their differential activity patterns during forward and backward crawling, we hypothesized that VO muscles play a key role in controlling protopodium motion. We therefore sought to determine whether and how VO contraction selectively drives progression of only one neighboring protopodium per cycle.

To address this, we first performed simultaneous calcium imaging of VO muscles and high-resolution imaging of their flanking protopodia, n and n+1, in intact behaving larvae **(Figure 2A– C, Video S3)**. During forward crawling, peak VO activity in segment n coincided with folding and swing of protopodium n+1, while protopodium n remained in stance. During backward crawling, the opposite relationship was observed: peak VO activity in segment **n** coincided with folding and swing of protopodium n, while protopodium n+1 remained in stance **(Figure 2A–C, Video S3)**.

**Figure 2.**
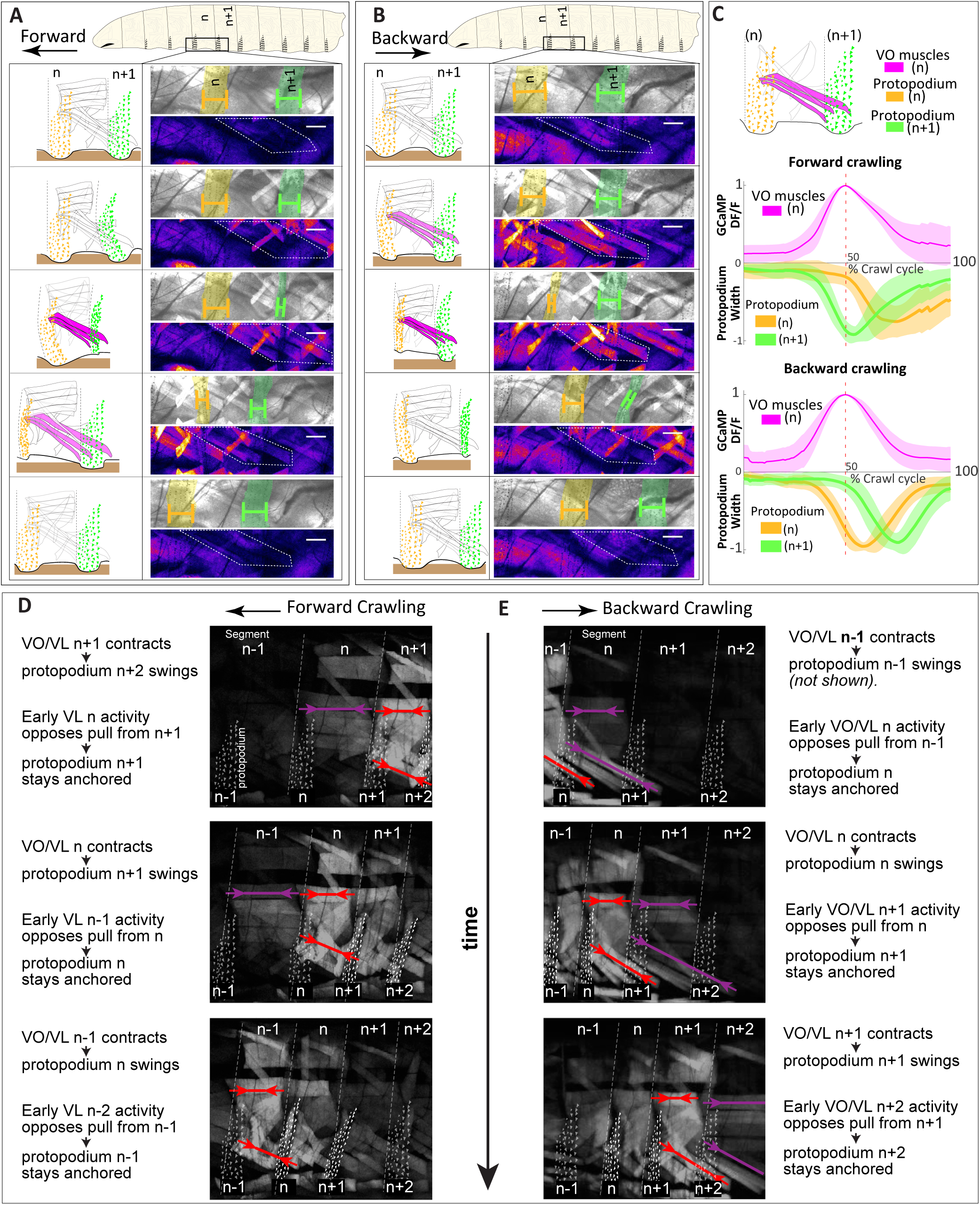
Behavior-specific VO/VL timing coordinates protopodium swing and anchoring during forward and backward crawling. **(A,B)** Schematics and time-lapse images illustrating the relationship between VO muscle activity and protopodium movement during forward **(A)** and backward **(B)** crawling. Scale bars: 50 μm. During forward crawling, peak VO activity in segment *n* coincides with folding and swing of the posterior protopodium, *n+1*. During backward crawling, peak VO activity in segment *n* coincides with folding and swing of the anterior protopodium, *n*. **(C)** VO muscle calcium activity and protopodium width plotted across a normalized crawl cycle, aligned to peak VO activity in segment *n*. Protopodium width was used as a proxy for folding associated with swing. During forward crawling, peak VO activity coincides with minimal width of protopodium *n+1*, whereas during backward crawling it coincides with minimal width of protopodium *n*. Sample sizes: forward crawling, 16 bouts from 7 animals; backward crawling, 17 bouts from 7 animals. Genotype: *44H10-GCaMP6f*. **(D,E)** Mechanical model for direction-specific protopodium movement generated by VO/VL timing and neighboring-segment anchoring. See **Video S4**. **(D)** During forward crawling, early VL activity in the next anterior segment is proposed to oppose force transmission and stabilize the anterior protopodium in stance, allowing VO/VL contraction in the active segment to fold and swing the posterior protopodium. **(E)** During backward crawling, early VO/VL activity in the next posterior segment is proposed to oppose force transmission and stabilize the posterior protopodium in stance, allowing VO/VL contraction in the active segment to fold and swing the anterior protopodium. Red arrows indicate VO/VL contraction in the active segment. Purple arrows indicate early neighboring VL or VO/VL activity proposed to oppose pull and increase resistance to stretch. Dashed outlines mark protopodia. *P n* denotes the protopodium of segment *n*. The vertical arrow indicates time.

To explain how differential VO activation timing could contribute to protopodium movement during forward and backward crawling, we propose a mechanical model based on the relative timing of VL and VO activity, the oblique geometry of VO muscles, and the anchoring state of neighboring protopodia **(Figure 2D, Video S4)**. When a muscle group contracts within a segment, it exerts pulling forces at its attachment sites, drawing the anterior and posterior sides of the segment toward the muscle or segment center. Thus, productive protopodium movement requires an asymmetric mechanical balance: centerward force transmission must be counteracted on one side of the contracting segment so that one flanking protopodium remains anchored in stance while the other folds and swings.

In the absence of such counterbalancing, contraction of VL and VO muscles in segment n would be expected to pull both protopodium n and protopodium n+1 toward the segment center. For efficient forward crawling, the pull on protopodium n must be opposed so that it remains anchored in stance, while protopodium n+1 is free to move. We propose that this opposition arises, at least in part, from early activity of VL muscles in the next anterior segment, n−1. At this phase of the crawl cycle, VL but not VO muscles in segment n−1 show an early calcium rise but do not undergo obvious shortening **(Figure 2D)**. This partial VL activation in segment n−1 may increase muscle tone and resistance to stretch. This mechanically resistant state could counteract force transmission from segment n, thereby helping stabilize protopodium n in stance. At the same time, VO contraction in segment n can fold, lift, and swing protopodium n+1. Thus, during forward crawling, delayed VO activation relative to VL activation in the adjacent anterior segment may help convert centerward force transmission within segment **n** into posterior protopodium swing.

Similarly, during backward crawling, contraction of VO and VL muscles in segment n would pull both flanking protopodia toward the segment center unless this force transmission is counterbalanced. In contrast to forward crawling, productive backward crawling requires the pull on protopodium n+1 to be opposed, so that protopodium n+1 remains anchored in stance while protopodium n is free to move **(Figure 2E)**. We propose that this opposition arises, at least in part, from early VO/VL activity in the next posterior segment, n+1.

At this phase of the backward crawl cycle, VO and VL muscles in segment n+1 are activated nearly synchronously, with VO activity sometimes occurring slightly earlier, while segment **n** is undergoing contraction. The geometry of VO muscles may make the VO component of this early posterior activity especially well-suited to provide mechanical resistance during backward crawling. VO muscles are obliquely oriented: their anterior attachments lie near the anterior boundary of their segment of origin, whereas their posterior extent reaches toward the next posterior protopodial region. Thus, simultaneous with VO/VL contraction in segment n, early VO/VL activation in segment n+1 may increase muscle tone and resistance to stretch. This mechanically resistant state could counteract force transmission from segment n, thereby stabilizing protopodium n+1 in stance as VO contraction in segment **n** folds, lifts, and swings the anterior protopodium n. Thus, early VO/VL activation during backward crawling may help convert centerward force transmission within segment **n** into anterior protopodium swing.

Together, these observations link behavior-specific VO/VL timing to direction-specific protopodium mechanics. They suggest that VO muscles do not act in isolation, but within a direction-dependent mechanical context established by neighboring protopodial anchoring, muscle tone, VO/VL timing, VO muscle geometry, and substrate contact. This provides a plausible mechanical basis by which VO contraction in the same segment can be associated with posterior protopodium swing during forward crawling and anterior protopodium swing during backward crawling **(Figure 2, Video S3 and Video S4)**.

### Mechanical modeling predicts a major mechanical contribution of VO muscles to protopodium gait generation

To examine how VO and VL muscles mechanically contribute to protopodium movement during crawling, we developed a kinematic model that builds upon and extends a previously published framework. Because larval crawling occurs primarily in the sagittal plane and ventral muscle anatomy is bilaterally symmetric, we modeled the VO–VL–protopodium system as a planar four-bar mechanism, with elastic elements representing deformable muscle-linked body-wall segments **(Figure 3A; Figure S3)**. This simplified representation was supported by in vivo tracking of ventral muscle and protopodium trajectories during forward and backward crawling, which showed limited out-of-plane motion **(Figure 3B)**.

**Figure 3.**
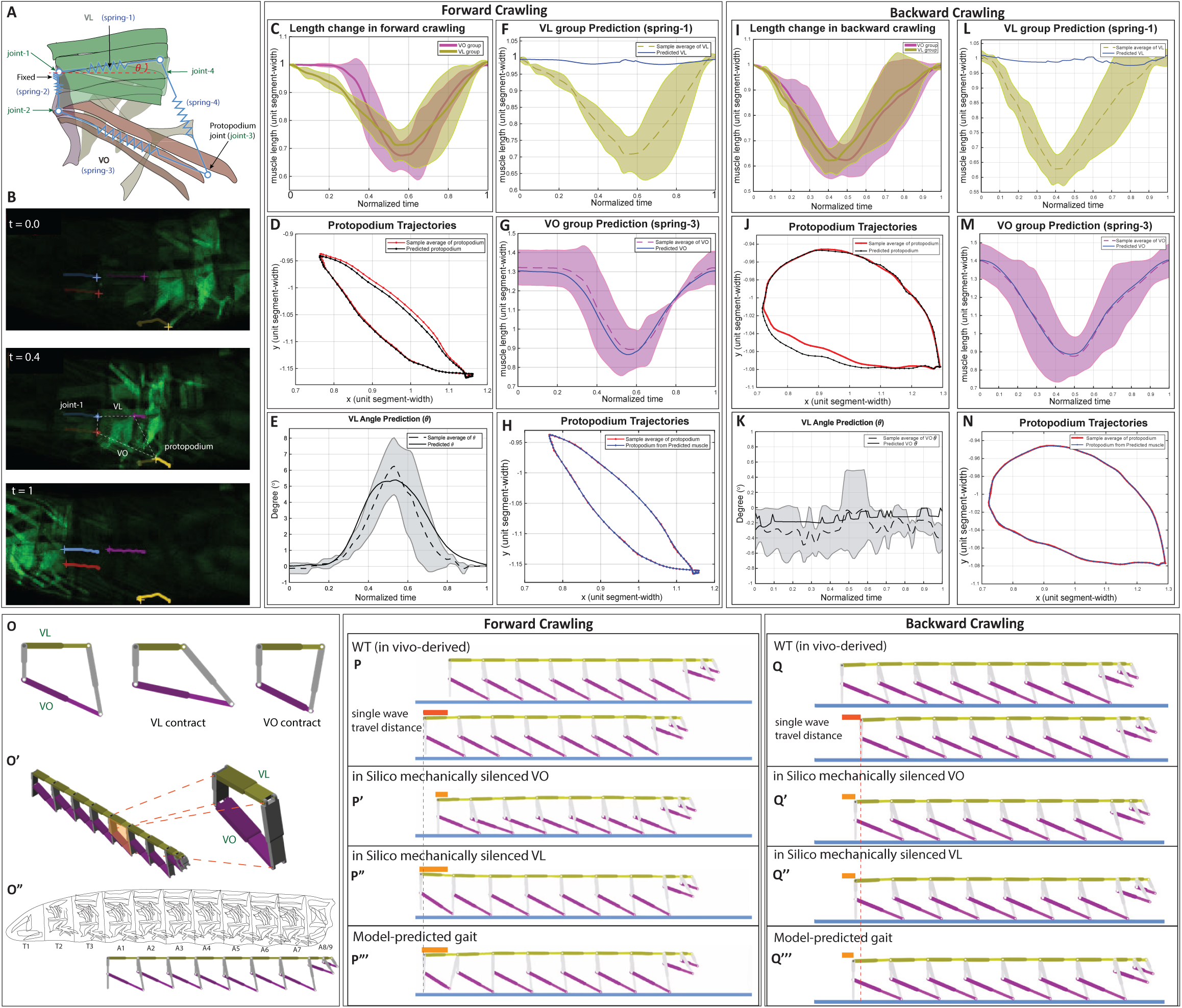
Kinematic and robotic models identify VO muscles as major drivers of direction-specific protopodium movement. **(A)** Four-bar kinematic model of one abdominal segment. VL and VO muscles are represented by spring-1 and spring-3; spring-2 and spring-4 maintain four-bar geometry. Protopodium-joint motion represents segmental protopodium gait. **(B)** Representative forward-crawling bout showing in vivo tracking of VO, VL, and the protopodium joint. Colored traces mark tracked points used for modeling. **(C–H)** Forward-crawling kinematic model. **(C)** In vivo-measured normalized VO and VL length dynamics; solid lines indicate means and shaded regions indicate ± s.d. **(D)** Measured protopodium trajectory compared with the trajectory predicted from measured VO/VL lengths and VL angle. **(E–G)** Inverse-model predictions of VL angle, VL length, and VO length compared with experimental mean ± s.d. Predicted VO length dynamics more closely matched the observed VO length-change profile, whereas predicted VL shortening was smaller than observed in vivo. **(H)** Protopodium trajectory regenerated from model-predicted muscle dynamics. **(I–N)** Backward-crawling kinematic model. **(I)** In vivo-measured normalized VO and VL length dynamics. **(J)** Measured backward protopodium trajectory compared with the trajectory predicted from measured muscle dynamics. **(K–M)** Inverse-model predictions of VL angle, VL length, and VO length compared with experimental mean ± s.d. As in forward crawling, predicted VO length dynamics more closely matched the observed VO length-change profile than predicted VL dynamics. **(N)** Backward protopodium trajectory regenerated from model-predicted muscle dynamics. **(O–O″)** Robotic model composed of repeated four-bar modules. **(P–P‴)** Forward robotic simulations. WT in vivo-derived muscle dynamics generated forward displacement; VO silencing markedly reduced displacement; VL silencing preserved displacement but caused skidding; model-predicted muscle dynamics also generated forward movement. **(Q–Q‴)** Backward robotic simulations. WT in vivo-derived muscle dynamics generated backward displacement, whereas in silico silencing of either VO or VL reduced displacement. Simulations driven by model-predicted muscle dynamics produced still lower displacement, likely reflecting features not fully captured by the current model, including passive tissue deformation and substrate grip. Together, the models support a major mechanical contribution of VO muscles to direction-specific protopodium movement, while indicating that VL muscles and substrate-contact mechanics also shape displacement.

Using experimental tracking data from five larvae, we first quantified VO and VL muscle length changes, VL horizontal angle, and protopodium trajectories during forward crawling **(Figure 3C–H)** and backward crawling **(Figure 3I–N)**. Consistent with calcium imaging and muscle-length measurements in **Figure 1**, VL muscles shortened before VO muscles during forward crawling, whereas VO and VL shortening were more closely synchronized during backward crawling **(Figure 3C, I)**. In this kinematic model, muscle length changes and VL angular displacement were used to represent the geometric constraints imposed by ventral body-wall deformation.

We first asked whether experimentally measured muscle kinematics were sufficient to reproduce the observed forward protopodium trajectory. When measured VO/VL length changes and VL angle were provided as model inputs, the predicted protopodium trajectory closely resembled the experimentally measured trajectory, with minor offsets likely reflecting tissue deformation, substrate contact, or other body-wall forces not included in the simplified model **(Figure 3D)**. Thus, the model captured key geometric relationships linking ventral muscle dynamics to protopodium movement during forward crawling.

We next inverted the model by providing the experimentally measured protopodium trajectory as the input and asking which muscle dynamics would be predicted to generate it. The predicted VL angular trajectory followed the experimentally observed temporal profile **(Figure 3E)**. However, the predicted VL length change was substantially smaller than the VL shortening measured in vivo **(Figure 3F)**, whereas the predicted VO length change more closely matched the experimentally measured VO dynamics **(Figure 3G)**. Additional inverse-model predictions for all four spring elements are shown in **Figure S4**. When these predicted muscle dynamics were used to regenerate the protopodium trajectory, the simulated trajectory closely reproduced the experimental gait **(Figure 3H)**. These results suggest that, within the geometric constraints of the model, VO shortening together with VL rotation is sufficient to account for much of the forward protopodium trajectory, whereas large VL shortening is not strictly required for protopodium gait generation.

We applied the same modeling approach to backward crawling. Experimentally measured VO/VL muscle dynamics **(Figure 3I)** generated a predicted backward protopodium trajectory that closely matched the measured trajectory **(Figure 3J)**. In the inverse model, the predicted VL angular trajectory was relatively small and variable **(Figure 3K)**, and the predicted VL length change was again smaller than the VL shortening measured in vivo **(Figure 3L)**. By contrast, the predicted VO length change more closely matched the experimentally observed VO dynamics **(Figure 3M)**. Using the predicted muscle dynamics to regenerate the backward protopodium trajectory again reproduced the measured trajectory **(Figure 3N)**. Thus, the kinematic model could represent both forward and backward protopodium gaits and consistently predicted a larger contribution of VO shortening than VL shortening to protopodium trajectory generation.

To further test these predictions in a physics-based context, we implemented a segmented robotic model composed of repeated four-bar mechanisms interacting with a frictional substrate **(Figure 3O–O″, Video S5)**. This model was not designed to quantitatively reproduce all aspects of larval crawling, including free-crawling speed, absolute cycle duration, whole-animal displacement, or the contributions of all body-wall muscles. Instead, it provided an independent test of whether the muscle dynamics identified by the kinematic model could generate effective segmental displacement under simplified substrate-contact conditions.

Under control conditions derived from in vivo measurements, the forward-crawling robotic model advanced by approximately 70% of a relaxed segment width per peristaltic cycle **(Figure 3P)**. In silico mechanical silencing of VO contraction markedly reduced forward displacement **(Figure 3P′)**. In contrast, in silico mechanical silencing of VL contraction preserved much of the forward displacement, although skidding was observed, likely due to reduced VL-linked body-wall deformation and rearward force balance (**Figure 3P″)**. Simulations driven by the muscle dynamics predicted by the kinematic model produced forward displacement comparable to the VL-silenced condition **(Figure 3P‴)**. These results support the prediction that VO contraction plays a major mechanical role in generating effective forward protopodium gait, whereas VL contraction contributes to broader body-wall mechanics that are not fully captured by protopodium displacement alone.

We next examined backward crawling in the physics-based simulation **(Figure 3Q–Q‴)**. Under control conditions derived from in vivo measurements, the backward-crawling robotic model advanced by approximately 53% of a relaxed segment width per peristaltic cycle **(Figure 3Q)**. Although this simulation produced effective backward displacement, the magnitude of displacement was sensitive to the simplified substrate-contact geometry. In larvae, the ventral body wall maintains broad contact with the substrate, whereas the robotic model interacts with the substrate through more restricted contact points. This difference is expected to have a larger effect during backward crawling, in which oblique VO-mediated pulling likely depends more strongly on protopodial grip or adhesion.

Nevertheless, in silico silencing of either VO or VL reduced backward displacement relative to the control condition **(Figure 3Q′–Q″)**, suggesting that backward displacement in this simplified simulation depends on both ventral muscle components. Simulations driven by model-predicted muscle dynamics produced still lower backward displacement **(Figure 3Q‴)**, likely reflecting features not fully captured by the current model, including distributed passive tissue deformation, detailed substrate grip, and the mechanical effects of neighboring body-wall muscles.

Together, the physics-based simulations support the prediction that VO muscles make a major mechanical contribution to effective protopodium gait, while also suggesting that VL muscles contribute to aspects of protopodium displacement, particularly during backward crawling. At the same time, the robotic model is intentionally simplified and is best interpreted as a test of mechanical sufficiency rather than a complete quantitative reconstruction of larval locomotion.

### VO muscles receive input from both tonic and phasic motor neurons

In *Drosophila* larvae, glutamatergic excitatory MNs are broadly classified into phasic Is MNs, which form small synaptic boutons, and tonic Ib MNs, which form larger boutons on their target muscles [15–17, 25–27]. Previous retrograde labeling by Mauss et al. identified two pairs of tonic Ib MNs that innervate VO muscles 15–17: MN15/16, which innervates VO15 and VO16, and MN15–17, which innervates all three VO muscles [17].

We cross-validated these assignments by identifying genetic drivers that label VO MNs and characterizing their dendritic arborization in the VNC and their peripheral innervation patterns at the muscle field. To access tonic VO MNs, we used three genetic drivers: vGlut-AD∩Nkx6-DBD split-Gal4, NB7-1-Gal4, and FD4-Gal4. The vGlut-AD∩Nkx6-DBD split-Gal4 driver labels MN15/16, MN15–17, and several VL MNs [28, 29]. NB7-1-Gal4 labels MN15–17 and five tonic MNs innervating dorsal longitudinal (DL) muscles [30] **(Figure 4, Figure S5-S7)**.

**Figure 4.**
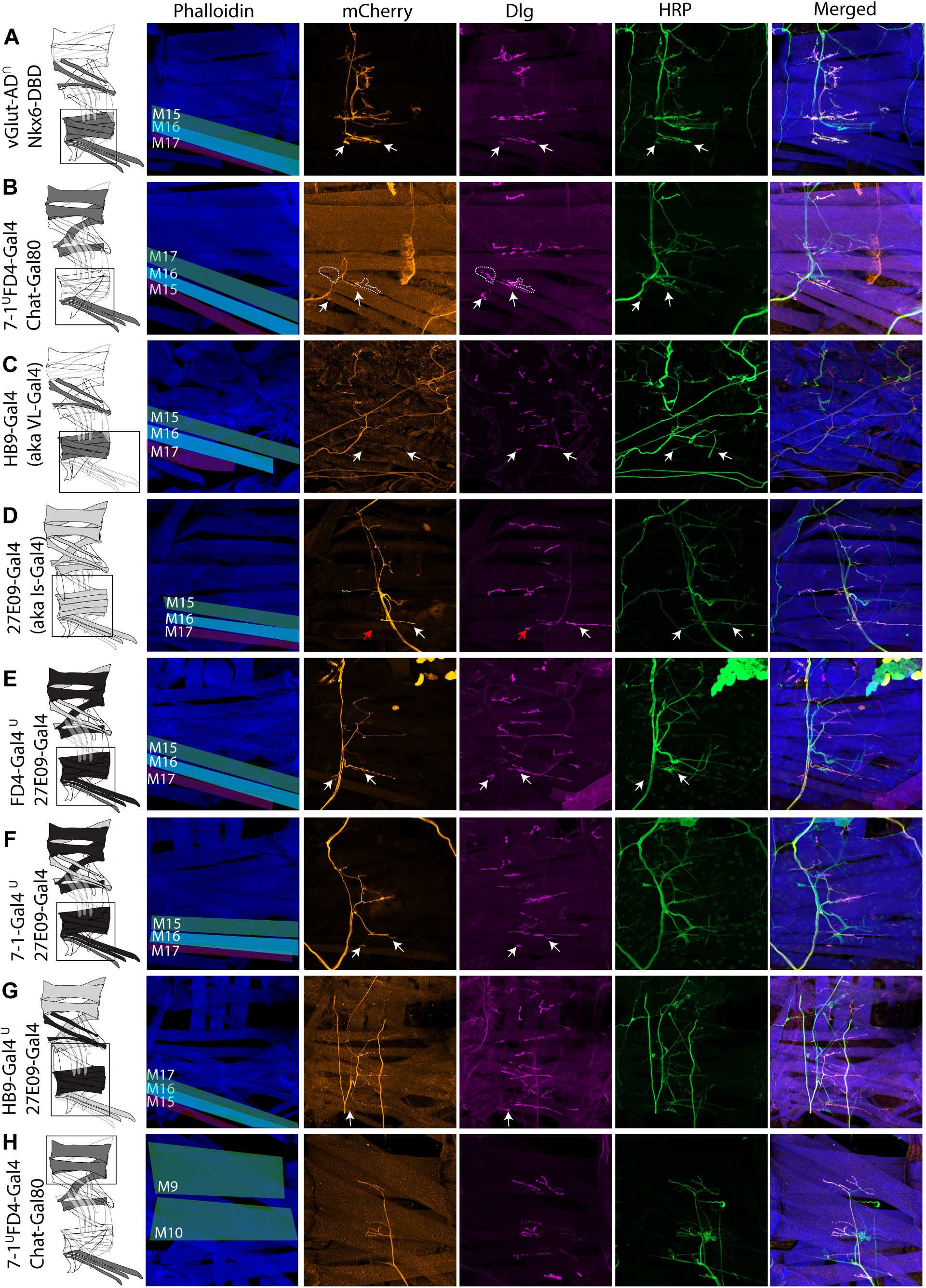
Peripheral expression patterns and muscle targets of MN-Gal4 lines used in this study. Representative confocal images of larval body wall muscle fillets show the peripheral expression patterns of MN-Gal4 and split-Gal4 drivers. mCherry (orange) reports Gal4 expression; phalloidin (blue) labels muscles; anti-HRP (green) labels neuronal membranes; and Discs large (Dlg, magenta) marks postsynaptic densities at tonic and phasic neuromuscular junctions (NMJs). Merged images are shown at right. Schematics at left summarize the muscle groups innervated by MN drivers. **(A)** *vGlut-AD^∩^Nkx6-DBD* labels tonic VO MNs (VO-MN15/16 and VO-MN15–17) as well as tonic VL MNs. White arrows indicate VO NMJs where large Dlg-positive boutons colocalize with presynaptic mCherry, consistent with tonic innervation. **(B)** *7-1^∪^FD4-Gal4,ChAT-Gal80* selectively labels tonic VO-MN15–17 but not VO-MN15/16. Dashed outlines indicate Dlg-positive VO NMJs lacking mCherry, corresponding to unlabeled VO-MN15/16 inputs. **(C)** *HB9-*Gal4 *(a.k.a VL-Gal4)* labels tonic VL MNs but no VO MNs. Arrowheads indicate VO NMJs lacking mCherry signal. **(D)** *27E09-Gal4 (a.k.a Is-Gal4)* labels the phasic MNISNb/d, which innervates VL muscles and VO15–16. Red arrows mark Dlg-positive VO17 NMJs lacking mCherry, consistent with absent MNISNb/d innervation. **(E–F)** *FD4^∪^27E09* **(E)** and *7-1^∪^27E09* **(F)** label both tonic VO-MN15–17 and phasic MNISNb/d, but not VO-MN15/16. **(G)** *HB9^∪^27E09* labels tonic VL MNs and MNISNb/d without labeling VO MNs. **(H)** *7-1^∪^FD4-Gal4, ChAT-Gal80* additionally labels tonic MNs innervating dorsal longitudinal muscles. Together, these data define VO motor innervation: convergent tonic input from VO-MN15/16 and VO-MN15–17, with additional phasic input from MNISNb/d to VO15–16.

FD4-Gal4, a driver characterized in this study, labeled a similar tonic MN population, including MN15–17 and DL-innervating tonic MNs, as well as a small number of non-MN cells **(Figure 4, Figure S5-S7)**. MultiColor FlpOut (MCFO) [31] analysis revealed that the main non-MN cells labeled by FD4-Gal4 were cholinergic sensory neurons, including chordotonal sensory neurons, which have been implicated in vibration sensing, and class IV multidendritic sensory neurons, which are involved in nociception **(Figure S7)**. Importantly, we did not observe class I multidendritic proprioceptors, the sensory-neuron class most strongly associated with larval crawling dynamics, in the FD4-Gal4 MCFO examples analyzed. FD4-Gal4 also labeled occasional unidentified interneurons.

NB7-1-Gal4 did not show the same sensory-neuron off-target profile as FD4-Gal4. Although occasional brain-lobe neurons were labeled by NB7-1-Gal4, FD4-Gal4 and NB7-1-Gal4 did not share the same non-MN expression pattern. Nevertheless, both lines labeled VO-MN15–17. Because NB7-1-Gal4 and FD4-Gal4 both provide access to MN15–17-containing tonic VO MN populations, we refer to these drivers collectively as VO-MN15–17-Gal4 when describing functional manipulations.

Previous work also suggested that VO15 and VO16, but not VO17, receive additional input from the phasic MN ISNb/d-Is, also known as RP5 [17]. Using 27E09-Gal4, which labels RP5 and the phasic MN ISN-Is/RP2 [26, 32], we confirmed that RP5 forms small phasic boutons on VO15 and VO16, but not VO17 **(Figure 4)**. Thus, VO muscles receive convergent tonic and phasic motor input: tonic input from MN15/16 and MN15–17, and phasic input from RP5 onto VO15 and VO16. In the following sections, we use these MN-Gal4 lines to perform gain- and loss-of-function manipulations to directly test the sufficiency and necessity of VO and VL muscle activity for protopodium dynamics.

### Contraction of VO muscles drives robust protopodium folding and ventral bending in larvae

We hypothesized that if VO muscles directly drive protopodium folding, optogenetic activation of VO MNs should be sufficient to induce protopodium folding. To test this, we optogenetically activated tonic VO-MN15–17 using Gal4-driven UAS-Chrimson-mCherry in larvae with a straight posture on a flat substrate. Light stimulation consistently induced robust ventral curvature, with larvae adopting a C-shaped posture in which VO muscles and protopodia were positioned on the concave side of the bend **(Figure 5A, Video S6)**. Because light exposure synchronously activates targeted MNs across multiple segments, this gain-of-function assay tests whether activation of a given MN population is sufficient to induce protopodium folding and body bending, rather than reproducing the sequential intersegmental motor pattern of crawling.

**Figure 5.**
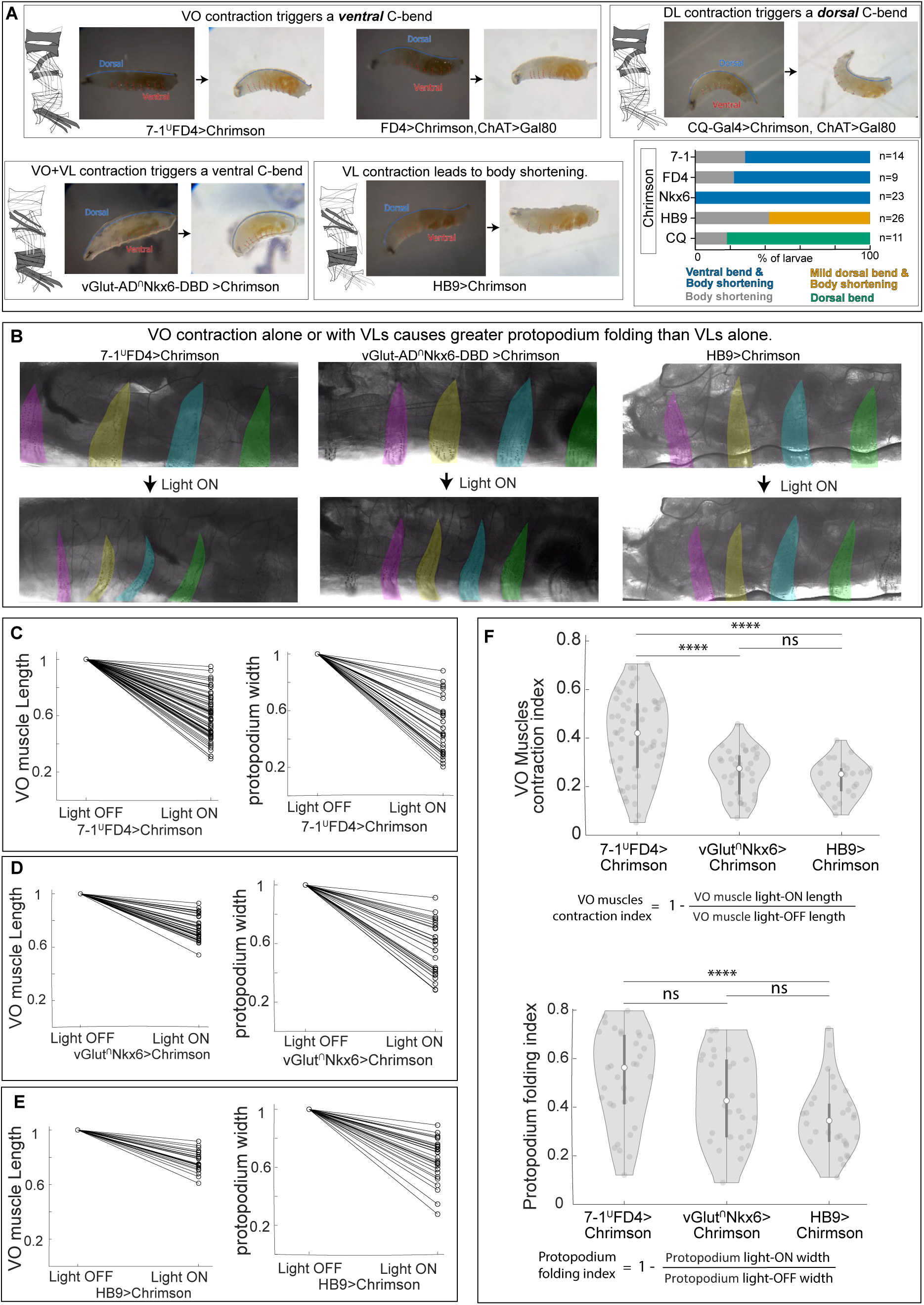
VO muscle activity is sufficient to drive protopodium folding. **(A)** Representative larval body postures following Chrimson-mediated optogenetic activation of distinct MN subsets. Dorsal trachea and ventral protopodia are indicated. Schematics show target muscle groups for each MN-Gal4 line. Activation of VO-MN15–17 using *NB7-1-Gal4* or *FD4-Gal4* induced a ventral C-bend accompanied by protopodium folding. *FD4-Gal4* activation in the presence of *ChAT-Gal80* produced a similar phenotype, indicating that the response is primarily driven by MN activation rather than cholinergic sensory-neuron activation. Co-activation of VO, VL, and DO MNs using *vGlut-AD∩Nkx6-DBD split-Gal4* also produced ventral bending and protopodium folding. In contrast, activation of dorsal MNs using *CQ-Gal4* induced a dorsal C-bend, while activation of VL/DO MNs using *HB9-Gal4* primarily caused body shortening. Stacked bar plots summarize postural responses across genotypes: *NB7-1-Gal4*, N = 14; *FD4-Gal4*, N = 9; *vGlut-AD∩Nkx6-DBD split-Gal4*, N = 23; *HB9-Gal4*, N = 26; *CQ-Gal4*, N = 11. **See Video S6**. **(B–F)** VO activation, either alone or together with VL/DO activation, drives greater protopodium folding than VL/DO activation alone. **(B)** High-resolution images of protopodia before and after optogenetic activation of VO-MN15–17 alone, VO + VL/DO MNs, or VL/DO MNs alone. **(C–E)** Quantification of VO muscle length and protopodium width following activation of VO-MN15–17 **(C)**, VO + VL/DO MNs **(D)**, or VL/DO MNs **(E)**. Values were normalized to the pre-stimulation frame. Because these experiments were performed under confocal illumination, the pre-stimulation frame is used as an operational baseline. **(F)** Violin plots comparing VO muscle contraction index and protopodium folding index across conditions. Indices were defined as 1 − (light ON/light OFF). Each point represents one VO muscle or protopodium. Apparent VO length changes in *HB9-Gal4>UAS-Chrimson* larvae likely reflect passive deformation associated with VL/DO-driven segment shortening rather than direct VO MN activation. Statistics: Kruskal– Wallis test with Dunn’s post hoc test; ns, not significant; ****, *P* < 0.0001. Sample sizes: *FD4-Gal4*, 13 trials from 8 animals; *HB9-Gal4*, 12 trials from 8 animals; *vGlut-AD∩Nkx6-DBD split-Gal4*, 17 trials from 7 animals. See **Video S7**.

Because FD4-Gal4 MCFO analysis revealed cholinergic sensory-neuron expression, we next tested whether the FD4-Gal4 gain-of-function phenotype required activation of these sensory neurons. To do this, we suppressed Gal4 expression in cholinergic sensory neurons using ChAT-Gal80. Optogenetic activation of FD4-Gal4 in the presence of ChAT-Gal80 still produced robust protopodium folding and ventral bending **(Figure 5A, Video S6)**. In addition, activation with NB7-1-Gal4, an independent VO-MN15–17 driver that does not share the sensory-neuron off-target profile of FD4-Gal4, produced a similar ventral C-bend and protopodium-folding phenotype **(Figure 5A, Video S6)**. Together, these results indicate that the FD4-Gal4 gain-of-function phenotype does not require activation of cholinergic sensory neurons and is most consistent with activation of the labeled VO-MN15–17 population.

We next asked whether activation of other ventral or dorsal muscle groups could produce a similar effect. Optogenetic activation using CQ-Gal4, a driver that targets dorsal longitudinal (DL) MNs and excludes VO MNs, failed to induce ventral protopodium folding and instead produced dorsal bending **(Figure 5A, Video S6)**. This comparison is particularly informative because the VO-MN15–17 drivers also label DL MNs. Thus, despite simultaneous activation of DL MNs, activation of VO-MN15–17 was sufficient to drive a ventral C-bend, whereas activation of DL MNs without VO-MN15–17 produced a dorsal C-bend.

To examine the contribution of VL and dorsal oblique (DO) muscles, we activated tonic MNs innervating these muscles using HB9-Gal4 [33]. In straight larvae, HB9-driven activation caused body shortening accompanied by mild dorsal curvature but did not produce pronounced ventral protopodium folding **(Figure 5A, Video S6)**. In contrast, simultaneous activation of VO, VL, and DO muscles using vGlut-AD∩Nkx6-DBD split-Gal4, which targets tonic VO MNs, including MN15/16 and MN15–17, together with VL MNs, resulted in ventral bending and protopodium folding comparable to activation of VO-MN15–17 alone **(Figure 5A, Video S6)**. Thus, across intact-larva gain-of-function experiments, the presence of VO MN activation was the condition most consistently associated with robust ventral bending and protopodium folding.

To further characterize the relationship between muscle contraction and protopodium folding, we combined optogenetic activation of selected MN populations with high-resolution confocal imaging of muscles and protopodia **(Figure 5B–F, Video S7)**. Activation of VO-MN15–17 produced VO muscle shortening accompanied by robust protopodium narrowing **(Figure 5B, C, F)**. Co-activation of VO, VL, and DO MNs also induced protopodium narrowing **(Figure 5B, D, F)**. By contrast, activation of VL/DO MNs with HB9-Gal4 produced a smaller reduction in protopodium width and only modest apparent changes in VO muscle length **(Figure 5B, E, F)**. Because HB9-Gal4 does not label VO MNs **(Figure 4)**, the apparent VO length change in this condition likely reflects passive deformation associated with VL/DO-driven segment shortening rather than direct VO MN activation.

Together, these experiments demonstrate that VO muscle activity is sufficient to drive robust protopodium folding and ventral bending. VL/DO activation can alter body shape and produce modest protopodium narrowing, but under these optogenetic activation conditions, it is less effective than VO activation in driving protopodium folding **(Figure 5F)**.

### VO muscles are necessary for protopodium folding and contribute to protopodium stride during crawling

To determine which muscle groups are required for efficient crawling and protopodium dynamics, we performed MN loss-of-function experiments targeting different combinations of MNs innervating VO and VL muscles. Using MN-Gal4 drivers **(Figure 6A)** to express UAS-GtACR1, we silenced VO and/or VL MN activity while imaging larval locomotion under blue-light illumination. FD4-Gal4 and NB7-1-Gal4 target one tonic VO MN class, VO-MN15–17, while leaving the second tonic VO MN class, MN15/16, intact. HB9-Gal4 targets all tonic VL MNs; vGlut-AD∩Nkx6-DBD targets tonic VO and VL MNs; and 27E09-Gal4 targets phasic Is MN inputs, including phasic inputs to VO and VL muscles. These driver combinations allowed us to compare silencing of phasic MNs alone, tonic VL MNs, one tonic VO MN class, tonic VO and VL MNs together, and tonic plus phasic MN inputs.

**Figure 6.**
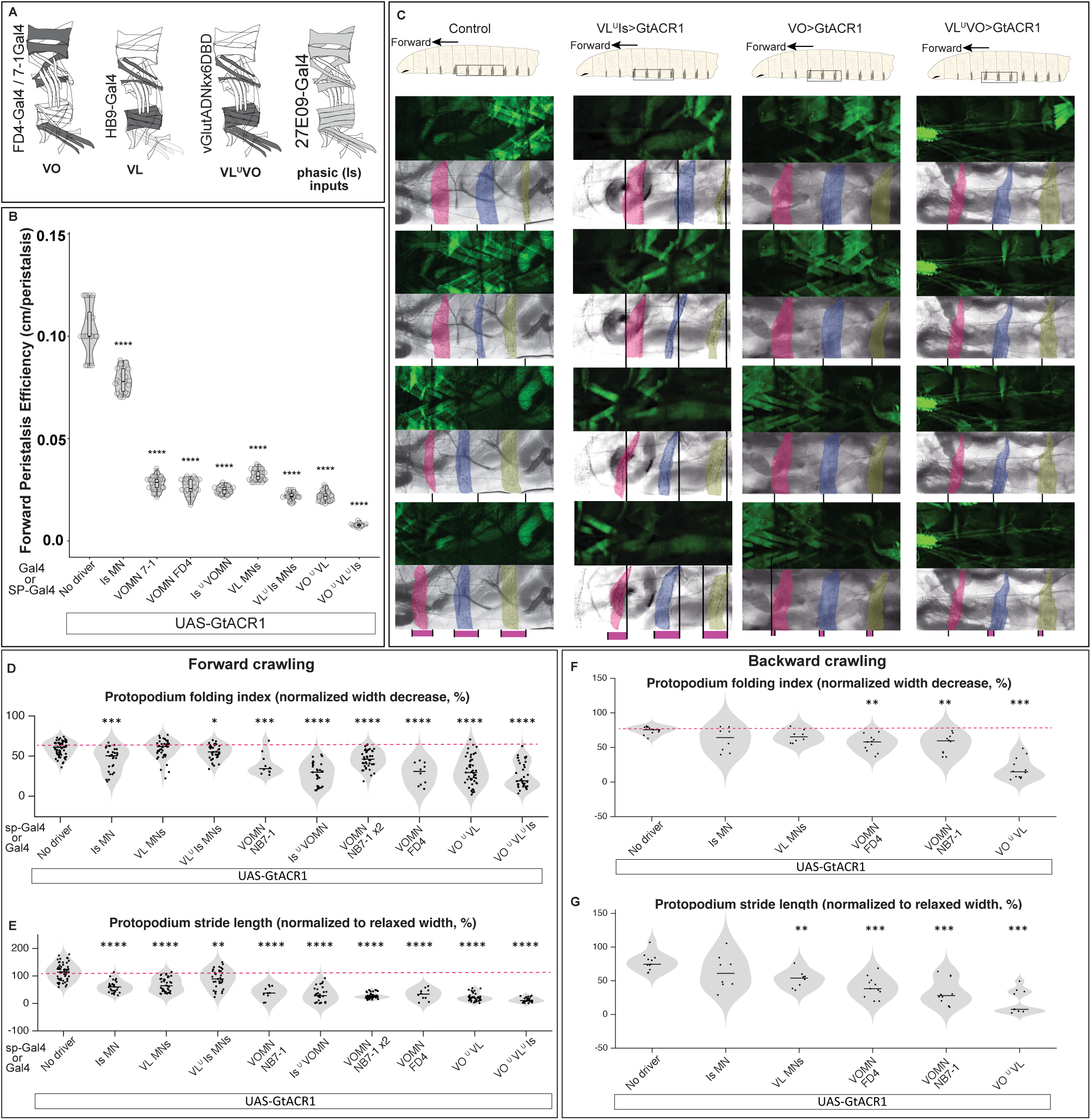
VO muscles are necessary for protopodium folding and contribute to protopodium stride during crawling. **(A)** Schematic of MN-Gal4 and split-Gal4 drivers used for GtACR1-mediated silencing. *FD4-Gal4* and *NB7-1-Gal4* drive silencing in one tonic VO MN class, VO-MN15–17, but not in MN15/16. *HB9-Gal4* targets tonic VL MNs; *vGlut-AD∩Nkx6-DBD* targets tonic VO and VL MNs; and *27E09-Gal4* targets phasic Is MN inputs. Light gray indicates phasic MN inputs, and dark gray indicates tonic MN inputs. **(B)** GtACR1-mediated silencing of MN subsets reduces forward crawling efficiency, defined as distance traveled per peristaltic cycle. Silencing phasic Is MNs modestly reduced efficiency, whereas tonic MN silencing caused stronger impairments. Silencing tonic VL MNs with *HB9-Gal4* reduced efficiency, consistent with a role for VL muscles in locomotor displacement. Silencing one tonic VO MN class also reduced efficiency, and co-silencing tonic VO/VL or tonic plus phasic inputs caused stronger defects. Violin plots show peristalsis efficiency (N=10–22 larvae per genotype; each dot represents one animal). Values are normalized to the control median for visualization only. See Video S8. **(C)** Time-lapse images of protopodium movement and muscle activity during forward crawling. GCaMP6f reports muscle activity. Color-matched protopodia indicate the same segment across frames; magenta bars denote stride length. See **Video S9**. **(D,E)** Forward protopodium dynamics. **(D)** Protopodium folding index. Silencing one tonic VO MN class strongly reduced folding, whereas tonic VL silencing had weaker effects. **(E)** Protopodium stride length. Silencing either VO or VL MNs reduced stride length, and combined silencing produced stronger defects. “NB7-1 x2” denotes larvae carrying two copies of *NB7-1-Gal4* to increase Gal4-driven GtACR1 expression **(F,G)** Backward protopodium dynamics. **(F)** Silencing one tonic VO MN class reduced folding despite continued MN15/16 activity, whereas tonic VL silencing caused a weaker folding defect; combined VO/VL silencing further reduced folding. **(G)** Silencing either VO or VL MNs reduced stride length, with combined VO/VL silencing producing the strongest defect. Statistics: Kruskal– Wallis tests followed by planned Wilcoxon rank-sum comparisons with Holm–Bonferroni correction. Asterisks indicate *P* < 0.05, 0.01, 0.001, and 0.0001. Sample sizes in D–G: N = 3–6 larvae per genotype; 1–3 protopodia per larva.

Because FD4-Gal4 labels a small number of cholinergic sensory neurons, we considered whether sensory-neuron silencing could contribute to the FD4-Gal4 loss-of-function phenotype. However, previous work showed that inhibiting class II, class III, class IV, or chordotonal sensory neurons does not substantially disrupt larval crawling [34]. In addition, NB7-1-Gal4 does not share the same sensory-neuron off-target profile as FD4-Gal4, yet FD4-Gal4 and NB7-1-Gal4 produced similar defects in protopodium folding and crawling when used to silence VO-MN15–17. These observations argue that the shared loss-of-function phenotypes observed with FD4-Gal4 and NB7-1-Gal4 primarily reflect perturbation of their common VO-MN15–17 population rather than line-specific sensory-neuron off-targets.

As an initial measure of locomotor output, we quantified crawling efficiency [35, 36], defined as the distance traveled per forward peristaltic cycle, in L3 larvae crawling within a straight groove on an agarose substrate. Acute silencing of VO and/or VL MN inputs significantly reduced crawling efficiency **(Figure 6B, Video S8)**, consistent with effective GtACR1-mediated perturbation and indicating that these motor outputs contribute to forward locomotion. However, reduced crawling efficiency could arise from distinct mechanisms. For example, silencing VL MNs may disrupt visceral pistoning and intersegmental body-mass transport [37] without directly impairing protopodium folding, whereas silencing VO MNs is expected to compromise protopodium folding itself. We therefore directly examined protopodium dynamics using high-resolution imaging during forward crawling.

We assessed two protopodium-specific parameters during forward crawling: protopodium folding index and protopodium stride length **(Figure 6C–E, Video S9)**. The folding index measures the extent of protopodium deformation during the fold-and-lift phase, whereas stride length measures the distance moved by an individual protopodium during a crawl cycle. In control larvae, protopodia folded robustly and moved anteriorly during each forward peristaltic cycle. Silencing phasic Is MN inputs reduced protopodium stride length but had a comparatively modest effect on folding **(Figure 6D–E)**. Silencing tonic VL MNs also reduced protopodium stride length, but did not significantly compromise protopodium folding **(Figure 6D–E)**. This pattern is consistent with VL muscles contributing to overall locomotor displacement and body-wall mechanics, while having a less direct role in protopodium folding.

In contrast, silencing one tonic VO MN class produced strong defects in both protopodium folding and stride length. Silencing VO-MN15–17 using FD4-Gal4 or NB7-1-Gal4, while leaving MN15/16 intact, significantly reduced the folding index and shortened protopodium stride length during forward crawling **(Figure 6D–E)**. These defects were further exacerbated when phasic MN inputs were co-silenced. Combined silencing of tonic VO and VL MNs also reduced both folding and stride length, and simultaneous silencing of tonic VO and VL MNs along with phasic inputs produced the strongest impairment in forward protopodium movement **(Figure 6D–E)**. Thus, during forward crawling, VO MN activity is particularly important for protopodium folding, whereas both VO and VL MN activity contribute to the full stride of protopodium movement.

During backward crawling, we again quantified both protopodium folding index and protopodium stride length **(Figure 6F–G)**. Silencing one tonic VO MN class reduced protopodium folding during backward crawling, whereas tonic VL MN silencing did not substantially reduce folding **(Figure 6F)**. Similar to forward crawling, silencing either VO-MN15–17 or tonic VL MNs reduced backward protopodium stride length **(Figure 6G)**. Combined VO/VL MN silencing produced the strongest impairment in backward stride length and also reduced folding **(Figure 6F–G)**. Thus, the backward-crawling data are consistent with the forward-crawling results: VO MN activity is particularly important for protopodium folding, while both VO and VL MN activity contribute to protopodium stride length.

Together, these results demonstrate that VO muscle activity is required for normal protopodium folding during both forward and backward crawling, and that VO and VL MN activity both contribute to protopodium stride length. Importantly, loss of activity in a single tonic VO MN class is sufficient to disrupt protopodium folding and reduce stride length, even though the second tonic VO MN class remains intact; by contrast, HB9-Gal4 targets a broader tonic VL motor pool, yet VL silencing does not significantly compromise protopodium folding. This difference supports the conclusion that VO activity has a particularly important role in protopodium folding. Moreover, by independently manipulating tonic and phasic MN inputs, these experiments reveal distinct contributions of tonic and phasic motor pathways to crawling efficiency, protopodium folding, and protopodium stride dynamics.

### VO MNs receive excitatory and inhibitory inputs from an intersegmentally connected PMN network

The behavior-specific timing of VO muscle activity prompted us to investigate the PMN circuits that drive VO MNs. Building on our previously published PMN–VO MN connectome in segment A1 of the larval VNC [1], we extended the reconstruction to include VO MNs and their premotor inputs in segments A2 and A3. Analysis across six hemisegments **(A1–A3)** identified A06c and A27h as major bilaterally connected sources of synaptic input to tonic VO MNs **(Figure 7A,B, Figure S8–S11, Supplementary Table 1)**. Across segments A1–A3, the combined bilateral A06c contribution accounts for approximately 17.6%, 18.0%, and 23.2% of total synaptic input to VO MNs, respectively, whereas the combined bilateral A27h contribution accounts for approximately 10.6%, 10.6%, and 13.5% **(Supplementary Table 1)**. A06c is a GABAergic inhibitory PMN whose output is strongly enriched on MN15/16 and MN15–17 **(Figure S9, Supplementary Tables 2 and 3)**, whereas A27h is a cholinergic excitatory PMN [4] that forms numerous synapses onto VO MNs and fewer synapses onto other MN classes **(Figure 7B)**. Notably, both A06c and A27h are bilaterally connected, such that a PMN in one hemisegment synapses onto VO MNs in both left and right hemisegments **(Figure 7B, Figure S10, Supplementary Tables 2 and 3)**. In addition, A27h establishes direct synapses onto A06c within the same segment, forming an incoherent type-1 feedforward loop **(Figure 7A)**.

**Figure 7.**
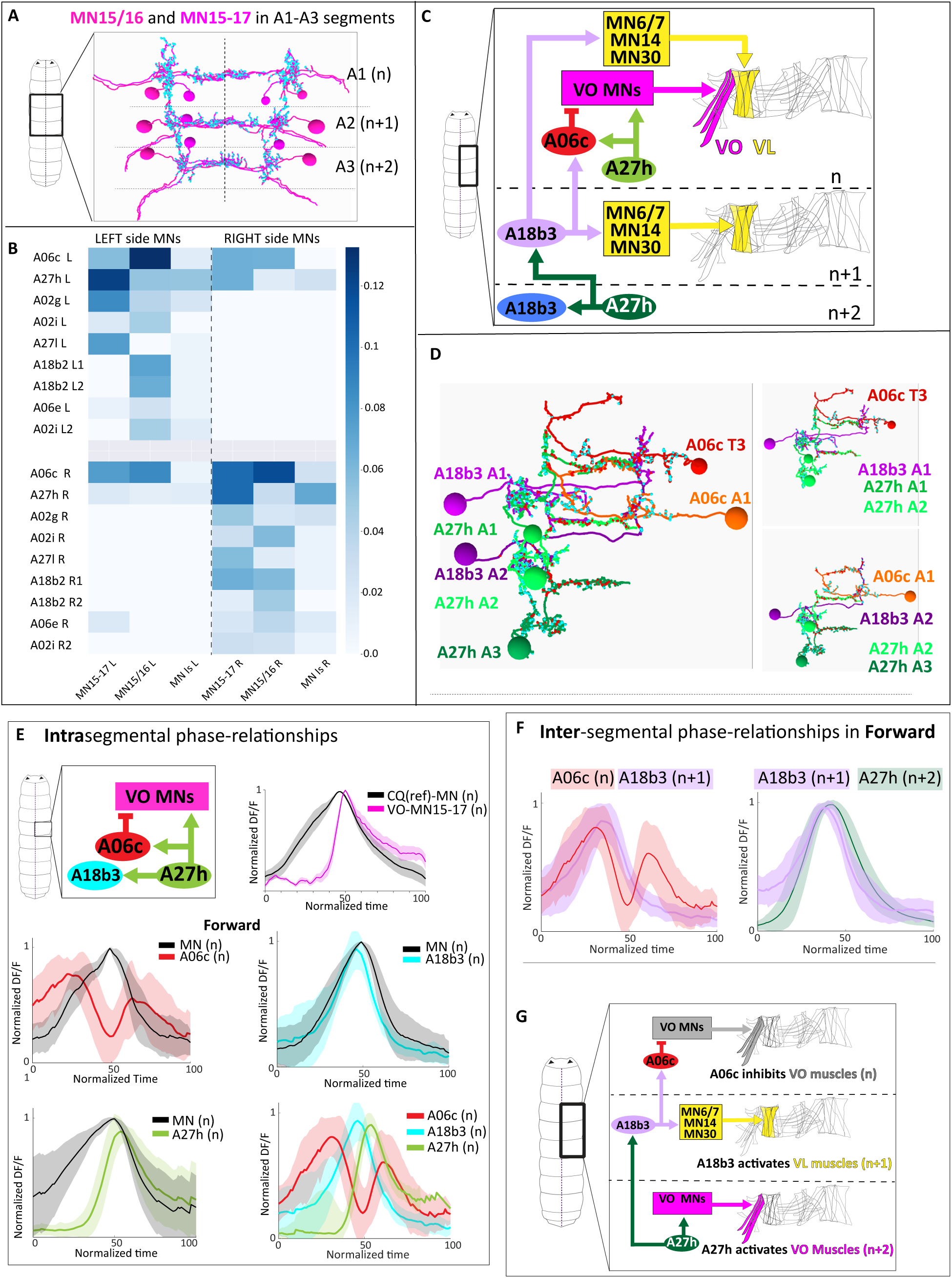
The coordinated activity of A06c, A18b3, and A27h PMNs defines the activity window for VO MNs during forward crawling. **(A)** EM reconstructions of tonic VO MNs, including MN15/16 and MN15–17, across segments A1–A3. VO MNs receive premotor input from the excitatory PMN A27h and the inhibitory PMN A06c. **(B)** Heatmap showing synaptic connectivity between PMNs and VO MNs. PMNs are shown as rows and VO MNs as columns, divided by left and right hemisegments. A06c and A27h are among the major premotor inputs to VO MNs. **(C)** Schematic of the intersegmental A27h–A18b3–A06c motif. A18b3 in segment **n+1** receives input from A27h in segments n+1 and n+2, connects to A06c in segment **n**, and contacts VL MNs in segments n and n+1. A27h in segment n+2 also connects to VO MNs in its own segment. **(D)** TEM reconstructions of A06c, A18b3, and A27h neurons across consecutive segments. Red and cyan dots indicate pre- and postsynaptic sites, respectively. **(E)** Intrasegmental phase relationships between CQ-labeled reference MNs, VO MNs, and PMNs during fictive forward locomotion. *CQ-LexA* was used to express *GCaMP6m* in reference MNs, and *FD4-Gal4* or PMN-Gal4/split-Gal4 drivers were used to express *UAS-jRCaMP1b* in VO MNs or PMNs. VO MNs activate later than CQ-labeled reference MNs. A06c exhibits two peaks of activity, one before and one after reference MN activation. A27h peaks between the two A06c peaks, whereas A18b3 is synchronous with early-firing reference MNs. **(F)** Intersegmental phase relationships during fictive forward locomotion. A18b3 in segment **n+1** is active in phase with the first peak of A06c in segment **n**, and A27h in segment **n+2** is active in phase with A18b3 in segment **n+1**. **(G)** Model summarizing how the A27h–A18b3–A06c motif may coordinate VO and VL MN/muscle timing across segments during forward crawling. Calcium activity was aligned to a normalized 0–100% fictive locomotor cycle. Backward PMN activity patterns are shown in **Figure S12C**.

Several additional PMNs, including A18b2/Ghost, A02i, and A02g, also synapse onto VO MNs **(Figure 7B)**. However, compared with A06c and A27h, these neurons form fewer synapses onto VO MNs and exhibit unilateral connectivity, contacting only one side of each bilateral MN pair **(Figure 7B, Figure S10, Supplementary Table 1)**. Thus, the connectome identifies A06c and A27h as prominent bilateral PMN inputs to tonic VO MNs, while also revealing additional unilateral PMNs that likely contribute to VO MN regulation.

Extending the connectome analysis to adjacent segments revealed that A06c receives input from previously reported excitatory neurons, including Canon/A18g [11] and A18b3 [1, 38], in posterior segments. Importantly, we identified a tri-segmental circuit motif involving A18b3 and two major PMNs upstream of VO MNs, A27h and A06c **(Figure 7C,D, Supplementary Table 4)**. In this motif, A18b3 in segment **n+1** receives excitatory input from A27h in segments **n+1** and **n+2** and synapses onto inhibitory A06c in the next anterior segment, **n (Figure 7C,D)**. A18b3 also synapses onto tonic VL MNs **(MN6/7, MN14, and MN30)** in its own segment and in the next anterior segment **(Figure 7C)**. Morphologically, A18b3 corresponds to the previously described forward-dedicated cholinergic lateral interneuron 1 (CLI1) [38] **(Figure S11)**.

Together, these anatomical data identify intrasegmental **A27h–A06c** and intersegmental **A27h– A18b3–A06c** premotor motifs as candidate circuits for shaping VO MN activity during larval locomotion. In the following sections, we test the functional roles of these PMNs during movement, with particular emphasis on the **A27h–A18b3–A06c** motif and its role in forward crawling.

### The coordinated activity of A06c, A18b3, and A27h PMNs defines the activity window for VO MNs during forward crawling

To begin to understand how the A27h–A18b3–A06c motif may regulate VO MN activity, we performed calcium imaging in isolated larval CNS preparations. During fictive locomotion in isolated CNS preparations, spontaneous motor activity propagates from posterior to anterior segments during forward locomotion and from anterior to posterior segments during backward locomotion. We first asked whether VO and VL MNs exhibit behavior-specific activity patterns consistent with the VO/VL muscle calcium-imaging and muscle-length measurements shown in **Figures 1 and 2**.

We expressed UAS-jRCaMP1b in VO and VL MNs using the vGlut-AD∩Nkx6-DBD split-Gal4 driver. Using EM-reconstructed MN morphology as a guide, we identified dendritic regions corresponding to VO and VL MNs, allowing calcium signals to be measured selectively from VO, VL, or combined VO/VL dendritic regions. Consistent with the muscle calcium-imaging and muscle-length data shown in **Figures 1 and 2**, VO MNs activated later than VL MNs during fictive forward locomotion. In contrast, VO and VL MNs showed more synchronous activity during fictive backward locomotion **(Figure S12A,B)**.

We next examined the phase relationship between VO MNs and DL MNs using dual-color calcium imaging. We used CQ-LexA to express GCaMP6m in DL MNs, which include a subset of type Ib MNs 2, 3, 4, 9, and 10 and are hereafter referred to as reference MNs. In parallel, we used FD4-Gal4 to express jRCaMP1b in VO MNs. These experiments confirmed that VO MNs activate later than the CQ-labeled reference MNs during fictive forward locomotion **(Figure 7E)**. Together, these MN imaging experiments show that the behavior-specific VO/VL timing observed in intact crawling larvae is also reflected in fictive locomotor activity in isolated CNS preparations.

Having established these MN phase relationships, we next examined how A06c, A18b3, and A27h PMNs are recruited during fictive forward and backward locomotion. We expressed GCaMP6m in reference MNs using CQ-LexA and expressed jRCaMP1b in A06c, A27h, or A18b3 PMNs using Gal4 or split-Gal4 drivers. Calcium activity was recorded from isolated larval CNS preparations during spontaneous fictive locomotor waves.

During fictive forward locomotion, A06c PMNs in a given segment exhibited two distinct peaks of activity: one preceding and one following peak activity of the reference MNs in the same segment **(Figure 7E, Video S10)**. In contrast, during fictive backward locomotion, A06c activity was restricted to a single peak occurring after reference MN activation **(Figure S12C, Video S10)**. The absence of the early inhibitory peak during backward locomotion suggests that VO MNs do not receive the same early A06c-mediated inhibition in this behavior, consistent with earlier activation of VO muscles during backward crawling. Based on these behavior-specific activity patterns, we refer to A06c as a bi-pattern PMN.

We next examined the excitatory PMN A27h. Consistent with previous reports [1, 4, 39], A27h was inactive during backward locomotion **(Figure S12C, Video S10)**, indicating that it does not provide excitatory drive to VO MNs during this behavior. In contrast, during forward locomotion, A27h reached peak activity between the first and second peaks of A06c, coinciding with the VO MN activity window **(Figure 7E, Figure S12, Video S10)**. We therefore refer to A27h as a forward-dedicated PMN.

We then analyzed the activity of A18b3 relative to A06c, A27h, and the reference MNs. Consistent with prior work [38], A18b3 was selectively active during forward locomotion and was synchronous with early-firing MNs **(Figure 7E, Figure S12, Video S10)**. Based on EM connectomics, A18b3 in segment **n+1** synapses onto A06c PMNs in anterior segments **n** and **n−1**, while receiving input from A27h in its own and posterior segments **n+1** and **n+2 (Figure 7C,D)**. To determine how this multi-segmental architecture operates during forward crawling, we examined intersegmental phase relationships using new dual-color calcium imaging of A18b3 and A06c, together with reanalysis of the datasets used in **Figure 7F**.

During fictive forward locomotion, A27h in segment n+2 and A18b3 in segment n+1 were active in phase with the first peak of A06c in segment n **(Figure 7F,G)**. Together with the connectome, this timing is consistent with a sequence in which A27h is positioned to activate A18b3, which in turn contributes to activation of A06c in the next anterior segment. While A27h in segment n+2 provides excitatory input to late-firing VO MNs in its own segment, it also engages A18b3 in segment n+1, which is positioned to excite VL MNs, including MN6/7, MN14, and MN30, and promote early inhibition of VO MNs in the next anterior segment via A06c **(Figure 7F,G)**.

The second peak of A06c activity may be influenced by intrasegmental excitation from A27h through the A27h→A06c feedforward motif **(Figure 7E)**. A06c likely integrates A27h-derived excitation with additional excitatory and inhibitory inputs, which could shape the timing of the second A06c peak. This interpretation is consistent with previous work [5] showing that phase delays between larval motor pools can arise from the balance of premotor excitation and inhibition rather than from delayed excitation alone.

Together, these data indicate that the **A27h–A18b3–A06c** intersegmental motif defines a temporally constrained window of VO MN activation during forward crawling.

### Connectome-based monosynaptic connections within the A27h–A18b3–A06c motif are functional in the isolated larval CNS

To determine whether the synaptic connections identified in the EM connectome are functional, we combined optogenetic activation with calcium imaging in isolated larval CNS preparations **(Figure 8A–D)**. To test the A18b3→A06c connection, we expressed Chrimson in A18b3 and GCaMP6m in A06c. Optogenetic activation of A18b3 elicited a robust increase in A06c calcium signal, indicating functional connectivity between these neurons **(Figure 8A,A′)**. This response persisted in the presence of tetrodotoxin (TTX), supporting a monosynaptic connection.

**Figure 8.**
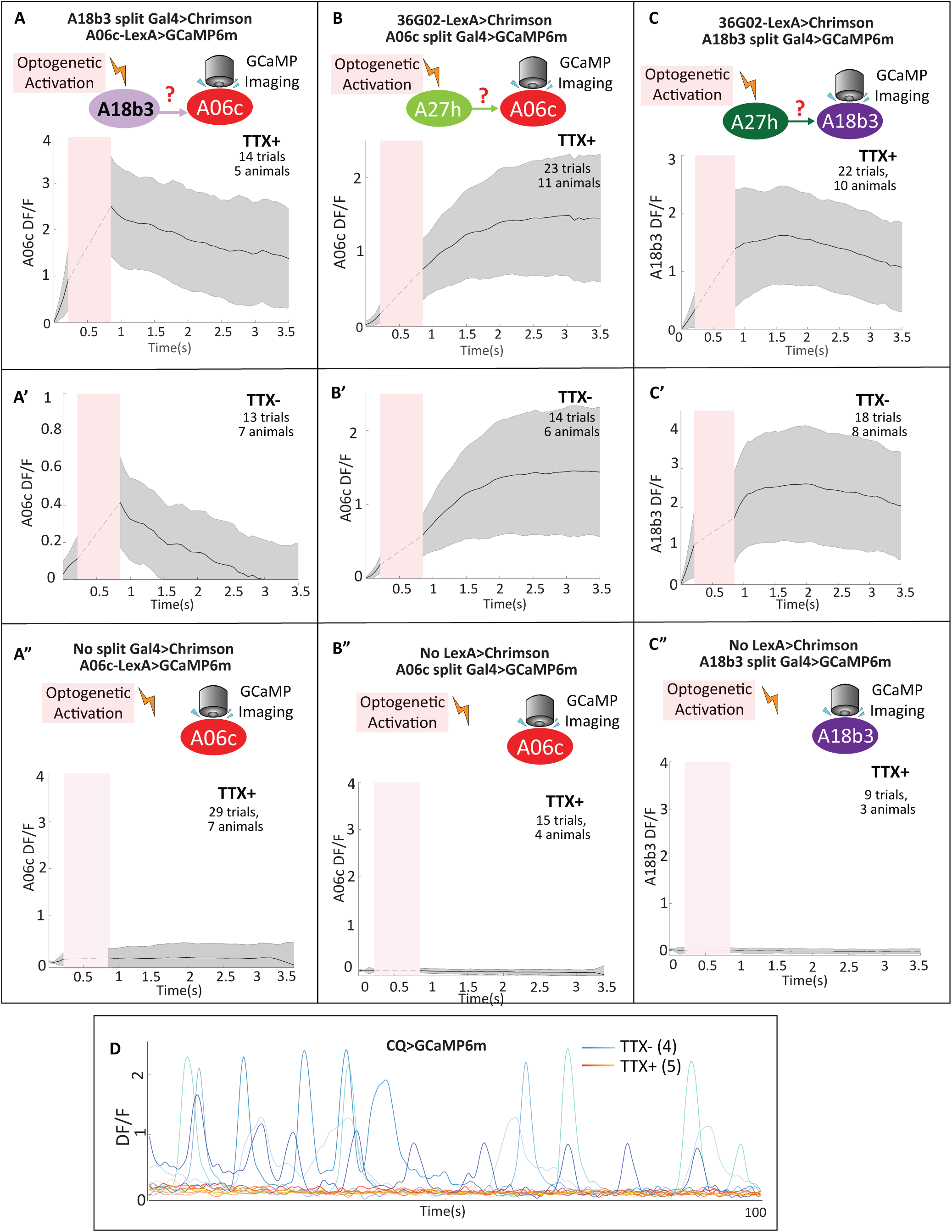
Functional validation of connectome-predicted monosynaptic connections within the A27h–A18b3–A06c premotor motif. **(A)** Optogenetic activation of A18b3 (*A18b3 split-Gal4>UAS-Chrimson*) while imaging calcium activity in A06c (*A06c-LexA>GCaMP6m*). **(A,A′)** A18b3 activation induces a robust increase in A06c ΔF/F both in the absence (TTX−) and presence (TTX+) of tetrodotoxin, consistent with a functional monosynaptic connection. **(A″)** Control animals lacking the A18b3 split-Gal4 driver show little or no A06c calcium response. **(B)** Optogenetic activation of A27h (*36G02-LexA>LexAop-Chrimson*) while imaging A06c (*A06c split-Gal4>GCaMP6m*). **(B,B′)** A27h activation elicits a strong A06c calcium response that persists under TTX, indicating monosynaptic connectivity. **(B″)** No response is observed in control animals lacking the 36G02-LexA driver. **(C)** Optogenetic activation of A27h (*36G02-LexA>LexAop-Chrimson*) while imaging A18b3 (*A18b3 split-Gal4>GCaMP6m*). **(C,C′)** A27h activation produces a calcium response in A18b3 under both TTX− and TTX+ conditions. **(C″)** Control animals lacking the 36G02-LexA driver show no detectable response. Across calcium-response traces, solid lines indicate the mean, shaded regions indicate mean ± s.d., and pink bars denote the period of optogenetic stimulation. Total numbers of larvae and trials are indicated in each panel. **(D)** Tetrodotoxin suppresses spontaneous network activity in isolated larval CNS preparations. Representative GCaMP6m traces from CQ MNs show robust spontaneous activity in the absence of TTX (blue), which is abolished following TTX application (orange), confirming effective blockade of polysynaptic and action potential-dependent network activity.

We next tested the predicted excitatory connections from A27h to A06c and A18b3. In the initial experiments, we expressed Chrimson in A27h using 36G02-LexA and recorded calcium activity in either A06c or A18b3. Optogenetic activation of A27h induced calcium responses in both downstream neurons **(Figure 8B,C)**, and these responses were preserved in the presence of TTX, consistent with monosynaptic connectivity **(Figure 8B′,C′)**. To address potential off-target concerns associated with 36G02-LexA, we performed an additional validation experiment using the more specific A27h split-Gal4 line to drive Chrimson while recording calcium responses in A06c. Activation of A27h using this split-Gal4 line also induced a calcium increase in A06c **(Figure S13A)**, consistent with the 36G02-LexA result and supporting the functional A27h→A06c connection with an independent A27h driver.

To further assess the specificity of the 36G02-LexA optogenetic experiments, we performed negative-control experiments using neurons that are not predicted to be monosynaptically downstream of A27h. Based on previous studies, Ifb-Bwd and Canon/A18g are recruited during backward, but not forward, crawling [9, 11]. Furthermore, according to the EM connectome, A27h is not monosynaptically upstream of either Ifb-Bwd or Canon/A18g. We therefore reasoned that, in the presence of TTX, optogenetic activation of A27h with 36G02-LexA should not evoke calcium responses in Ifb-Bwd or Canon/A18g. As predicted, 36G02-LexA-mediated activation did not induce detectable GCaMP responses in either Ifb-Bwd or Canon/A18g **(Figure S13C,D)**. These negative controls support the specificity of the A27h optogenetic-validation experiments.

In addition, because previous connectome analysis [11] identified Canon/A18g as a candidate backward-related input to A06c, we tested whether the predicted Canon/A18g→A06c connection is functional. Using a previously published Canon/A18g split-Gal4 line to drive Chrimson while recording calcium activity in A06c, we found that optogenetic activation of Canon/A18g induced a calcium increase in A06c **(Figure S13E)**. This result validates the functional connectivity of the Canon/A18g→A06c pathway and identifies a candidate route through which backward-related circuitry could recruit A06c.

To exclude nonspecific effects of optogenetic stimulation or imaging [40], we also performed matched genetic-control experiments lacking the relevant Gal4 or LexA driver for Chrimson expression. In these control conditions, optogenetic stimulation failed to elicit calcium responses in the recorded downstream neurons **(Figure 8A″–C″; Figure S13B)**. These controls indicate that the observed responses require specific driver-dependent Chrimson expression and are unlikely to be caused by light artifacts or nonspecific network activity.

Together, these results demonstrate that the A27h→A06c, A27h→A18b3, and A18b3→A06c connections identified in the EM connectome are functional, monosynaptic, and genetically specific in isolated larval CNS preparations. They also validate the backward-related Canon/A18g→A06c connection and show that A27h activation does not nonspecifically recruit unrelated backward-active neurons that are not monosynaptically downstream of A27h. These data provide a validated circuit framework for interpreting premotor control of VO MN activity relevant to forward crawling and for understanding how distinct upstream inputs may recruit the shared inhibitory PMN A06c during different locomotor behaviors.

### Disrupting the A27h–A18b3–A06c motif compromises intersegmental and intrasegmental coordination of VO and VL muscles during forward locomotion

To determine how the A27h–A18b3–A06c motif regulates crawling, we silenced these PMNs individually or in dual combinations and examined muscle activity patterns using calcium imaging in intact behaving larvae. Our EM connectome and calcium-imaging data suggest that coordinated activity of A27h in segment n+2, A18b3 in segment n+1, and A27h in segment n generates two sequential activity peaks in A06c in segment n, thereby defining a temporal window for VO MN activation during forward locomotion **(Figure 9A)**. A simple prediction from this model is that silencing A06c or its upstream neurons, A18b3 and A27h, would compromise this temporal window and cause premature activation of VO muscles during forward crawling.

**Figure 9.**
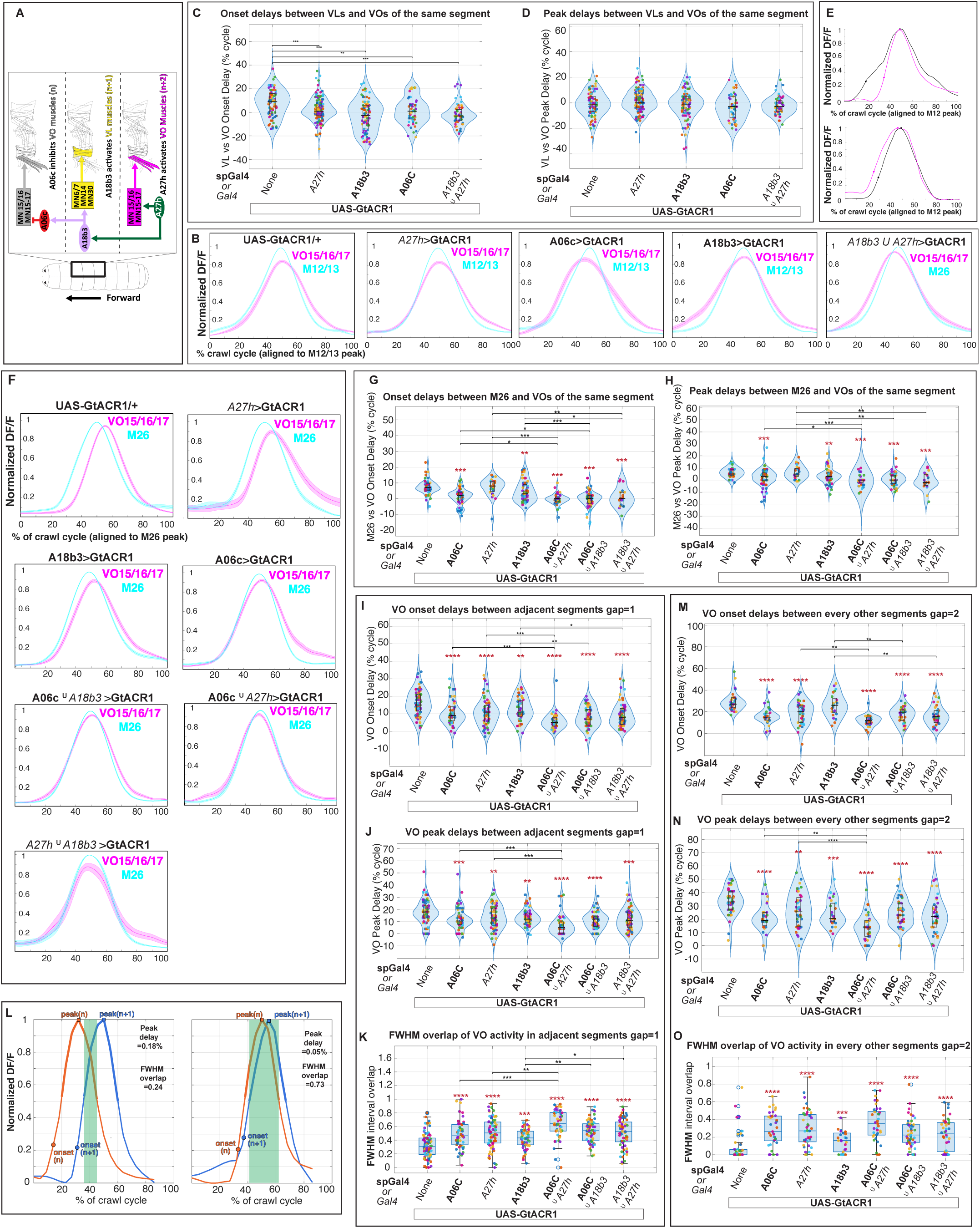
Disruption of the A27h–A18b3–A06c motif alters intrasegmental VO/VL timing and increases intersegmental VO synchrony during forward locomotion. **(A)** The tri-segmental premotor motif. **(B–E)** Intrasegmental timing of pooled VO15/16/17 activity relative to pooled VL M12/13 activity from lateral-view recordings. Disruption of the A27h– A18b3–A06c motif shifts the onset of VO activity earlier relative to VL activity. **(B)** Cycle-normalized calcium profiles aligned to the M12/13 peak. **(C,D)** Onset delay **(C)** and peak delay **(D)** between VL M12/13 and VO15/16/17 within the same segment. Positive values indicate that VO activity occurs after VL activity. **(E)** Representative normalized calcium traces of VO15/16/17 and M12/13 across genotypes. **(F–H)** Intrasegmental VO timing relative to M26. PMN perturbations make VO activity more synchronous with the earlier-activated reference muscle M26. **(F)** Cycle-normalized calcium profiles of pooled VO15/16/17 and M26 aligned to the M26 peak. **(G,H)** Onset delay **(G)** and peak delay **(H)** between M26 and VO15/16/17. Positive values indicate that VO activity occurs after M26. **(I–O)** Intersegmental VO15/16/17 coordination. Silencing A06c, A27h, or A18b3 reduces intersegmental delays and increases overlap of VO activity across adjacent and non-adjacent segments. **(I–K)** Onset delay **(I)**, peak delay **(J)**, and FWHM interval overlap **(K)** between adjacent VO segments, gap = 1. **(L)** Representative examples of temporally segregated and highly overlapping VO activity between adjacent segments; green shading marks FWHM overlap. **(M–O)** Onset delay **(M)**, peak delay **(N)**, and FWHM interval overlap **(O)** between every other VO segment, gap = 2. Solid lines indicate means; shaded regions indicate ± SEM. Each dot represents one crawl-bout measurement pooled across larvae. For **B–H**, N = 5–15 larvae; n = 19–80 measurements. For **I–K** and **M–O**, N = 5–15 larvae; n = 12–96 measurements. Statistics: Kruskal–Wallis tests followed by planned Wilcoxon rank-sum comparisons with Holm–Bonferroni correction. Brown asterisks indicate differences from control; black brackets indicate significant pairwise genotype differences. Only significant comparisons are shown. Italic PMN labels indicate conventional Gal4 lines; bold PMN labels in the figure indicate split-Gal4 lines. The union symbol (∪) denotes combined PMN silencing.

However, A27h and A18b3 are not expected to act solely through A06c. A27h synapses onto A06c and VO MNs, but also contacts additional PMNs. Similarly, A18b3 synapses onto A06c and a few other PMNs in the next anterior hemisegment and also contacts VL MNs. Thus, loss of A18b3 or A27h function could mimic aspects of A06c loss of function, but may also produce additional muscle-recruitment defects that cannot be explained solely by altered A06c activity. Based on these predictions, and because the A27h–A18b3–A06c motif spans three consecutive segments, we hypothesized that disrupting this circuit would impair both intersegmental and intrasegmental coordination of VO and VL muscle activity.

To test this, we silenced A06c, A27h, or A18b3 PMNs individually or in combination and examined VO muscle activation timing during forward crawling. PMNs were silenced using A06c split-Gal4, 36G02-Gal4, 47E12-Gal4, A18b3 split-Gal4, or dual-driver combinations to drive UAS-GtACR1 expression **(Figure 9; Videos S11–S12)**. Muscle activity was monitored using muscle-specific expression of GCaMP6f. Because some dual-silencing experiments required combining two split-Gal4 lines, we examined the expression patterns of the corresponding dual split-Gal4 combinations **(Figure S11B,C)**. The *A18b3 split-Gal4 ∪ A06c split-Gal4>UAS-GtACR1-EGFP* combination labeled the expected segmentally repeated PMN populations in the VNC, with only occasional non-repetitive off-target cells. The *A27h split-Gal4 ∪ A06c split-Gal4>UAS-GtACR1-EGFP* combination also labeled the expected A27h and A06c PMNs in the VNC, although additional labeling was observed in the brain lobes **(Figure S11B,C)**. Importantly, we did not observe obvious additional segmentally repeated off-target neurons in the VNC for either dual split-Gal4 combination.

We first assessed whether silencing the A27h–A18b3–A06c motif alters the intrasegmental phase relationship between VO15/16/17 muscles and VL muscle groups. Because this analysis required simultaneous visualization of VO muscles and muscles spanning the dorsoventral axis, intact larvae were imaged from a lateral view. Consistent with the model presented in **Figure 2**, VL muscles in control larvae began to show activity, measured by GCaMP6f signal rise, before VO muscles. In contrast, disrupting components of the A27h–A18b3–A06c motif shifted VO activity earlier. In larvae with A27h loss of function, VO activity became more synchronous with VL activity. In larvae with individual A18b3 or A06c loss of function, as well as dual A18b3/A27h silencing, VO activity occurred even earlier relative to VL activity **(Figure 9B–E)**. These results are consistent with a model in which A06c-mediated inhibition of VO MNs, together with early excitation of VL MNs through A18b3, generates the normal onset delay between VL and VO muscles during forward crawling.

Using the same lateral-view recordings, we also examined the timing of VO muscle activity relative to other body-wall muscles **(Figure S14; Video S11)**. This broader comparison confirmed that disrupting the A27h–A18b3–A06c motif alters the normal temporal relationship between VO muscles and other muscle groups during forward crawling, further supporting a role for this PMN motif in maintaining behavior-specific intrasegmental muscle timing.

For the remaining intra- and intersegmental analyses, we imaged larvae from the ventral side. This orientation allowed larvae to crawl in a more natural posture and placed VO muscles in a favorable focal plane for quantifying their propagation across adjacent and non-adjacent segments. It also allowed us to compare VO15/16/17 activity with muscle 26 (M26) **(Figure 9F–H; Video S12)**. This comparison is grounded in circuit architecture: M26 is innervated by tonic MN26 and not by phasic MN ISNb/d/RP5 **(Figure S16)**, and MN26 receives minimal inhibitory input from A06c and substantially fewer excitatory synapses from A27h than VO MNs **(Supplementary Table 5)**. In control animals, M26 became active earlier than VO muscles. In contrast, consistent with the hypothesis that A06c imposes an early inhibitory delay on VO MNs during forward locomotion, silencing A06c or its upstream PMNs, A18b3 or A27h, led to more synchronous activation of VO and M26 muscles, indicative of premature VO activation.

We then quantified intersegmental coordination between adjacent segments, n and n+1, by measuring onset delay, peak delay, and two complementary metrics of activity overlap within individual crawl bouts: overlap of full width at half maximum (FWHM) activity windows and integrated area overlap **(Figure 9I–L; Figure S15; Video S12)**. In control animals, VO muscle activity propagated with consistent positive delays and limited FWHM and area overlap, reflecting orderly posterior-to-anterior peristaltic progression. Silencing any single PMN reduced onset and peak delays and increased both FWHM and area overlap, while dual PMN perturbations produced more pronounced compression of intersegmental delays and substantially increased overlap. These changes indicate aberrant synchrony of VO activity across adjacent segments.

We next examined coordination between non-adjacent segments separated by one intervening segment, n and n+2, to assess longer-range effects on peristaltic propagation **(Figure 9M–O; Figure S15; Video S12)**. As observed for adjacent segments, PMN perturbations reduced onset and peak delays and increased both FWHM and area overlap across non-adjacent segments. Dual perturbations produced the strongest compression of delays and the greatest increase in overlap. These results indicate that the A27h–A18b3–A06c motif is required to maintain appropriate temporal spacing of VO muscle activation across multiple segmental distances.

Together, these forward-crawling experiments demonstrate that the A27h–A18b3–A06c premotor motif is required for preserving the temporal structure of VO muscle activation. Disrupting any component of this tri-segmental network shifts VO activity earlier, reduces intersegmental timing delays, increases temporal overlap of VO activity across segments, and alters intrasegmental VO/VL timing during forward crawling.

We next examined how this forward-crawling motif relates to backward locomotion. During backward fictive locomotion, A27h and A18b3 are inactive, whereas A06c remains active but lacks its early activity peak **(Figure S12C)**. In addition, optogenetic validation showed that the backward-related PMN Canon/A18g can functionally activate A06c **(Figure S13E)**. We therefore reasoned that loss of A27h or A18b3 should have little effect on backward crawling, whereas A06c loss of function may disrupt backward VO coordination. Consistent with this prediction, A06c silencing reduced intersegmental onset delays and increased both FWHM and area overlap during backward crawling **(Figure S17)**. In contrast, individual A18b3 or A27h silencing, as well as dual A18b3/A27h silencing, did not substantially affect these parameters during backward crawling **(Figure S17)**. These results support the conclusion that A27h and A18b3 function preferentially in the forward-crawling premotor motif, whereas A06c contributes to both forward and backward locomotor coordination, potentially through behavior-specific upstream inputs.

### Disrupting the A27h–A18b3–A06c motif impairs protopodium dynamics and crawling efficiency during forward locomotion

Our calcium-imaging and connectomic analyses indicate that the A27h–A18b3–A06c premotor motif constrains the timing of VO muscle activation during forward crawling. Disruption of this motif shifts VO activity earlier, compresses intersegmental delays, and increases temporal overlap of VO activity across segments **(Figure 9)**. Because efficient protopodium folding and stride likely depend on properly coordinated activation and relaxation of VO and VL muscles across neighboring segments, we reasoned that disrupting this premotor motif would impair protopodium dynamics and reduce forward locomotor performance.

To test this, we silenced A06c, A27h, or A18b3 PMNs individually or in dual combinations using Gal4- or split-Gal4-driven expression of *UAS-GtACR1* and examined protopodium dynamics during forward crawling in intact larvae **(Figure 10A,B; Video S13)**. A06c split-Gal4 selectively targets A06c PMNs. Because 36G02-Gal4 labels A27h together with additional neurons, including M and A03g neurons [39, 41, 42], we also used an A27h split-Gal4 line developed in our lab [42]. Similarly, because 47E12-Gal4 labels A18b3 together with A18a and occasional sensory neurons [38], phenotypes were confirmed using an A18b3 split-Gal4 line. Unlike the lateral-view muscle-calcium-imaging experiments in **Figure 9**, these assays did not require simultaneous quantification of VO/VL phase relationships across the dorsoventral muscle field, allowing greater flexibility in combining Gal4 and split-Gal4 reagents for dual PMN silencing.

**Figure 10.**
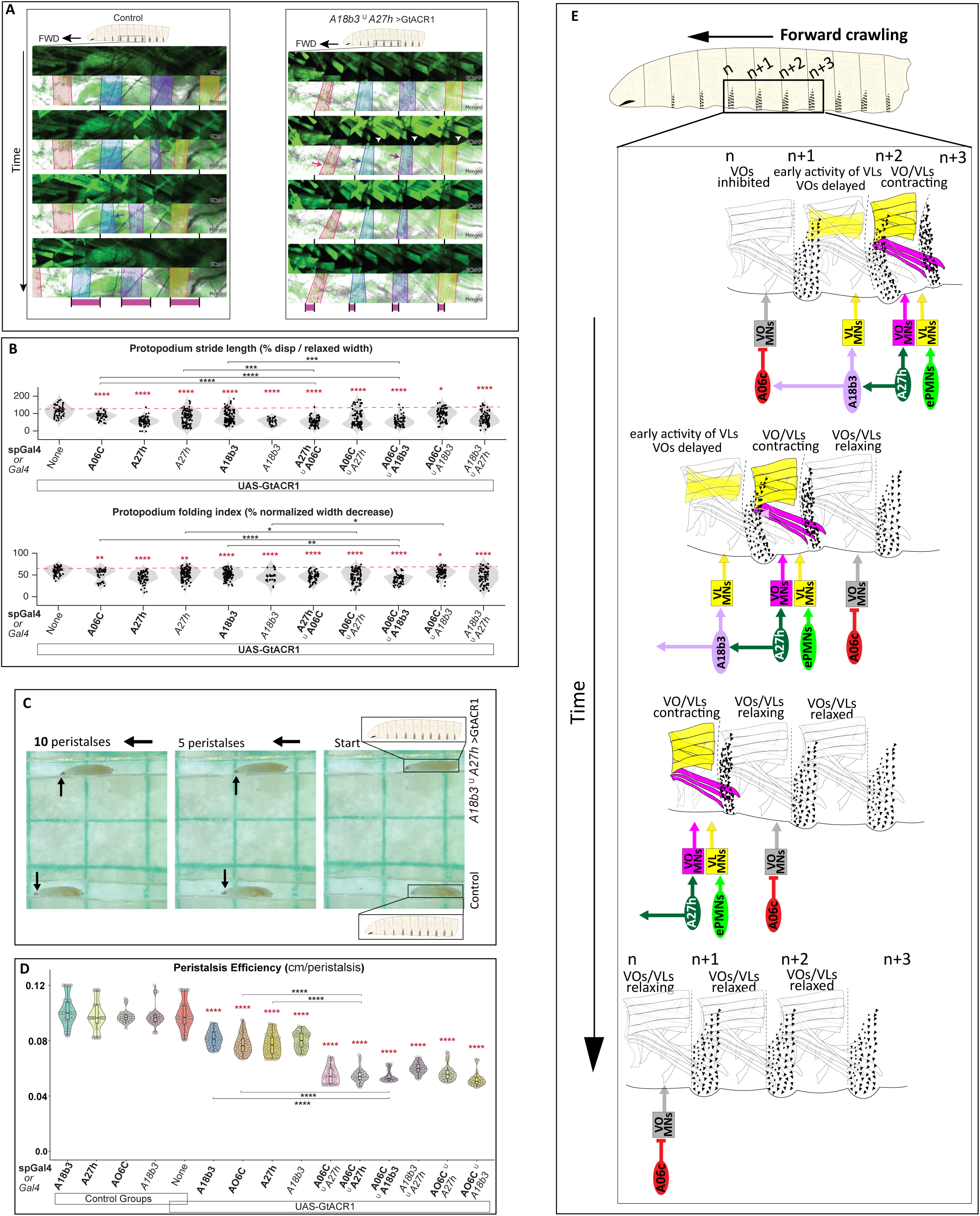
Disrupting the A27h–A18b3–A06c premotor motif impairs protopodium folding, stride length, and crawling efficiency during forward locomotion. **(A)** Time-lapse images of protopodium movement and VO muscle calcium activity (GCaMP6f) during forward crawling in control larvae and larvae with dual PMN silencing (*A18b3 ∪ A27h>UAS-GtACR1*). Protopodia from the same segment are color-coded; bottom magenta bars indicate stride length for a single crawl bout. Dual PMN silencing reduces stride length and induces multi-segmentally synchronous VO activation (white arrowheads). **(B)** Violin plots quantifying protopodium stride length (top) and folding index (bottom) following PMN silencing. Conventional Gal4 lines are shown in italics, and split-Gal4 lines are shown in boldface in the figure. For each genotype, N = 9–22 larvae were analyzed (1–3 protopodia per larva; n = 30–107 protopodia). Brown asterisks denote differences from control; black brackets indicate significant genotype comparisons. **(C)** Still images showing forward crawling performance after 5 and 10 peristaltic cycles in straight agarose tracks. Dual PMN-silenced larvae travel shorter distances per cycle. Grid spacing: 0.5 cm. **(D)** Violin plots of forward peristalsis efficiency, measured as distance traveled per cycle, for larvae expressing *UAS-GtACR1* under the indicated drivers (n = 40 larvae per genotype). (B,D) Statistics: Kruskal–Wallis tests followed by planned Wilcoxon comparisons versus each control with Holm–Bonferroni correction; brown asterisks indicate the most conservative adjusted *P* value. Exploratory all-pairs Dunn’s tests with Holm correction are shown by black brackets. Across panels (A–D), dual PMN silencing produces more severe defects than single PMN perturbations. Asterisks indicate *P* < 0.05, 0.01, 0.001, and 0.0001. Italic and boldface PMN labels in the figure denote conventional Gal4 and split-Gal4 lines, respectively. **(E)** Model summarizing how the A27h–A18b3–A06c motif defines the VO activation window during forward locomotion. A27h activity in posterior segments is proposed to recruit A18b3 in the adjacent segment, which in turn drives the first firing of A06c in the next anterior segment to transiently delay VO activation. In parallel, A18b3 primes VL MNs, while subsequent A27h-associated VO activation promotes protopodium swing.

High-resolution imaging revealed that silencing any single PMN significantly reduced protopodium folding and stride length during forward crawling **(Figure 10A,B; Video S13)**. Across conditions, dual PMN silencing produced stronger defects than single PMN silencing, consistent with broader disruption of the premotor motif. These results indicate that A06c, A27h, and A18b3 each contribute to proper protopodium mechanics during forward locomotion.

We next asked whether these protopodium-level defects translate into impaired whole-animal locomotion. Quantification of forward peristalsis efficiency, defined as the distance traveled per peristaltic cycle, showed that silencing individual PMNs reduced crawling efficiency, whereas dual PMN silencing produced the strongest deficits **(Figure 10C,D; Video S14)**. Thus, perturbing the A27h–A18b3–A06c motif compromises both local protopodium dynamics and whole-animal forward locomotor performance.

These behavioral defects are consistent with the muscle-timing defects observed in **Figure 9**, in which disruption of the A27h–A18b3–A06c motif caused premature VO activation, reduced intersegmental delays, and increased overlap of VO activity across segments. However, A27h and A18b3 are not expected to act solely through A06c or VO MN timing. A27h synapses onto VO MNs and A06c but also contacts additional PMNs, while A18b3 synapses onto A06c, additional PMNs, and VL MNs. Therefore, the stronger phenotypes observed after dual PMN silencing likely reflect combined disruption of VO timing, VL/VO coordination, intersegmental propagation, and additional premotor pathways in which A27h or A18b3 participate.

Together, these findings support a model in which the A27h–A18b3–A06c premotor motif is required for coordinated forward crawling by maintaining appropriate VO/VL timing, limiting aberrant overlap of VO activity across segments, and supporting normal protopodium folding and stride. A schematic summary of the proposed sequence linking PMN activity, VO/VL muscle timing, protopodium mechanics, and crawling output is presented in **Figure 10E**.

## Discussion

Animals generate flexible locomotor behaviors by coordinating shared muscles and motor neurons in behavior-specific ways. Here, we investigated how *Drosophila* larvae use ventral body-wall muscles to control protopodium dynamics during crawling. By combining calcium imaging, optogenetic perturbation, connectome-based circuit analysis, and biomechanical modeling, we show that VO muscles are major contributors to protopodium movement. VO activity is particularly important for protopodium folding, whereas both VO and VL activity contribute to protopodium stride length and forward crawling efficiency. We further identify a tri-segmental premotor excitation-inhibition motif that shapes VO timing during forward crawling.

Our findings establish protopodia as actively controlled, limb-like structures whose movement depends on coordinated ventral muscle activity. Although larvae lack discrete limbs, protopodia exhibit distinct folding, swing, and stance-like phases during crawling, with direction-specific patterns of detachment from and recontact with the substrate that parallel general principles of terrestrial locomotion [19, 21, 23, 43–45]. VO muscles are anatomically well positioned to influence these movements because they lie near the protopodia and have an oblique orientation across the ventral body wall. Consistent with this anatomy, optogenetic activation of VO MNs induces robust protopodium folding and ventral bending.

The functional perturbation experiments separate three related levels of locomotor output: protopodium folding, protopodium stride length, and whole-animal crawling efficiency. Folding provides a relatively direct measure of local protopodium deformation, whereas stride length and peristaltic efficiency are broader locomotor outputs that may be influenced by additional factors, including body-wall deformation, intersegmental coordination, substrate contact, visceral pistoning, and force transmission across adjacent segments. This distinction helps explain why VL MN silencing reduces protopodium stride length and crawling efficiency without producing the same folding defect as VO MN silencing. Thus, VO muscles appear to be especially important for the local fold-and-lift component of protopodium movement, whereas VO and VL muscles together shape the broader mechanical output of crawling.

The muscle calcium-imaging and length-tracking data further show that VO/VL timing differs between forward and backward crawling. During forward crawling, VL activity precedes VO activity, whereas during backward crawling VO and VL activity are more closely synchronized. Our mechanical model provides a functional explanation for this difference, suggesting that delayed VO activation during forward crawling is not simply a timing feature, but may be important for protopodium gait. Because contraction within a segment can transmit force toward both neighboring protopodia, productive movement requires one side of the contracting segment to remain mechanically stabilized while the other side moves. During forward crawling, early VL activity in the next anterior segment may increase local resistance and help stabilize the anterior protopodium in stance before VO/VL contraction in the active segment folds and swings the posterior protopodium. By contrast, during backward crawling, early VO/VL activity in the next posterior segment may provide the stabilizing influence needed for anterior protopodium swing. Thus, behavior-specific VO/VL timing may help determine which neighboring protopodium remains anchored and which protopodium moves.

Kinematic and robotic modeling support this mechanical interpretation. The inverse kinematic model predicted VO length dynamics that closely matched in vivo measurements, whereas predicted VL length changes were smaller than those observed in vivo, suggesting that VO shortening and VL rotation can account for much of the protopodium trajectory within the model geometry. Robotic simulations provided a reduced test of mechanical sufficiency: VO silencing strongly reduced forward displacement, whereas VL silencing preserved displacement but produced skidding; during backward crawling, silencing either VO or VL reduced displacement. Because this model does not fully capture passive tissue deformation, detailed protopodial grip, or neighboring-segment anchoring, it is best interpreted as a simplified mechanical test rather than a complete reconstruction of larval locomotion.

Having established a functional rationale for delayed VO activation during forward crawling, we then asked how this delay is generated at the circuit level. The connectome and calcium-imaging data identify a tri-segmental premotor motif that is well positioned to impose this forward-specific VO timing. In this motif, premotor excitation and inhibition are organized across adjacent segments so that VO MN activation is temporally constrained relative to VL activity, neighboring-segment anchoring, and the progression of the forward locomotor wave.

At the circuit level, we identify A06c and A27h as major premotor inputs to VO MNs and show that A18b3 links this VO-regulating network across segments. During forward fictive locomotion, the inhibitory PMN A06c exhibits two activity peaks that bracket the VO MN activation window. Consistent with previous reports [1, 4, 38], A27h and A18b3 are recruited only during forward locomotion, with timing and connectivity consistent with a tri-segmental A27h–A18b3–A06c motif that constrains VO activation. Functional optogenetic experiments in isolated CNS preparations support key monosynaptic connections predicted by the EM connectome, including A27h→A06c, A27h→A18b3, and A18b3→A06c. These experiments also validate Canon/A18g→A06c, identifying a candidate backward-related input to the shared inhibitory PMN A06c. Together, these data indicate that premotor excitation and inhibition are organized across segments to regulate VO MN timing.

Perturbing this motif disrupts the temporal organization of forward crawling. Silencing A06c, A27h, or A18b3 shifts VO activity earlier, reduces intersegmental timing delays, increases temporal overlap of VO activity across segments, and impairs forward protopodium folding, stride, and crawling efficiency. Dual PMN perturbations produce stronger phenotypes than single perturbations, consistent with broader disruption of the premotor network. This result is expected because A27h and A18b3 are not dedicated exclusively to A06c or VO MNs, but are embedded within previously described premotor networks [1, 4, 5, 7, 46]. A27h provides excitatory input to VO MNs and A06c, forming an incoherent type-1 feedforward loop that is positioned to both promote VO MN activity and constrain its timing through A06c-mediated inhibition. A27h also interacts with additional interneurons implicated in locomotor coordination, including A27j2/Gdl [1, 4, 5, 7, 46]. Similarly, A18b3, previously described as the forward-active CLI1 neuron [38], provides input to A06c but also contacts additional premotor and motor targets, including A14a/iIN-1, A27j2/Gdl, and multiple VL MNs [1, 4, 5]. Thus, perturbing A27h or A18b3 is expected to affect not only VO timing through A06c, but also broader premotor pathways involved in intersegmental coordination, stride rhythm, and muscle-group timing. These features support a modular and hierarchical organization of the larval premotor network and emphasize that the A27h–A18b3–A06c motif should be interpreted within its broader circuit context.

These findings also clarify how the forward-crawling motif relates to backward locomotion. A27h and A18b3 are inactive during backward fictive locomotion, whereas A06c remains active but lacks its early inhibitory peak. Consistent with these activity patterns, silencing A27h or A18b3 does not substantially alter backward VO coordination, whereas A06c silencing reduces intersegmental timing delays and increases overlap of VO activity during backward crawling. Thus, A27h and A18b3 appear to function preferentially in the forward-crawling motif, while A06c is reused across crawling directions in a behavior-specific manner. The functional Canon/A18g→A06c connection provides one candidate route through which backward-related circuitry could recruit this shared inhibitory PMN.

Several questions remain open. First, although our backward-crawling analyses identify behavior-specific differences in PMN recruitment and A06c timing, they do not provide a complete circuit-level explanation of backward crawling. Future work should determine how backward-active PMNs, including Canon/A18g and Ifb-Bwd [9, 11], interact with A06c and other premotor targets. Second, the upstream mechanisms that select forward- versus backward-crawling circuit states remain unresolved. Command-like neurons, sensory feedback, and neuromodulatory inputs may all contribute to selecting which premotor pathways are recruited and how shared downstream targets such as A06c are temporally organized. Finally, future mechanical models that incorporate detailed substrate grip, passive body-wall deformation, and neighboring-segment interactions should further clarify how circuit timing is transformed into protopodium forces and whole-animal locomotion.

More broadly, our findings support a model in which locomotor flexibility arises through behavior-specific organization of premotor excitation and inhibition within shared motor networks, rather than through entirely dedicated MN pools. Similar principles operate in other organisms, including vertebrate spinal circuits that use excitatory and inhibitory interneurons to shape MN burst timing and phase relationships [47–50], limbed animals that flexibly modify gait patterns [19, 20, 23, 24, 43, 51–58], annelids and mollusks that use multifunctional circuits to generate distinct motor outputs [59–62], and zebrafish and mammals in which interneuron recruitment and commissural pathways regulate speed, direction, and bilateral coordination [53, 55, 56, 63–66]. Although the larval motor system differs in anatomy and biomechanics, the premotor motif described here illustrates how circuit timing can be transformed into muscle coordination, body mechanics, and behavior. By linking a connectome-defined premotor motif to MN activity, muscle timing, protopodium mechanics, and locomotor output, this work provides a framework for understanding how shared neuromuscular systems generate flexible behavior.

## Acknowledgements

We thank Jim Truman for unpublished fly lines targeting A18b3 and A06c neurons. We thank Chris Q. Doe, Wanhe Li, and Lewis Sherer for comments on early versions of the manuscript. We thank Chris Q. Doe, Todd Laverty, and Gerry Rubin for fly stocks. We thank Albert Cardona, Richard D. Fetter, and the HHMI Janelia FlyEM Project Team for providing the raw data for the whole-CNS EM volume. We thank Akira Fushiki, Eri Hasegawa, and Maarten Zwart for annotating neurons, and Keiko Hirono for generating transgenic constructs. This study was supported by NIH-NINDS 1R01NS142268-01A1, the Texas A&M University Division of Research Targeted Proposal Teams (TPT) funding program, Zarin Ascend FY25 and FY26, and two internal grants from the Texas A&M College of Arts and Sciences, Zarin STRP FY24 and FY25.

## Author Contributions

*Conceptualization* - Y.H., A.S., I.S.G., and A.A.Z.

*Data curation* - A.S., Y.H., L.O., L.C., V.E.M., P.M.M., N.B., S.A., D.M.P., Y.W., I.S.G., and A.A.Z.

*Formal* Analysis - A.S., Y.H., L.O., and A.A.Z.

*Funding Acquisition* - A.A.Z.

*Investigation* - A.S., Y.H., L.O., D.M.P., and A.A.Z

*Methodology* - A.S., Y.H., L.O., L.C., V.E.M., P.M.M., N.B., S.A., Y.W., D.M.P., I.S.G., and A.A.Z.

*Project Administration* - A.A.Z.

*Resources* - A.S., Y.H., L.O., and A.A.Z.

*Software* - A.S., Y.H., D.M.P., I.S.G., and A.A.Z.

*Supervision* – I.S.G., and A.A.Z.

*Validation* - A.S., Y.H., D.M.P., I.S.G., and A.A.Z.

*Visualization* - A.S., Y.H., L.O., Y.W., D.M.P., I.S.G., and A.A.Z.

*Writing, original draft* - Y.H., and A.A.Z.

*Writing, review & editing* - A.S., Y.H., L.O., D.M.P., and A.A.Z.

## Methods

### Fly Genetics

A complete list of fly stocks is provided in the Key Resources Table (Supplementary Information).

#### Inclusion and ethics statement

This study used Drosophila melanogaster as a model organism. No human participants were involved. All animal procedures were conducted in accordance with institutional guidelines and approved protocols.

### Confocal Muscle Calcium and Protopodium Imaging

Larval imaging was performed on a 25 × 25 × 2 mm agarose gel pad (1.5% agarose) used as the imaging substrate. L2–L3 larvae (equal numbers of males and females) were washed in distilled water and individually placed on the agarose pad, then gently constrained with a 22 × 22 mm coverslip. Larvae were positioned lateral or ventral side up with VO muscles in the focal planes. Imaging was conducted on a Zeiss LSM900 confocal microscope using either a 5× or 10× objective. Muscle GCaMP fluorescence was imaged using a 488 nm laser (AF488 channel). For simultaneous muscle and protopodium imaging, a brightfield channel was acquired in parallel. During optogenetic silencing of MNs or PMNs expressing GtACR1, the 488 nm laser was also used to activate the inhibitory channel. Muscle calcium activity and protopodium movements were recorded during forward or backward peristaltic crawling. Image sequences were acquired as rectangular frames (256 px in length with variable width) spanning the full dorsoventral extent of the body wall, with more than three abdominal segments captured in each movie.

### Muscle GCaMP Data Analysis

For analysis of muscle GCaMP signals in Figure 1 and Figure S2, segment-level activity was defined using a principal component analysis (PCA)–based approach, as previously described (Zarin et al., eLife, 2019 [1]). Because it was not possible to image all ~30 muscles within a hemisegment simultaneously, we analyzed segments containing at least one representative muscle from each of the four previously defined co-active muscle groups. Fluorescence traces from these representative muscles within a segment were subjected to PCA, and the first principal component (PC1) was used as a composite measure of segmental activity. Segmental onset and offset were defined from this PCA-derived trace and used to align individual crawl cycles to a normalized 0–100% time axis, with normalized time points 35 and 65 corresponding to segmental onset and offset, respectively.

For intra-segmental timing analyses in **Figure 9B–H**, ΔF/F traces were analyzed on a per-bout basis without PCA. For lateral-view recordings in **Figure 9B–E**, the timing relationship between pooled VO muscles 15/16/17 and pooled reference VL muscles 12/13 within the same segment was quantified using a custom MATLAB pipeline (`intra_segmental_VO_vs_M12_13_multigroup_v6_QC_FIXED_v2.m).` Raw onset and peak timing were computed in seconds, and normalized onset and peak positions were computed on a 0–100% crawl-cycle time axis. VO15/16/17 traces were pooled and compared with pooled M12/13 reference traces from the same segment and crawl bout. Timing differences were calculated as VO minus M12/13, so positive values indicate that VO activity occurred after M12/13 activity. Files were excluded if onset indices were undefined, invalid, or occurred at or after peak activation.

For ventral-view recordings in **Figure 9F–H**, the timing relationship between pooled VO15/16/17 activity and muscle 26 (M26) activity was quantified within the same segment using a separate custom MATLAB pipeline (`INTER_RF_v11_INTRplots_INTERplots_VOonly.m).` M26 was used as an intrasegmental reference muscle because it is innervated by tonic MN26, does not receive detectable phasic ISNb/d/RP5 input, and receives little or no direct input from the key VO-regulating PMNs analyzed in this study. Onset and peak delays were calculated as VO15/16/17 minus M26, so positive values indicate that VO activity occurred after M26 activity.

For intersegmental coordination analyses in **Figure 9I–O**, VO15/16/17 activity was quantified across segment pairs separated by user-defined intersegmental gaps using `INTER_RF_v11_INTRplots_INTERplots_VOonly.m.` Adjacent segment pairs were analyzed with gap = 1, and every-other segment pairs were analyzed with gap = 2. Pairing was restricted to traces from the same crawl bout and matched segmental sequence. For each segment pair, onset delay and peak delay were calculated in both real time and cycle-normalized time. Intersegmental overlap was quantified using two complementary metrics. Full-width at half-maximum (FWHM) overlap was calculated from the overlap between the two segments’ FWHM activity intervals. Integrated area overlap was calculated as the integral of the point-wise minimum of the two normalized calcium traces, thereby incorporating both overlap duration and signal amplitude. Unless otherwise stated, plotted Figure 9 metrics use cycle-normalized values.

### Protopodium Data Analysis

Protopodium folding and displacement were quantified using custom MATLAB scripts. A linear region of interest (ROI) was placed across each protopodium, perpendicular to the denticle belts, and the ROI length and position were adjusted frame by frame to track protopodium movement and changes in width. For analyses in Figure 2, VO muscle calcium activity and protopodium kinematics were quantified from the same animals and imaging sessions. For each crawl bout, three consecutive segmental regions were analyzed, and VO activity and protopodium width traces were aligned to the peak activity window of the middle VO segment. The onset and offset of this reference VO activity—defined as the half-width before and after the activity peak—were aligned to normalized time points 35 and 65, respectively, and all VO and protopodium traces from the same bout were resampled to this normalized 0–100% time axis.

For analyses in Figure 6 D-G and Figure 10B, protopodium folding and stride metrics were quantified using a dedicated custom MATLAB pipeline (`PROTO_RF_v3.m`). Protopodium width and midpoint position were extracted frame by frame for individual crawling bouts. Folding index was defined as the percent reduction from relaxed to maximally folded width. Stride length was calculated as the net protopodium displacement over the bout (xmid_end − xmid_start) and normalized to relaxed width, with forward direction determined independently for each bout from the sign of peak swing velocity. Group-level differences were assessed using Kruskal–Wallis tests. Planned comparisons versus WT controls, as well as additional non-WT pairwise comparisons, were evaluated using two-sided Wilcoxon rank-sum tests with Holm–Bonferroni correction.

### Muscle length measurement

Muscle length measurement (**Figure 1I**) was done with the same MATLAB scripts as the muscle GCaMP measurement, where linear ROIs were drawn across the length (along its direction of contraction) of each muscle. VL and VO muscle length was measured for each segment, and the time point of minimum length (“peak”) of VL was defined as normalized time 50, and peak +/- 0.5 peak-width time points were defined as normalized time 65 and 35, respectively. VO length was normalized to this 0-100 time scale, and data from different segments was compiled and plotted. Muscle length was normalized to a 0-1 scale, where 0 is minimum length (when muscle is fully contracted) and 1 is maximum length (resting).

### Modeling overview

To examine the mechanical contribution of ventral muscles to protopodium movement during forward crawling, we developed a reduced-order planar kinematic model of a single larval segment. VO and VL muscle groups were represented as elastic elements within a four-bar linkage, with additional abstract elements included solely to maintain kinematic closure. Muscle elements were modeled as linear springs, and all joints were assumed to be frictionless. Model parameters and constraints were derived from in vivo muscle calcium imaging, including experimentally measured ranges of muscle shortening and VL orientation. Forward kinematics were used to predict protopodium displacement from measured muscle dynamics, whereas inverse kinematics were used to estimate the muscle length changes required to reproduce observed protopodium trajectories. Model predictions were validated against experimental data and further tested in a physics-based robotic simulation.

### Optogenetic MN activation and muscle/protopodium imaging (concise)

For protopodium imaging with optogenetic MN (MN) activation (Figure 5), L3 larvae expressing 44H10::GCaMP, UAS-Chrimson-mCherry, and an MN-Gal4 driver were imaged using a Zeiss LSM900 confocal microscope. Larvae were raised in the dark and fed 100 µL of 0.5 mM all-trans-retinal (ATR) 24 h before imaging.

A 10 × 10 mm, 1.5% agarose gel containing a ~0.7 mm-wide, 1 mm-deep groove was prepared using the combined edge of five #1 coverslips. Individual larvae were placed ventral-side up in the groove, covered with a single coverslip to restrict movement, and a small drop of distilled water was added to prevent adhesion. Muscle calcium activity was imaged using a 488 nm laser (AF488 channel), which also activated Chrimson in MNs, while a brightfield channel simultaneously recorded protopodium movement.

The first imaging frame was defined as the resting (“light-off”) state and used to determine resting VO muscle length and protopodium width. Minimum VO length and protopodium width within the first 30 frames were defined as “light-on” values (Figure 5). All measurements were normalized to light-off values. VO length reduction was calculated as 1 − (light-on VO length / light-off VO length), and protopodium width reduction as 1 − (light-on protopodium width / light-off protopodium width).

Full-body imaging (Figure 5A) was performed using an iPhone 14 mounted on a Zeiss Stemi 508 microscope. Larvae were placed lateral-side up on 1.5% agarose under dim ambient light. Uniform white-light stimulation was provided by an 18 W LED positioned beneath the gel. Responses were recorded for ~10 s, with dorsal or ventral curvature classified as C-shaped bending and axial shortening without curvature classified as body shortening.

### Larval crawling efficiency analysis

A two-layer 1.5–2% agarose gel was prepared in a 140 mm (diameter) × 20 mm (depth) glass Petri dish. A 40 mL base layer was poured and allowed to solidify, after which a groove mold was placed on top and covered with an additional 30–40 mL agarose layer. The groove mold was designed using freely available CAD software (Tinkercad) and 3D printed in PLA using an Ultimaker S3 printer with Cura software (0.8 mm thickness, 15% infill; “Fast” setting). The mold consisted of a 111 × 75 × 1 mm rectangular base with twelve 75 × 1.2 × 5 mm (length × width × depth) ridges spaced ~8.7 mm apart, and a handle on the top surface for easy removal.

Larvae expressing UAS-GtACR1 crossed to w1118 (control) or to MN-Gal4 or PMN-Gal4 drivers (MN>GtACR1 for Figure 6; PMN>GtACR1 for Figure 10) were raised in the dark and fed 100 µL of 0.5 mM all-trans-retinal (ATR) 24 h before experiments. Wandering 4mm long L3 larvae were rinsed and gently placed into the grooves. Crawling distance was measured using translucent graph paper with 5 mm grids placed beneath the gel. An 18 W square LED (Oeegoo) positioned under the dish provided uniform white-light illumination to activate GtACR1.

Larval crawling was recorded for 1 min using an iPhone 14 camera (30 fps, 1440 × 1920 pixels). Videos were analyzed in Fiji. Forward crawling efficiency was defined as the distance traveled per peristaltic cycle and calculated as the inverse of the number of peristalses required to traverse 1 cm. Peristalses were manually counted, and each animal was analyzed only once.

### Calcium imaging in neurons

Dual-color neuronal imaging was done in isolated larval CNS preparations. L2–L3 larval CNS preparations were dissected in HL3.1 saline and mounted on 12 mm round Poly-D-Lysine Coverslips and imaged on a Zeiss LSM900 confocal microscope. For dual-color imaging, two neurons differentially expressing GCaMP6m (AF488 channel) and jRCaMP1b (AF568 channel) were imaged in the same focal plane and ROIs were placed on individual neurons.

Quantification of neuron fluorescence activity was done on Zeiss Zen software as CSV files and imported to MATLAB for analysis. DF/F is defined as (F-F0)/F0, where F0 is the 10^th^ percentile of the first 3^rd^ of the trace. All DF/F values were normalized into a 0-1 scale for graphing. Data from different fictive crawl cycles was aligned to a 0-100 normalized time scale in reference to CQ-MNs (i.e., *CQ-LexA, LexAop-GCaMP6m*). Specifically, the onset and offset of GCaMP6m signal (CQ-MNs) were defined as 0.5 peak-width before and after the peak and used as normalized time 35 and 65 in alignment of both GCaMP6m and jRCaMP1b signal. For A18b3 (n+1) vs A27h (n+2) (Figure 7F), a new set of data collected from *36G02-LexA>Aop-GCaMP6m, A18b3 split-Gal4>UAS-jRCaMP1b* was used (15 fictive waves, 4 animals). A27h (n+1) was measured together with A27h (n+2) and A18b3 (n+1), and was used for defining 0-100 normalized time. The onset and offset of A27h (n+1) signal were defined as 0.5 peak-width before and after the peak and used as normalized time 35 and 65 in alignment of A27h (n+1), A27h (n+2) and A18b3 (n+1) signal.

### Optogenetic validation of EM connectome

For optogenetic activation of upstream PMNs (A27h or A18b3) while imaging downstream PMN activity (A06c or A18b3) **(Figure 8)**, offspring from the following crosses were used: 1) *36G02-LexA; Aop-Chrimson-mCherry, UAS-GCaMP6m* × *A18b3 split-Gal4*, 2) *36G02-LexA; Aop-Chrimson-mCherry* × *A06c split-Gal4*, and 3) *A06c-LexA; Aop-GCaMP6m, UAS-Chrimson-mCherry* × *A18b3 split-Gal4*. Larvae were fed 100 µL of 0.5 mM ATR 24 h before the experiment. L3 larval CNS were dissected in HL3.1 saline alone, or HL3.1 with 3 µM Tetrodotoxin (TTX), and imaged under the Zeiss LSM900 confocal microscope with an AF488 channel to record GCaMP6m in A06c or A18b3. After 0.25 s of baseline imaging, a pulse of 561 nm laser was applied to the entire imaging field for 0.5 s to activate Chrimson in A27h or A18b3, after which the imaging of AF488 channel was resumed. Pre- and post-bleaching GCaMP6m traces from different animals were aligned and the mean of raw DF/F from different animals were plotted. The F0 for calculating DF/F is defined as the 10^th^ percentile value during the 0.25 s baseline imaging. For validating the effect of TTX, *CQ-LexA>Aop-GCaMP6m* larval CNS was dissected in HL3.1 saline alone, or HL3.1 with 3 µM TTX and recorded with a single AF488 channel to examine the spontaneous firing of CQ MNs (fictive motor activity).

### Fly husbandry

All flies were reared at 25°C and 50% relative humidity with a 12 h/12 h light/dark cycle in molasses fly food. For crosses involving GtACR1 and Chrimson, all offspring larvae were raised in the dark until the experiment. All comparisons between groups were based on studies with flies grown, handled, and tested together. Larvae for optogenetic experiments were fed with all-trans retinal (ATR) 24 hours before experiments (100 µL of 0.5 mM ATR solution for each vial).

### Stochastic neuron labeling with MCFO

The MultiColor FlpOut (MCFO) approach developed by Nern et. al. was used [31]. A18b3 or A06c split Gal4 lines were crossed with *R57C10-Flp, UAS-MCFO (HA-V5_FLAG)* to induce marker expression by Flp-recombinase excision randomly in A18b3 or A06c neurons. Confocal fluorescence imaging was done in isolated CNS where a sole A18b3 or A06c (among all VNC segments) was labeled to show single-neuron morphology.

### Immunohistochemistry

For larval body wall preparations, larvae were dissected along the dorsal midline in HL-3 saline, with the brain and internal organs removed. The body wall (“filet”) was pinned, fixed in 4% paraformaldehyde (PFA) for 10 min, washed in PBS containing 0.1% Triton X-100, and blocked in 5% normal donkey and goat serum. Primary antibodies were applied for 2 h at room temperature or overnight at 4 °C. Samples were 3X washed in PBST and incubated with secondary antibodies together with CF647–phalloidin to label muscle fibers, followed by additional PBST washes. Preparations were mounted in Fluoromount and stored at −20 °C prior to imaging.

For isolated larval CNS staining, L2–L3 CNS were dissected in HL3.1 saline and mounted onto poly-D-lysine–coated coverslips. Samples were fixed in 4% PFA diluted in PBST, washed, and blocked in 5% normal goat and donkey serum. Primary antibodies were applied overnight at 4 °C, followed by PBST washes and incubation with secondary antibodies for 2 h at room temperature. After final washes, samples were mounted in Fluoromount-G (SouthernBiotech) and stored at −20 °C or imaged by confocal microscopy. Details of all antibodies, sources, and dilutions are provided in the Key Resources Table.

### Electron microscopy and CATMAID reconstructions

The TEM volume is for a newly hatched first instar (L1) larval nervous system (Ohyama et al., 2015). This dataset is available by request from Albert Cardona (Cambridge University). Neurons were reconstructed in CATMAID (Saalfeld et al., 2009) [68] using a Google Chrome browser as previously described (Ohyama et al., 2015). Figure 7A,D and S8 were generated using CATMAID 3D widget, Python script, and Adobe Illustrator (Adobe, San Jose, CA).

### Statistics

All statistical analyses were performed in MATLAB (The MathWorks, Inc., Natick, MA) or RStudio. For the comparison between forward and backward crawling in wild-type larvae, data from Zarin et al. 2019 were used, and Student’s *t*-test was used for the boxplot in **Figure 1H**. Kruskal–Wallis tests were used to assess overall group effects, followed by planned Wilcoxon rank-sum comparisons versus WT with Holm–Bonferroni correction for hypothesis-driven analyses, or Dunn’s post hoc tests when all pairwise group differences were explored. In line plots, solid lines represent the mean of all measurements from the indicated animals or experiments, and shaded regions represent the standard deviation unless otherwise stated. This applies to the calcium-activity, muscle-length, and kinematic traces shown in **Figures 1G, 1I, 2C, 7E, 7F, 8A–D, 9B, and 9F**. Equal numbers of male and female animals were used in each experiment. Sample size was pre-determined by a power analysis with effect size d = 0.5, α = 0.05 and power = 0.8 in the freeware G*Power3.1 ([69] Faul et al. 2009).

## Data availability

The raw calcium imaging and behavioral datasets generated and analyzed in this study, including MATLAB (.mat) and Microsoft Excel (.xlsx) files, have been deposited in Dryad. The repository is currently accessible to editors and reviewers through a private peer review link and will be made publicly available upon publication of the article.

