## Supplementary info for "A Premotor Excitation-Inhibition Motif Shapes Protopodium Dynamics During Drosophila Larval Forward Crawling"

### Supplementary Results: Expanded mechanical modeling of protopodium gait generation

To further examine how ventral oblique **(VO)** and ventral longitudinal **(VL)** muscles contribute mechanically to protopodium movement during forward and backward crawling, we developed a kinematic model based on an earlier modeling framework [1]. This expanded version includes updated parameterization, additional comparison with in vivo measurements, and physics-based robotic simulations.

Because muscles can both actively contract and passively deform, we represented the VO–VL–protopodium system using elastic elements in a reduced-order biomechanical model [2, 3]. Larval crawling occurs largely in the sagittal plane, and the ventral musculature is bilaterally symmetric; therefore, we approximated the system as a planar four-bar mechanism [4] **(Figure 3A)**. In this model, VO and VL muscle groups are represented as deformable links, whereas additional auxiliary links preserve the four-bar geometry. These auxiliary elements are mathematical/mechanical constraints and are not intended to represent specific biological muscles.

This planar simplification assumes that sagittal-plane motion captures the dominant component of protopodium movement. This assumption is supported by in vivo tracking of ventral muscles and protopodia, which revealed limited out-of-plane displacement during crawling. The VL group rotated around the anterior joint by approximately 0–8°, while the corresponding vertical element remained nearly aligned with the body axis **(Figure 3A,B)**. Based on these observations, the model permits VL rotation while constraining selected elements to contraction and extension. Additional assumptions, including uniform elastic stiffness and frictionless rotary joints, are described in the Methods.

For forward crawling, we analyzed videos from five larvae and tracked joint positions and muscle attachment points **(Figure 3B)**. These measurements provided VL horizontal angle, VO and VL muscle length changes **(Figure 3C)**, and the corresponding protopodium path **(Figure 3D)**. In agreement with calcium-imaging and muscle-length data, VL shortening preceded VO shortening during forward crawling. When the experimentally measured VO/VL lengths and VL angle were used as model inputs, the resulting protopodium path was similar to the measured path, with small differences likely arising from tissue deformation, substrate interactions, or forces from neighboring segments not included in the simplified model **(Figure 3D)**.

We then used the inverse model to estimate the muscle dynamics needed to produce the measured protopodium path **(Figure 3E–G)**. The predicted VL angle followed the observed temporal pattern and remained within the experimental range **(Figure 3E)**. Predicted VO length dynamics closely resembled the measured VO dynamics **(Figure 3G)**, whereas the predicted VL length changes were smaller than the VL shortening measured in vivo **(Figure 3F)**. Thus, within the constraints of the four-bar geometry, VO shortening combined with VL rotation can explain much of the forward protopodium path without requiring the full extent of VL shortening observed in vivo. When the predicted muscle dynamics were fed back into the model, they regenerated a protopodium path close to the experimental path **(Figure 3H)**.

The same modeling strategy was then applied to backward crawling. We tracked joints and muscle lengths from backward-crawling videos and used the extracted length changes and angles as model inputs without changing the model parameters **(Figure 3I)**. The model output reproduced the measured backward protopodium path reasonably well, indicating that the same geometric framework can describe protopodium movement in both crawling directions **(Figure 3J)**. In the inverse analysis, the predicted VL angle remained within the observed range **(Figure 3K)**, and the predicted VO length dynamics were similar to the measured VO dynamics **(Figure 3M)**. In contrast, predicted VL length changes were again smaller than those measured in vivo **(Figure 3L)**. Regenerating the backward protopodium path from the predicted muscle dynamics produced a close match to the measured path **(Figure 3N)**. These results indicate that the kinematic model captures key geometric features of both forward and backward protopodium movement and repeatedly assigns a larger trajectory-generating role to VO shortening than to VL shortening.

To test whether these modeled muscle dynamics could generate movement in a physical context, we implemented a segmented robotic model made from repeated four-bar modules interacting with a frictional surface **(Figure 3O–O″, Video S5)**. The goal of this simulation was not to reproduce larval locomotion quantitatively in every respect. In particular, the model does not fully capture free-crawling speed, absolute cycle duration, whole-animal displacement, all body-wall muscles, or the full complexity of substrate contact. Rather, it provides a simplified mechanical test of whether measured or model-predicted VO/VL dynamics can produce segmental displacement.

For mechanical-silencing simulations, the silenced muscle element was not removed from the model. Instead, its active length command was blocked and the corresponding piston was treated as a passive elastic element. Thus, the silenced VO or VL element could still shorten or extend in response to forces generated by neighboring segments and then return toward its neutral length. This implementation approximates passive deformation of non-activated muscle while allowing us to test the effect of removing active VO or VL contraction.

In the forward-crawling simulation, control inputs derived from in vivo measurements produced an advance of approximately 70% of a relaxed segment width per peristaltic cycle **(Figure 3P)**. Removing active VO contraction in silico strongly reduced forward displacement **(Figure 3P′)**. By contrast, removing active VL contraction preserved much of the forward displacement, but the model exhibited skidding, likely due to reduced VL-linked body-wall deformation and rearward force balance **(Figure 3P″)**. Inputs generated by the kinematic inverse model also produced forward movement, with displacement similar to the VL-silenced simulation **(Figure 3P‴)**. These simulations support the idea that VO contraction is a major mechanical contributor to forward protopodium gait, whereas VL contraction influences additional body-wall and substrate-interaction mechanics that are not captured by protopodium displacement alone.

We performed analogous physics-based simulations for backward crawling **(Figure 3Q–Q‴)**. Compared with forward crawling, backward displacement was more sensitive to the simplified substrate-contact assumptions of the robotic model. One likely reason is that backward crawling may depend more strongly on protopodial grip or adhesion, because oblique VO-mediated forces must help pull the segmental chain in the opposite direction. Consistent with recent measurements of ground-reaction forces in freely crawling larvae [5], this suggests that detailed substrate interactions are likely important for accurately reproducing backward locomotion. Because the current robotic model represents substrate contact using simplified frictional interactions and does not explicitly implement protopodial adhesion, passive tissue deformation, or neighboring-segment anchoring, backward displacement in the simulation should be interpreted as a mechanical-sufficiency test rather than a quantitative reconstruction of larval backward crawling.

Even with these limitations, in silico removal of either VO or VL contraction reduced backward displacement relative to the control simulation **(Figure 3Q′–Q″)**. This indicates that, under the simplified robotic conditions, backward displacement depends on both ventral muscle components. Simulations driven by model-predicted muscle dynamics produced lower backward displacement **(Figure 3Q‴)**, likely because passive tissue deformation, detailed substrate grip, and neighboring-segment mechanics are not fully implemented in the current model.

Overall, the kinematic and robotic analyses provide complementary evidence that VO muscles make a major mechanical contribution to protopodium gait generation. They also indicate that VL muscles, substrate interactions, and neighboring-segment mechanics shape displacement, especially during backward crawling. Because the robotic model is deliberately simplified, it should be interpreted as a mechanical sufficiency test rather than a complete quantitative reconstruction of larval crawling. The model focuses on the mechanical requirements for protopodium gait generation and does not address other known functions of VL muscles during forward crawling, including visceral pistoning and intersegmental body-mass transport [6]. These model-based predictions are tested in the main text using motor neuron-specific optogenetic manipulations in vivo.

**Supplementary Material**

Assumptions, Constraints, and Limitations

The symmetric nature of the muscle group motivates the primary assumption that the lower half of the abdomen can be approximated with a planar mechanism. Calcium imaging provides muscle motion of the sagittal plane, which is reasonable to use in the planar mechanism aligned with the sagittal plane. Moreover, although muscles exhibit active contraction, they are assumed to be mechanical springs. Thus, it is reasonable to assume that the system tends to stay at a minimum energy level by relaxing muscles as much as possible. This assumption is used in the model to find the minimum energy states during locomotion to obtain the natural movements of the protopodium.

The limits of muscle contraction and extension are obtained from the sample data. Each muscle shows different limits, and the model uses these limits in the model and the simulations to retain realistic output. The minimum muscle lengths were identified as $l_{lower} = 0.6295, 0.4744, 0.7632, 0.8896, 1.0621$ and the maximum was $l_{maximum} =$

$1.0124, 0.6006, 1.4144, 1.2394, 1.6678$. Here, we considered the hypothetical springs ($l_1 to\ l_5$) in the mechanism as well. The angle $\theta$ limits were identified as $[-0.6190^0, 8.4375^0]$. Furthermore, since the proposed model is a kinematic description of the muscle group, it cannot account for passive deformations of the muscle due to neighboring segment activations. However, the physics-based simulation employed passive deformations, which are discussed in later sections.

Description of 4-bar Mechanism and Mathematical Model Derivation

The proposed model can generate protopodium motion when muscle contractions and extensions are provided. It is known as the forward model. It is constructed based on the mechanical structure illustrated in **Figure S3**. As discussed previously, the springs approximate the actual muscles. However, spring-2 and spring-4 are hypothetical muscles used to maintain the 4-bar mechanism. The spring-2 is fixed to maintain vertical orientation. The other actuators can rotate and deform freely. The coordinate transformation method is utilized to derive the motion of the protopodium (forward kinematics). The forward kinematics model describes the location of the protopodium when the lengths of springs, the angle, and the location of the joint-2 are given. The lengths of the respective springs are denoted as $\boldsymbol{l} = [l_1, l_2, l_3, l_4, l_5]$ and the orientation of the VL as $\theta$. The $l_5$ is a hypothetical spring used in the kinematic equation that does not represent any muscles or structure of the mechanical system. It is used to derive the kinematic model by connecting two triangles in the 4-bar mechanism, which are $l_1 l_4 l_5$ and $l_2 l_3 l_5$. These two equations are obtained by following a homogeneous coordinate transformation along two directions, as illustrated in **Figure S3**. The frame $f_0$ is moved along $l_1$ and $l_4$ to the protopodium joint and denoted as $f_p$ which is denoted as direction-2. Similarly, frame $f_0$ is moved along direction-1. Both equations describe the protopodium motion in terms of lengths of VO, VL, spring-2, spring-4, spring-5, and the angle. The following expressions describe the protopodium motion in the x-axis and y-axis with respect to the frame $f_0$.

$$\boldsymbol{T_1} = \boldsymbol{T_y}(-l_2) \cdot \boldsymbol{R_z}(\beta_1 - \frac{\pi}{2})^T \cdot \boldsymbol{T_x}(l_3)$$

$$\boldsymbol{T_1} = \begin{bmatrix} \sin(\beta_1) & -\cos(\beta_1) & 0 & \sin(\beta_1)\, l_3 \\ \cos(\beta_1) & \sin(\beta_1) & 0 & \cos(\beta_1)\, l_3 - l_2 \\ 0 & 0 & 1 & 0 \\ 0 & 0 & 0 & 1 \end{bmatrix}$$

$$\boldsymbol{p_1} = \begin{bmatrix} \sin(\beta_1)\, l_3 \\ \cos(\beta_1)\, l_3 - l_2 \\ 0 \end{bmatrix}$$

The expression of the direction-2 is as follows,

$$\boldsymbol{T_2} = \boldsymbol{R_z}(\theta) \cdot \boldsymbol{T_x}(l_1) \cdot \boldsymbol{R_z}(\pi - \beta_2)^T \cdot \boldsymbol{T_x}(l_4)$$

$$\boldsymbol{T_2} = \begin{bmatrix} -\cos(\theta + \beta_2) & \sin(\theta + \beta_2) & 0 & -l_4\cos(\theta + \beta_2) + \cos(\theta)\, l_1 \\ -\sin(\theta + \beta_2) & -\cos(\theta + \beta_2) & 0 & -l_4\sin(\theta + \beta_2) + \sin(\theta)\, l_1 \\ 0 & 0 & 1 & 0 \\ 0 & 0 & 0 & 1 \end{bmatrix}$$

$$\boldsymbol{p_2} = \begin{bmatrix} -l_4\cos(\theta + \beta_2) + \cos(\theta)\, l_1 \\ -l_4\sin(\theta + \beta_2) + \sin(\theta)\, l_1 \\ 0 \end{bmatrix}$$

where, $\beta_1 = \cos^{-1}\left(\frac{l_2^2 + l_3^2 - l_5^2}{2 l_2 l_3}\right)$, and $\beta_2 = \cos^{-1}\left(\frac{l_1^2 + l_4^2 - l_5^2}{2 l_1 l_4}\right)$ are the angles illustrated in **Figure S3**. The $\boldsymbol{p_1}$ and $\boldsymbol{p_2}$ are the position vectors of the frame $f_p$ from the frame $f_0$ which designates the protopodium location. Both equations describe the motion of the protopodium joint from two different directions. A framework is proposed to obtain the lengths $\boldsymbol{l}$ and $\theta$ from the protopodium location vector, which is known as inverse kinematics. A constraint optimization approach is used to solve the inverse model. The optimization searches for values $\boldsymbol{l}$ and $\theta$ that give the actual trajectory of the protopodium but consider limits and conditions. Moreover, it does not violate the muscle contraction and extension limits obtained from the samples. The following conditions are considered during the search for muscle lengths:

- Minimize the difference between the protopodium joint motion of the $l_1 l_4 l_5$ triangle ($f_p$) and the actual protopodium motion (average sample data) to accurately describe the locomotion of the larvae ($C_1$ in the cost function $J$).
- Minimize the difference between the protopodium joint motion of $l_1 l_4 l_5$ and $l_2 l_3 l_5$ triangles to obtain realistic muscle lengths ($C_2$ in the cost function $J$). This is to solve the inverse of the kinematic chain.
- Minimize the total energy of the springs to promote the model to stay in the relaxing position as much as possible ($C_3$ in the cost function $J$).
- Smoothing factor, which generates smooth length changes ($C_4$ in the cost function $J$).

When calculating the energy, a spring constant of $1.3\ Nmm^{-1}$ is considered. Since all the muscles are assumed to be linear springs with the same spring constant, $5.2 Nmm^{-1}$ is assigned to the VL group (group of 4 muscles), and $3.9\ Nmm^{-1}$ is assigned to the VO group (group of 3 muscles). The spring-2 and spring-4 are assigned higher stiffness values, such as $6.5\ Nmm^{-1}$ due to its representation of the anatomical structure. The following is the designed cost function $J$, for the optimization which encapsulates the above conditions,

$$J = 100C_1 + 10C_2 + 30C_3 + 10C_4$$

where,

$$C_1 = \sqrt{\left(\hat{x}_p - x_{p,1}\right)^2 + \left(\hat{y}_p - y_{p,1}\right)^2}$$

$$C_2 = \sqrt{\left(x_{p,1} - x_{p,2}\right)^2 + \left(y_{p,1} - y_{p,2}\right)^2}$$

$$C_3 = \sqrt{\sum_{i=1}^{5} K_i\left(l_i - l_{i,0}\right)^2}$$

$$C_4 = \sqrt{\sum_{i=1}^{5} (l_{i,t} - l_{i,t-1})^2}$$

Here, $\hat{x}_p$ and $\hat{y}_p$ are the position coordinates of the protopodium from sample data (average of in vivo measurements of $n = 5$). Similarly, $x_{p,j}$ and $y_{p,j}$ designates the $\boldsymbol{P}_j$ vector coordinates, where $j = \{1,2\}$ representing $f_p$ of both triangles. The $K_i$ is the spring stiffness constant of *i-th* muscle. The term $(l_{i,t} - l_{i,t-1})$ computes the length change between the previous time step and the current, which indirectly determines the speed of length change. By including this term in the optimization, drastic changes in the length can be minimized, resulting in smooth length changes with time. The numerical values in the cost equation are the weights assigned to each cost to indicate the importance of each sub-cost. The optimization utilizes the limits defined by the sample data.

Data Collection

Calcium-imaging data were collected from wild-type larvae in sagittal view. Videos were imported into Kinovea (Kinovea.com) to track VO and VL tendon endpoints and measure muscle length during forward and backward crawling. Positions were recalculated relative to the left tendon of the VL group (joint 1) and rotated so that VL was parallel to the horizontal axis of the camera. VL length was used as the unit length, and other distances were measured relative to VL. Gait time was normalized to compare length changes across samples; however, actual timing was used when computing the mean and standard deviation of gait time. A MATLAB script was used to compute lengths, normalizations, and angles.

Simulation Design

We developed the physics-based robotic simulation in the Simscape Multibody environment (MathWorks) **(Figure 3O–O″)**. Each four-bar module was implemented using controllable pistons, modeled as prismatic joints, to reproduce the time-varying VO and VL length dynamics **(Figure 3P–P‴ and Figure 3Q–Q‴)**. The joints were defined as revolute joints with negligible friction and stiffness. Piston lengths were controlled using either experimentally measured muscle-length trajectories or model-predicted trajectories derived from the inverse kinematic model **(Figure S4)**. Specifically, the solid blue traces in **Figure S4** were used to drive the model-predicted simulations shown in **Figure 3P‴ and Figure 3Q‴**, whereas the dashed experimental mean traces were used to drive the in vivo-derived control simulations shown in **Figure 3P and Figure 3Q**.

Nine four-bar modules were connected to generate a segmented robotic larva approximating abdominal segments **A1–A9 (Figure 3O′–O″)**. The posterior A8/A9 region was scaled down to better approximate larval body geometry and to help balance the model. The phase shift between adjacent segmental activations was set to 20% of the gait time for forward crawling and 35% of the gait time for backward crawling. The gait time was set to 1 s to maintain consistency across simulations.

The VO and VL input signals could be independently controlled, allowing in silico mechanical silencing of either muscle group. In mechanical-silencing simulations, the silenced muscle element was not removed from the model. Instead, its active length command was blocked and the corresponding piston was treated as a passive elastic element. Thus, the silenced VO or VL element could still shorten or extend in response to forces generated by neighboring segments and then return toward its neutral length. Because passive behavior could not be dynamically enabled or disabled within a single Simscape Multibody simulation, separate otherwise identical simulations with passive pistons were run for the corresponding VO- and VL-silencing conditions.

**Intra-segmental timing analysis: VO versus other muscles (Figure S14).**

For analyses comparing VO muscles to other muscles within the same segment (**Figure S14**), individual muscle onset times were computed relative to the earliest activated muscle in that segment. Muscle onset was defined as the time point at which ΔF/F reached 20% of the total activity amplitude (peak value minus pre-peak minimum). VO onset delay was calculated as the difference between the onset time of each VO muscle and the earliest onset among all muscles in the same segment. Onset times were calculated both in real time (seconds) and in normalized time.

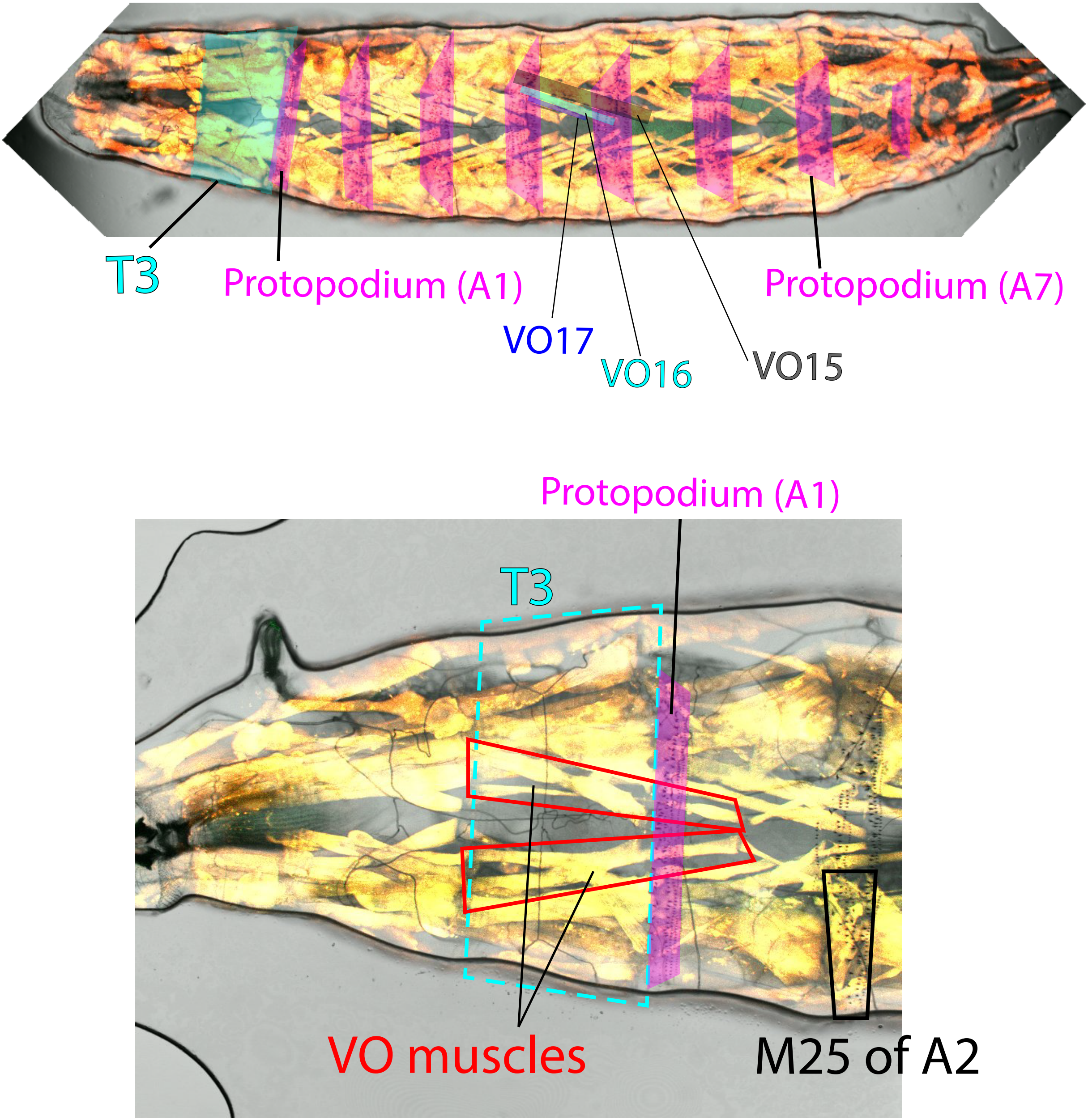

T3
Protopodium (A1)
VO17
VO16
VO15
Protopodium (A7)
Protopodium (A1)
T3
VO muscles
M25 of A2

**Figure S1. Anatomical organization of larval VO muscles and protopodia.**

**(Top)** Ventral view of a third instar (L3) larva expressing *44H10-LexA>Aop-mCherry*, showing merged AF555 (mCherry) and brightfield channels. Both left and right VO muscles are visible. VO muscles 15–17 spanning the protopodia of abdominal segments A4 and A5 are color-coded within a single hemisegment. Protopodia are outlined in magenta. The T3 segment is marked by a cyan box.

**(Bottom)** Enlarged view of segments T3 through A1. T3 contains two VO-like muscles (red boxes) that extend toward the protopodium region of segment A1. The A1 segment features a narrower protopodium composed of fewer denticle bands compared to more posterior abdominal segments. A1 also lacks muscle 25, which is present in A2 and highlighted by the black box, serving as a landmark for segment identification.

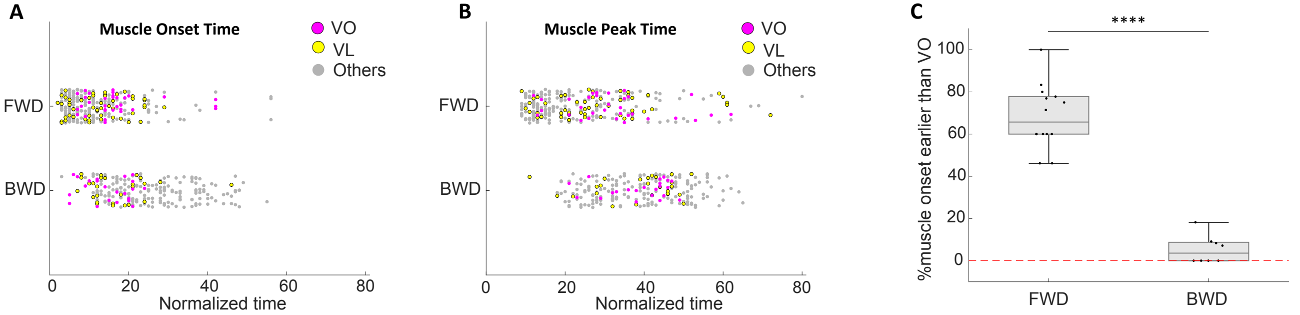
A
Muscle Onset Time
VO
VL
Others
FWD
BWD
0
20
40
60
80
Normalized time
B
Muscle Peak Time
VO
VL
Others
FWD
BWD
0
20
40
60
80
Normalized time
C
****
%muscle onset earlier than VO
100
80
60
40
20
0
FWD
BWD

**Figure S2. VO muscles are activated later in forward crawling and earlier in backward crawling.**

**A)** Scatter plot showing the onset timing of VO (magenta), VL (yellow), and all other muscles (gray) within the same segment. Onset timing is defined as the normalized time point at which GCaMP6f signal reaches 20% of its maximum amplitude. Data from multiple segments were normalized using a PCA-based method (see Materials and Methods) and pooled. In forward crawling, VO muscle onset lags behind most VL and other muscles. In backward crawling, VO muscles tend to activate earlier than other muscles. Each point represents a single muscle. Data reused from *Figure 1*.

**B)** Peak activation timing of the same muscle groups as in (A), based on the normalized time of peak GCaMP6f signal. In forward crawling, VO peak activity is generally later than other muscles, while in backward crawling, it is similar to or earlier than other muscles.

**C)** Proportion of non-VO muscles in each segment that activate earlier than the average VO onset time in that segment. In forward crawling, 40–100% of non-VO muscles activate before VO muscles, significantly higher than in backward crawling, where only 0–20% show earlier onset. Data reused from *Figure 1*. Analysis includes segments with ≥12 muscles and at least one VO muscle. Sample sizes: 13 segments from 5 animals (forward) and 8 segments from 4 animals (backward). Student’s t-test: **** $p < 0.0001$.

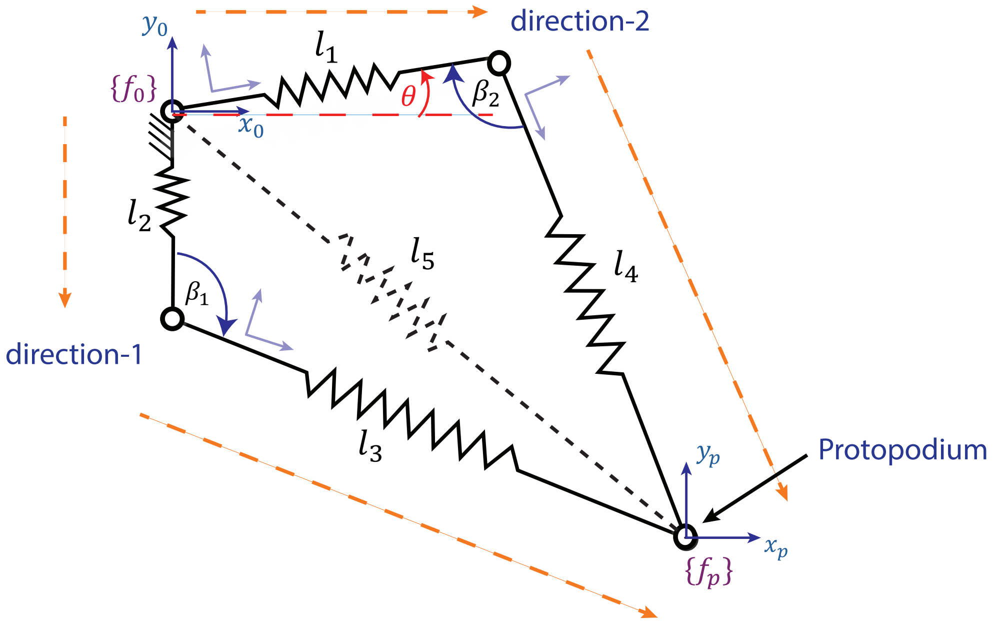

direction-2
$y_0$
$l_1$
$\{f_0\}$
$\theta$
$\beta_2$
$x_0$
$l_2$
$l_5$
$l_4$
$\beta_1$
direction-1
$l_3$
$y_p$
Protopodium
$x_p$
$\{f_p\}$

**Figure S3.** The kinematic diagram of the 4-bar mechanism. The length of the spring is denoted by $l_i$. The middle hypothetical spring (spring-5) is used to obtain the forward kinematic model of the protopodium joint. The frame-$\{f_0\}$ and the frame-$\{f_p\}$ represent the origin of the coordinate frame and the protopodium coordinate frame. The directions are depicted to indicate the kinematic chains utilized to obtain the kinematic model. The angle $\theta$ represents the VL horizontal angle, which is a parameter of the model. The $\beta_1$ and $\beta_2$ denote structural angles of the mechanism (Adapted from [1]) .

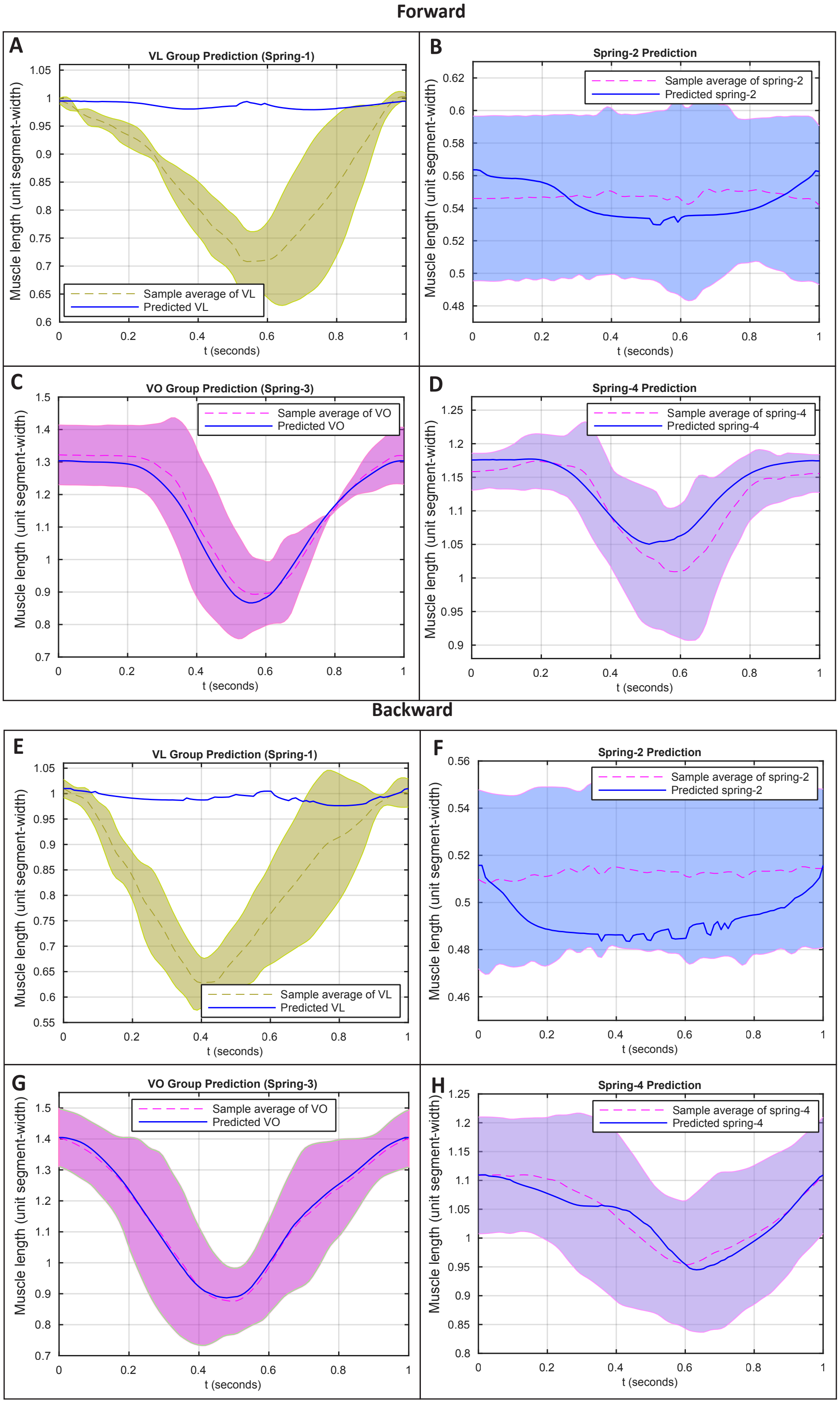

Forward
A
VL Group Prediction (Spring-1)
Sample average of VL
Predicted VL
Muscle length (unit segment-width)
t (seconds)
B
Spring-2 Prediction
Sample average of spring-2
Predicted spring-2
Muscle length (unit segment-width)
t (seconds)
C
VO Group Prediction (Spring-3)
Sample average of VO
Predicted VO
Muscle length (unit segment-width)
t (seconds)
D
Spring-4 Prediction
Sample average of spring-4
Predicted spring-4
Muscle length (unit segment-width)
t (seconds)
Backward
E
VL Group Prediction (Spring-1)
Sample average of VL
Predicted VL
Muscle length (unit segment-width)
t (seconds)
F
Spring-2 Prediction
Sample average of spring-2
Predicted spring-2
Muscle length (unit segment-width)
t (seconds)
G
VO Group Prediction (Spring-3)
Sample average of VO
Predicted VO
Muscle length (unit segment-width)
t (seconds)
H
Spring-4 Prediction
Sample average of spring-4
Predicted spring-4
Muscle length (unit segment-width)
t (seconds)

**Figure S4. Inverse kinematic model predictions of spring length dynamics underlying protopodium gait.**

The complete inverse-model predictions for the lengths of elastic elements in the four-bar mechanism are shown. Shaded regions represent the experimental mean (dashed line) ± s.d. across $n$ = 5 larvae, and solid blue lines indicate model-predicted length trajectories. **(A–D)** Forward crawling. **(A)** Predicted VL length dynamics, spring-1, required to reproduce the measured forward protopodium path. Predicted VL length changes are smaller than those observed in vivo. **(B)** Predicted length of spring-2, a hypothetical elastic element used to maintain kinematic closure. **(C)** Predicted VO length dynamics, spring-3, compared with experimental measurements. The similarity between predicted and measured VO dynamics supports a major role for VO shortening in the modeled protopodium gait.

**(D)** Predicted length of spring-4 (an auxiliary element that maintains four-bar geometry) remains within the experimentally observed range, further supporting the stability and plausibility of the inverse kinematic solution. **(E–H)** Backward crawling. **(E)** Predicted VL length dynamics required to reproduce the measured backward protopodium path. As in forward crawling, the predicted VL length changes are smaller than those measured in vivo. **(F)** Predicted length of spring-2. **(G)** Predicted VO length dynamics compared with experimental measurements. Predicted VO dynamics closely follow measured VO length changes during backward crawling. **(H)** Predicted length of spring-4. Together, these inverse-model predictions support a major role for VO length dynamics in generating the modeled forward and backward protopodium paths, whereas large VL shortening is not required under the kinematic constraints of the four-bar model.

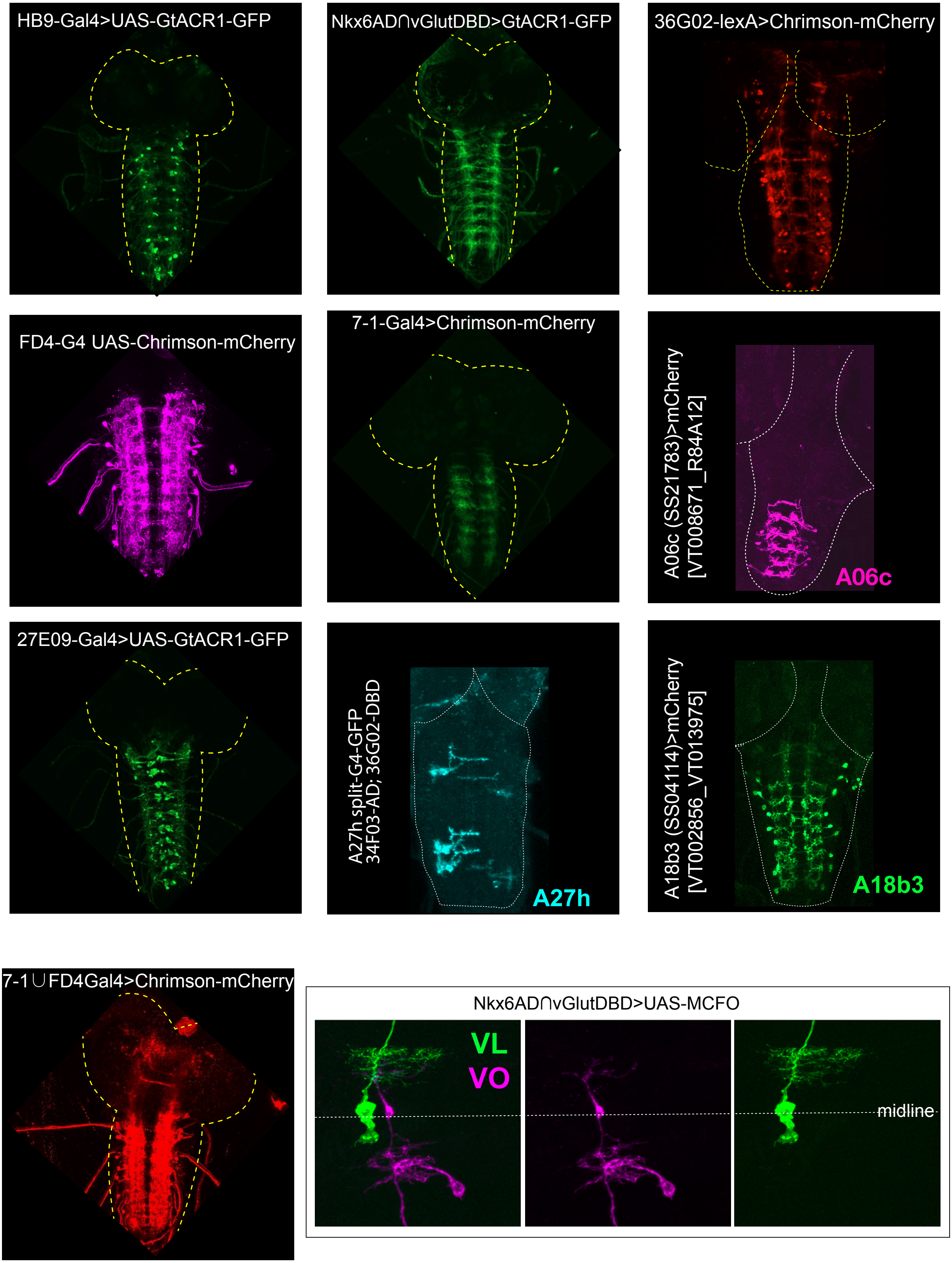
HB9-Gal4>UAS-GtACR1-GFP
Nkx6AD∩vGlutDBD>GtACR1-GFP
36G02-lexA>Chrimson-mCherry
FD4-G4 UAS-Chrimson-mCherry
7-1-Gal4>Chrimson-mCherry
A06c (SS21783)>mCherry
[VT008671_R84A12]
A06c
27E09-Gal4>UAS-GtACR1-GFP
A27h split-G4-GFP
34F03-AD; 36G02-DBD
A27h
A18b3 (SS04114)>mCherry
[VT002856_VT013975]
A18b3
7-1∪FD4Gal4>Chrimson-mCherry
Nkx6AD∩vGlutDBD>UAS-MCFO
VL
VO
midline

**Figure S5. CNS expression patterns of Gal4, LexA, and split-Gal4 driver lines used for MN and PMN manipulations.** Representative z-projections of isolated L3 larval CNS preparations expressing *UAS-GtACR1-eGFP*, *UAS-Chrimson-mCherry*, *LexAop-Chrimson-mCherry*, or related fluorescent reporters under the indicated driver lines. Driver patterns include *HB9-Gal4*, *vGlut-AD∩Nkx6-DBD split-Gal4*, *36G02-LexA*, *FD4-Gal4*, *7-1-Gal4*, *27E09-Gal4*, *A06c split-Gal4, A27h split-Gal4, A18b3 split-Gal4*, and the *7-1∪FD4-Gal4* driver. Yellow or white dashed outlines mark the larval CNS. Bottom-right panels show representative MCFO labeling from *vGlut-AD∩Nkx6-DBD split-Gal4*, illustrating labeled VO and VL MN morphologies.

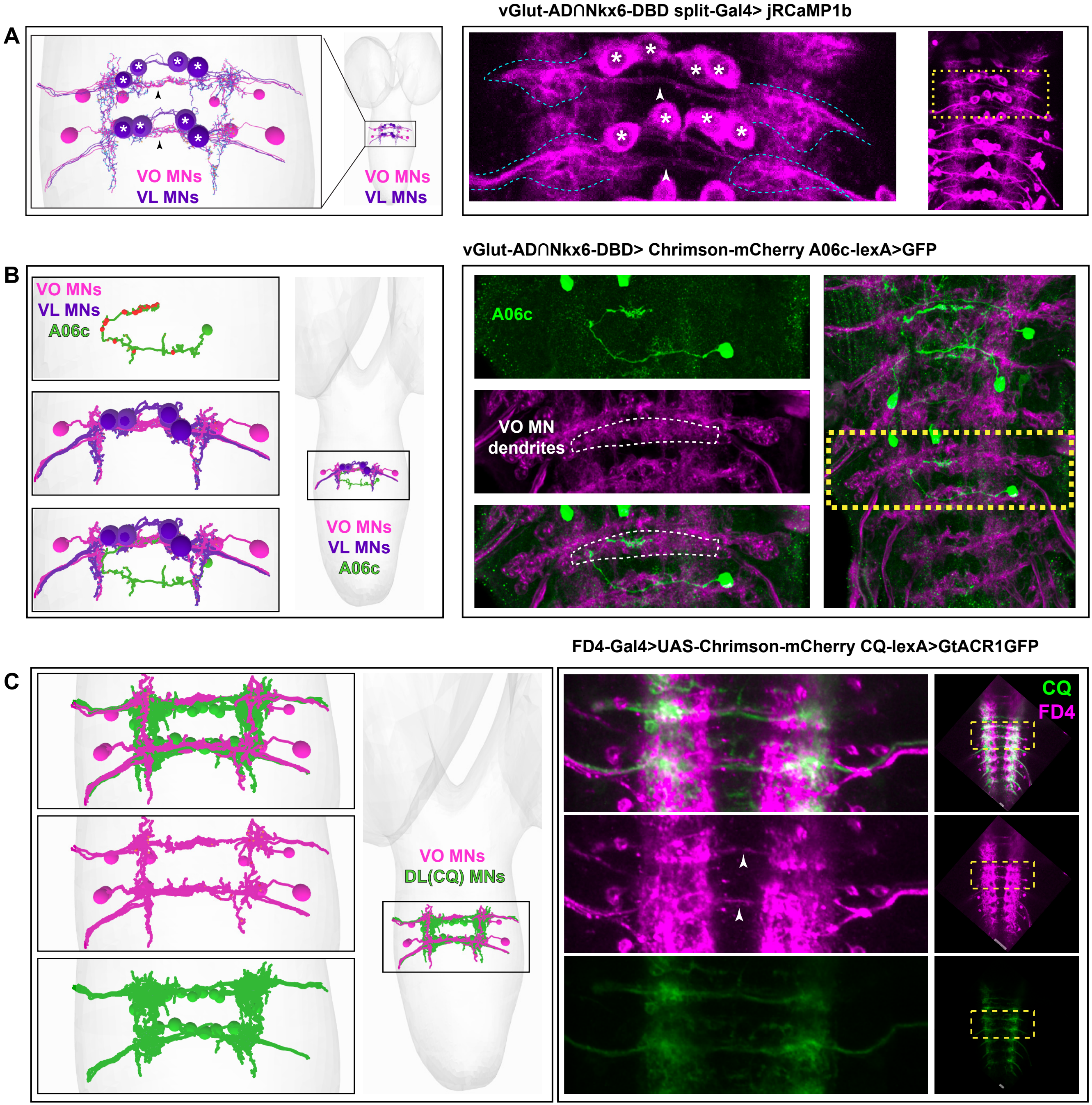
A
VO MNs
VL MNs
VO MNs
VL MNs
vGlut-AD∩Nkx6-DBD split-Gal4> jRCaMP1b
B
VO MNs
VL MNs
A06c
VO MNs
VL MNs
A06c
vGlut-AD∩Nkx6-DBD> Chrimson-mCherry A06c-lexA>GFP
A06c
VO MN
dendrites
C
VO MNs
DL(CQ) MNs
FD4-Gal4>UAS-Chrimson-mCherry CQ-lexA>GtACR1GFP
CQ
FD4

**Figure S6. EM reconstructions and confocal imaging confirm VO/VL MN dendritic organization and driver expression patterns used for calcium imaging.**

**(A)** Comparison of EM-reconstructed MN morphology and confocal fluorescence imaging for MNs targeted by *vGlut-AD∩Nkx6-DBD split-Gal4*. Left: EM reconstruction of VO and VL MNs targeted by the driver. VO MNs are shown in magenta and VL MNs in purple. Black arrowheads indicate VO MN dendrites, and asterisks mark VL MN cell bodies. Right: confocal image of *vGlut-AD∩Nkx6-DBD split-Gal4>UAS-jRCaMP1b* expression in the larval CNS. Asterisks mark VL MN cell bodies, and white arrowheads indicate the VO MN dendritic region, which is distinguishable from the labeled VL MN population.

**(B)** Relationship between A06c projections and VO/VL MN dendritic regions. Left: EM reconstruction of a single A06c neuron together with VL MNs and VO MNs. A06c is shown in green, VO MNs in magenta, and VL MNs in purple. Right: confocal image showing A06c in green and VO/VL MNs in magenta. The boxed region highlights the area where A06c axonal projections overlap the region containing VO MN dendrites, supporting correspondence between EM-defined connectivity and in vivo light-microscopy expression patterns.

**(C)** Comparison of CQ-LexA-labeled reference MNs and FD4-Gal4-labeled VO MNs. Left: EM reconstructions showing CQ reference MNs and VO MNs. Right: confocal images showing *CQ-LexA>GtACR1-GFP* and *FD4-Gal4>UAS-Chrimson-mCherry* expression in the larval CNS. VO MN dendrites cross the midline in vivo (white arrowheads), matching the EM reconstruction and supporting identification of the VO MN dendritic region used for calcium-imaging analyses. Together, these comparisons cross-validate the MN driver lines and confirm that the relevant dendritic regions identified in EM reconstructions can be distinguished in confocal recordings.

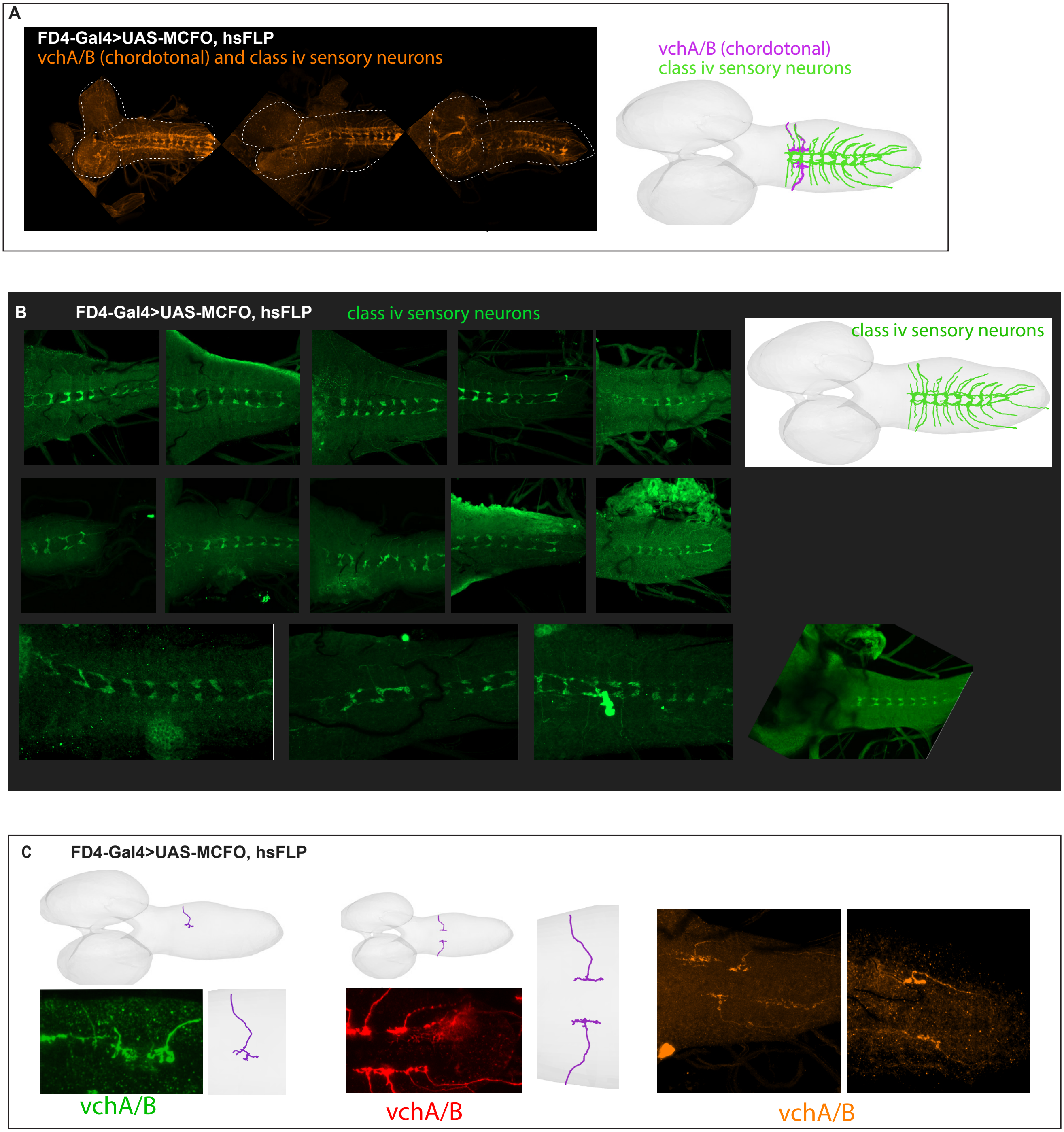
A
FD4-Gal4>UAS-MCFO, hsFLP
vchA/B (chordotonal) and class iv sensory neurons
vchA/B (chordotonal)
class iv sensory neurons
B
FD4-Gal4>UAS-MCFO, hsFLP
class iv sensory neurons
class iv sensory neurons
C
FD4-Gal4>UAS-MCFO, hsFLP
vchA/B
vchA/B
vchA/B

**Figure S7. MCFO analysis identifies FD4-Gal4-labeled sensory neurons in the CNS.**

**(A)** Representative CNS MCFO examples showing *FD4-Gal4*-labeled sensory neurons corresponding to vchA/B chordotonal neurons and class IV multidendritic sensory neurons. EM reconstructions at right show vchA/B and class IV sensory neurons used as ground truth to identify the flipped-out neurons. **(B)** Additional CNS MCFO examples showing *FD4-Gal4* labeling of class IV multidendritic sensory neurons across multiple preparations. The EM reconstruction at right illustrates the characteristic morphology and position of class IV sensory neurons used for identification. **(C)** CNS MCFO examples showing *FD4-Gal4* labeling of vchA/B chordotonal neurons. EM reconstructions are shown alongside representative flipped-out neurons to support neuronal identification. Genotype: *FD4-Gal4>UAS-MCFO, hsFLP*.

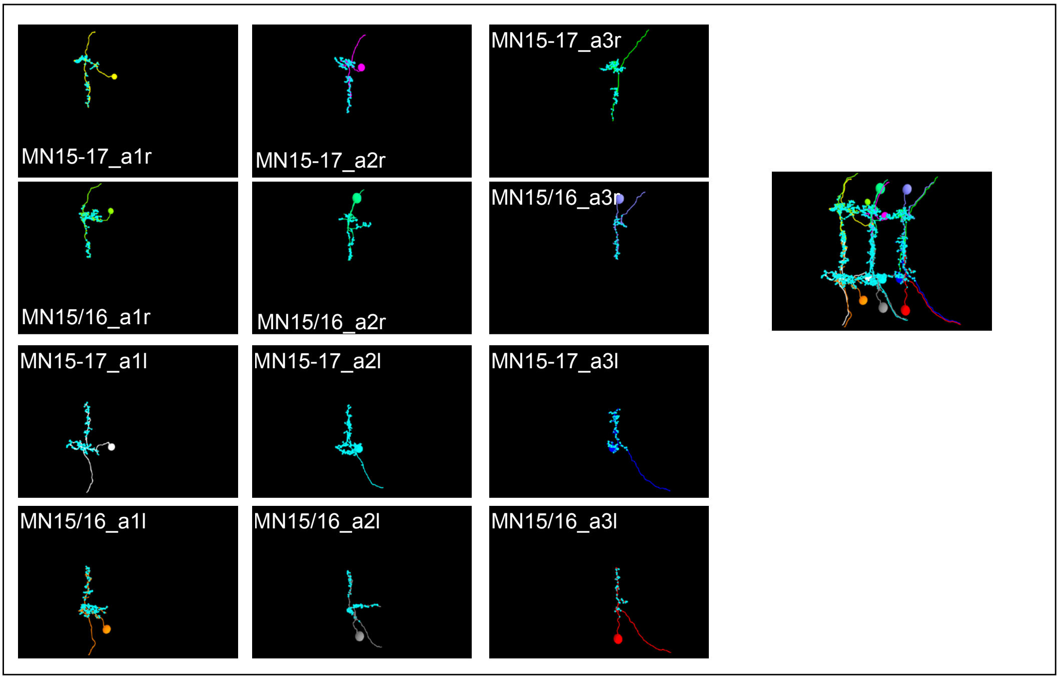
MN15-17_a1r
MN15-17_a2r
MN15-17_a3r
MN15/16_a1r
MN15/16_a2r
MN15/16_a3r
MN15-17_a1l
MN15-17_a2l
MN15-17_a3l
MN15/16_a1l
MN15/16_a2l
MN15/16_a3l

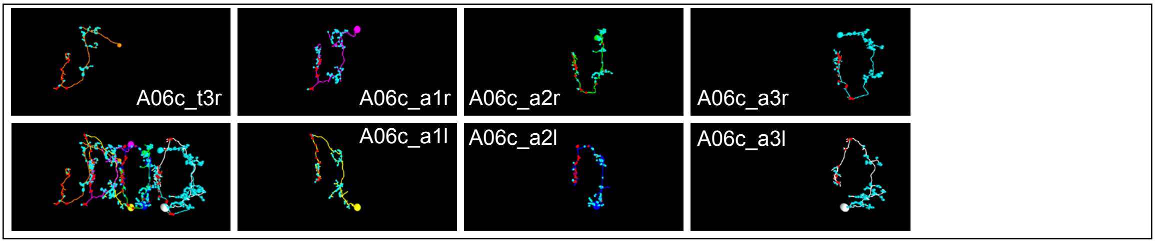
A06c_t3r
A06c_a1r
A06c_a2r
A06c_a3r
A06c_a1l
A06c_a2l
A06c_a3l

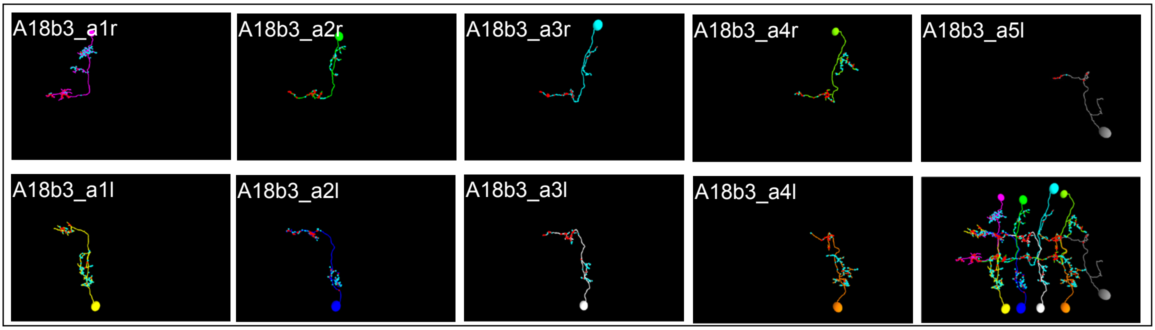
A18b3_a1r
A18b3_a2r
A18b3_a3r
A18b3_a4r
A18b3_a5l
A18b3_a1l
A18b3_a2l
A18b3_a3l
A18b3_a4l

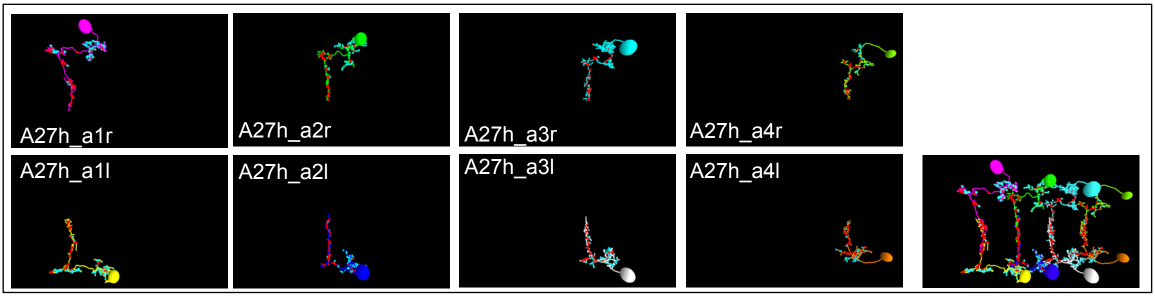
A27h_a1r
A27h_a2r
A27h_a3r
A27h_a4r
A27h_a1l
A27h_a2l
A27h_a3l
A27h_a4l

**Figure S8. Serial EM reconstruction** of VO motor neurons (MNs) and their upstream premotor partners A06c and A27h, along with the intersegmentally connected PMN A18b3, across multiple abdominal VNC segments.

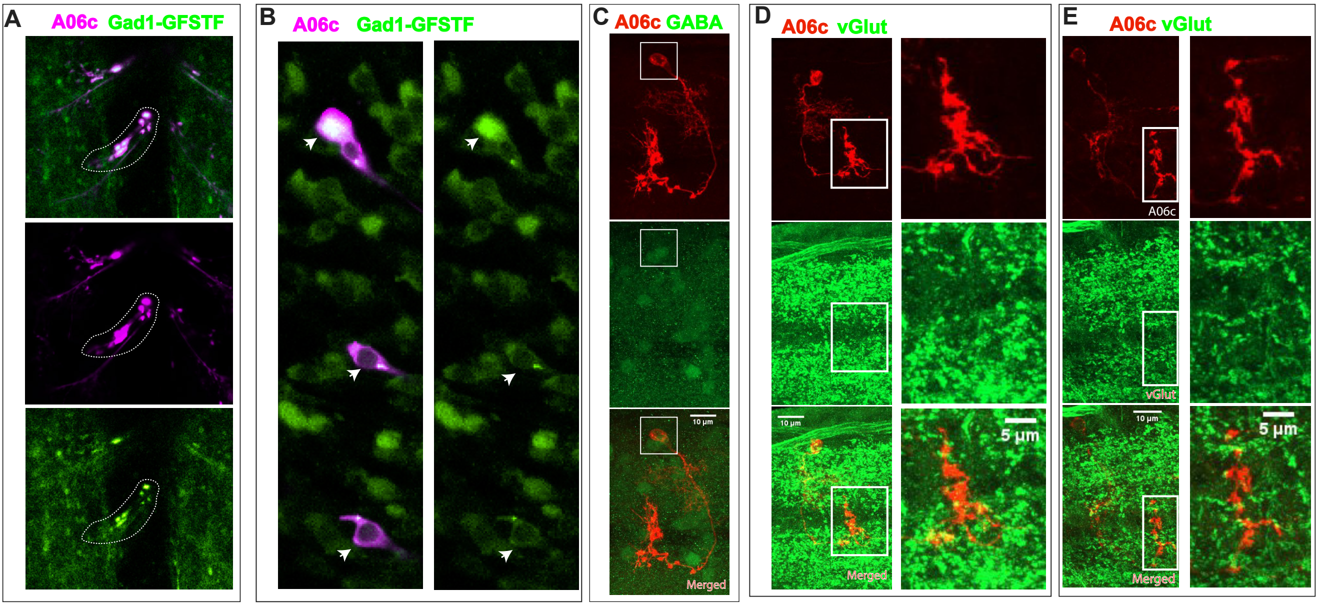
A
A06c Gad1-GFSTF
B
A06c Gad1-GFSTF
C
A06c GABA
10 µm
Merged
D
A06c vGlut
10 µm
5 µm
Merged
E
A06c vGlut
A06c
vGlut
10 µm
5 µm
Merged

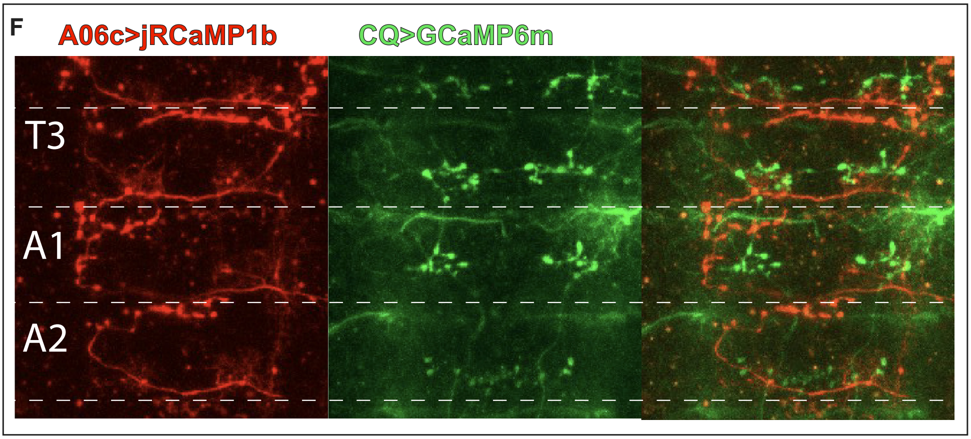
F
A06c>jRCaMP1b
CQ>GCaMP6m
T3
A1
A2

**Figure S9. A06c is a GABAergic inhibitory neuron but not glutamatergic.**
**(A–B)** Immunohistochemical staining showing an individual A06c neuron visualized by A06c split-Gal4>mCherry together with the *Gad1-GFSTF* reporter. Both the A06c cell body and axon terminals robustly express *Gad1-GFSTF*. **(C)** Co-immunostaining for GABA shows clear overlap with the A06c cell body, confirming its GABAergic identity.
**(D–E)** Immunostaining for vGlut does not overlap with A06c axon terminals or soma, indicating that A06c is not glutamatergic. **(F)** Expression pattern of A06c visualized by *jRCaMP1b* in the thoracic (T3) and abdominal (A1 and A2) segments. An off-target neuron labeled by *CQ>GCaMP6m* is shown as a segmental landmark.

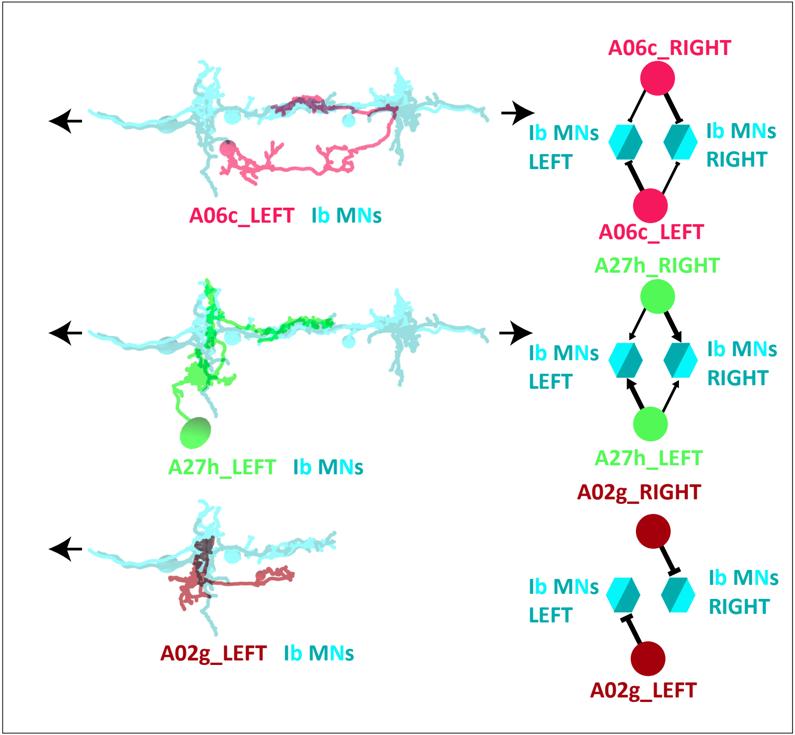

A06c_RIGHT
Ib MNs LEFT
Ib MNs RIGHT
A06c_LEFT
A06c_LEFT Ib MNs
A27h_RIGHT
Ib MNs LEFT
Ib MNs RIGHT
A27h_LEFT
A27h_LEFT Ib MNs
A02g_RIGHT
Ib MNs LEFT
Ib MNs RIGHT
A02g_LEFT
A02g_LEFT Ib MNs

**Figure S10. A06c and A27h show bilateral connectivity patterns with MNs**; a given PMN in the left or right hemisegment connects to MNs in both left and right hemisegments. In contrast, A02g is an example of a unilateral PMN, in which a PMN in the left or right hemisegment connects to MNs in only the corresponding hemisegment.

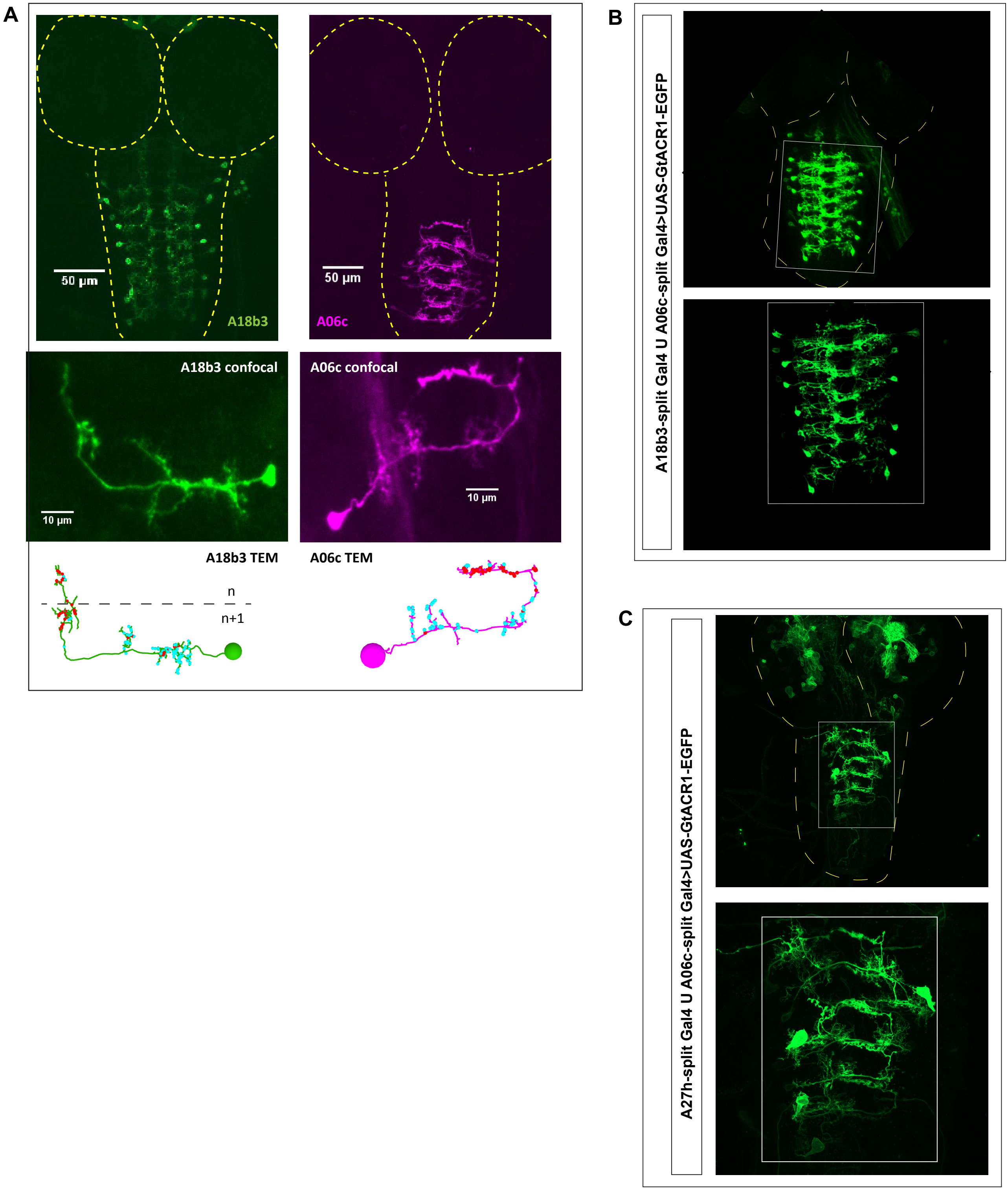

A
50 µm
A18b3
50 µm
A06c
A18b3 confocal
10 µm
A06c confocal
10 µm
A18b3 TEM
A06c TEM
n
n+1
B
A18b3-split Gal4 U A06c-split Gal4>UAS-GtACR1-EGFP
C
A27h-split Gal4 U A06c-split Gal4>UAS-GtACR1-EGFP

**Figure S11. Morphology and expression patterns of A18b3, A06c, and dual split-Gal4 combinations used for PMN silencing experiments. (A)** Confocal and TEM-based comparison of A18b3 and A06c morphology. Top row: expression patterns of the A18b3 split-Gal4 and A06c split-Gal4 lines in the L3 CNS. Middle row: confocal z-stack projections of single A18b3 or A06c neurons labeled using MCFO. Genotypes: *57c10-Flp, A18b3 split-Gal4>UAS-MCFO* or *57c10-Flp, A06c split-Gal4>UAS-MCFO*. Bottom row: A18b3 and A06c neurons reconstructed from TEM images of the L1 CNS. **(B,C)** Expression patterns of dual split-Gal4 combinations used for dual PMN silencing experiments. **(B)** *A18b3 split-Gal4 ∪ A06c split-Gal4>UAS-GtACR1-EGFP*. **(C)** *A27h split-Gal4 ∪ A06c split-Gal4>UAS-GtACR1-EGFP*. For **B** and **C**, upper panels show low-magnification views of the CNS, with the VNC boxed; lower panels show higher-magnification views of the boxed VNC region. Dashed outlines mark the brain lobes and VNC. Additional brain-lobe labeling is visible in the *A27h split-Gal4 ∪ A06c split-Gal4* combination, whereas the VNC expression pattern prominently labels the expected PMN populations. The union symbol **(∪)** denotes combined split-Gal4 expression in the same animal.

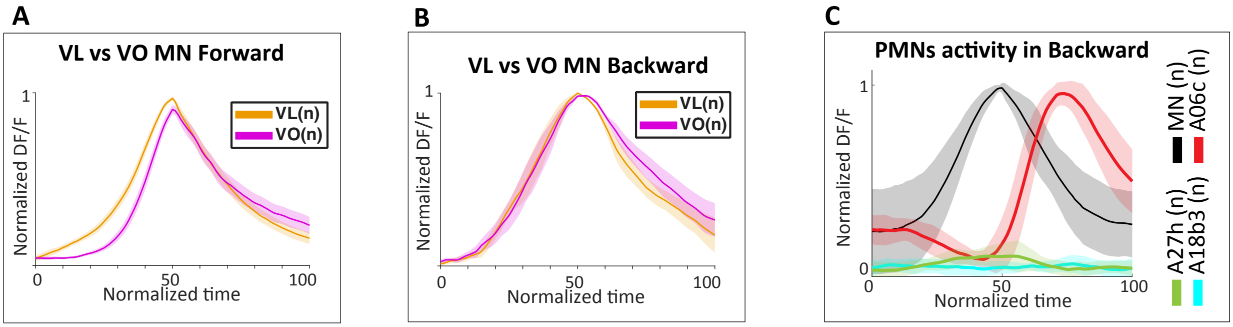
A
VL vs VO MN Forward
VL(n)
VO(n)
Normalized DF/F
Normalized time
B
VL vs VO MN Backward
VL(n)
VO(n)
Normalized DF/F
Normalized time
C
PMNs activity in Backward
MN (n)
A06c (n)
A27h (n)
A18b3 (n)
Normalized DF/F
Normalized time

**Figure S12. VO/VL MN timing and backward PMN activity during fictive locomotion in isolated larval CNS preparations. (A,B)** Phase relationship between VL and VO MN activity during fictive forward and backward locomotion. VO and VL MN activity was recorded by expressing *UAS-jRCaMP1b* with *vGlut-AD∩Nkx6-DBD split-Gal4*. Dendritic regions corresponding to VO and VL MNs were identified using EM-reconstructed MN morphology. **(A)** During fictive forward locomotion, VO MNs activate later than VL MNs. **(B)** During fictive backward locomotion, VO and VL MNs show more synchronous activity. **(C)** Phase relationships between A06c, A27h, and A18b3 PMNs and CQ-labeled reference MNs during fictive backward locomotion. PMN activity was recorded using PMN-Gal4 or PMN-split-Gal4 drivers to express *UAS-jRCaMP1b*, while *CQ-LexA* was used to express *GCaMP6m* in reference MNs. During backward locomotion, A06c activity occurs after reference MN activation, whereas A27h and A18b3 show little or no activity. Calcium traces are plotted on a normalized 0–100% fictive locomotor cycle.

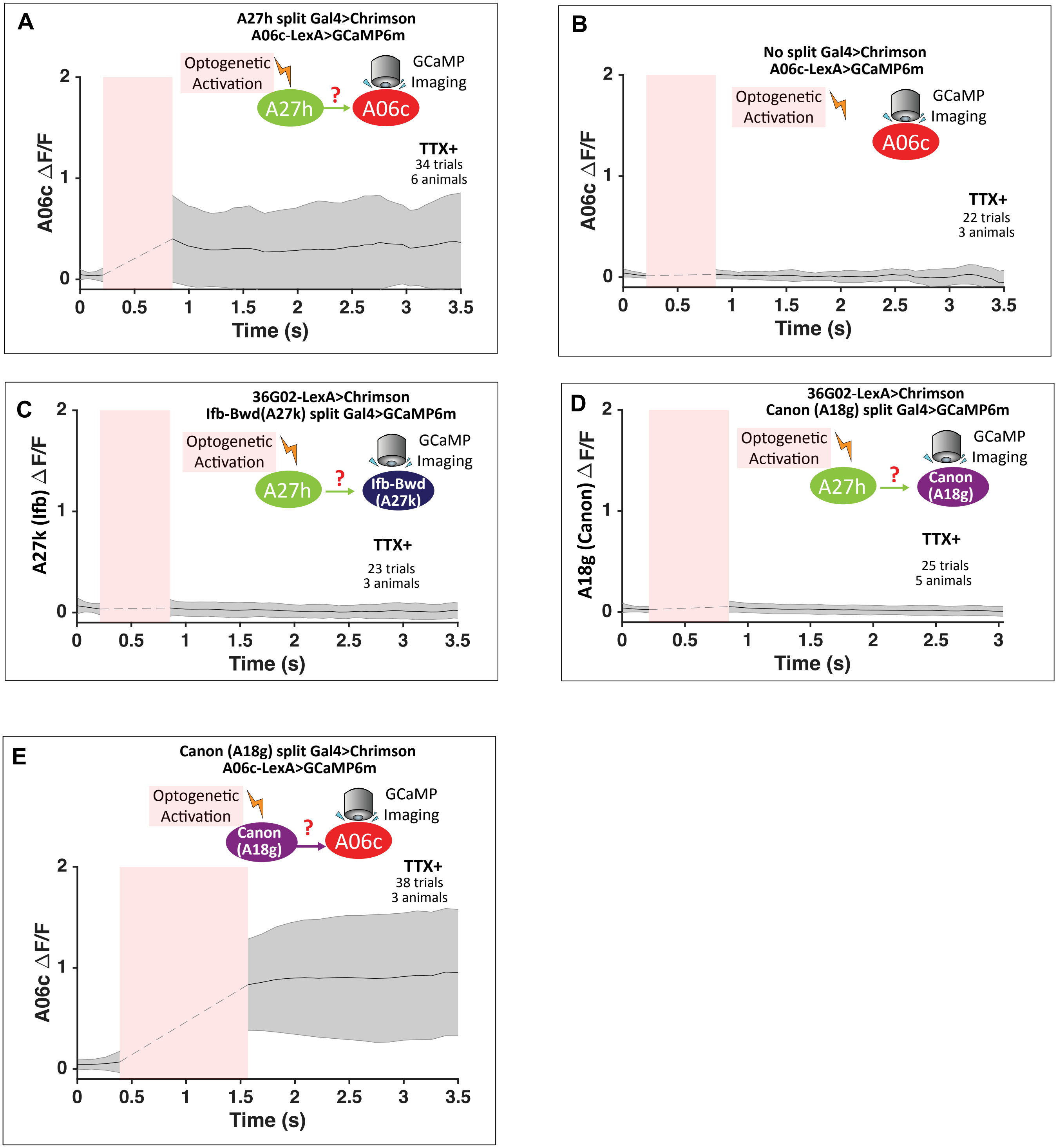
A
A27h split Gal4>Chrimson
A06c-LexA>GCaMP6m
Optogenetic Activation
GCaMP Imaging
A27h
?
A06c
TTX+
34 trials
6 animals
A06c ΔF/F
Time (s)
B
No split Gal4>Chrimson
A06c-LexA>GCaMP6m
Optogenetic Activation
GCaMP Imaging
A06c
TTX+
22 trials
3 animals
A06c ΔF/F
Time (s)
C
36G02-LexA>Chrimson
Ifb-Bwd(A27k) split Gal4>GCaMP6m
Optogenetic Activation
GCaMP Imaging
A27h
?
Ifb-Bwd (A27k)
TTX+
23 trials
3 animals
A27k (Ifb) ΔF/F
Time (s)
D
36G02-LexA>Chrimson
Canon (A18g) split Gal4>GCaMP6m
Optogenetic Activation
GCaMP Imaging
A27h
?
Canon (A18g)
TTX+
25 trials
5 animals
A18g (Canon) ΔF/F
Time (s)
E
Canon (A18g) split Gal4>Chrimson
A06c-LexA>GCaMP6m
Optogenetic Activation
GCaMP Imaging
Canon (A18g)
?
A06c
TTX+
38 trials
3 animals
A06c ΔF/F
Time (s)

**Figure S13. Additional optogenetic validation and specificity controls for Chrimson-GCaMP6 experiments in isolated larval CNS preparations.**

**(A)** Optogenetic activation of A27h using an independent split-Gal4 driver while imaging calcium activity in A06c. Genotype: *A27h split-Gal4>UAS-Chrimson; A06c-LexA>GCaMP6m*. A27h activation elicited an A06c calcium response in the presence of TTX, supporting the A27h→A06c connection with an independent A27h driver. **(B)** Control animals lacking the A27h split-Gal4 driver showed little or no A06c calcium response. Genotype: *No split-Gal4>UAS-Chrimson; A06c-LexA>GCaMP6m*. **(C,D)** Specificity controls for 36G02-LexA-mediated A27h activation. In the presence of TTX, optogenetic activation of A27h did not evoke detectable calcium responses in neurons not predicted to be monosynaptically downstream of A27h. **(C)** Imaging of Ifb-Bwd/A27k. Genotype: *36G02-LexA>LexAop-Chrimson; Ifb-Bwd/A27k split-Gal4>GCaMP6m*. **(D)** Imaging of Canon/A18g. Genotype: *36G02-LexA>LexAop-Chrimson; Canon/A18g split-Gal4>GCaMP6m*. **(E)** Optogenetic activation of Canon/A18g while imaging A06c. Genotype: *Canon/A18g split-Gal4>UAS-Chrimson; A06c-LexA>GCaMP6m*. Canon/A18g activation induced an A06c calcium response in the presence of TTX, supporting a functional Canon/A18g→A06c connection. Across calcium-response traces, solid lines indicate the mean, shaded regions indicate mean ± s.d., and pink bars denote the period of optogenetic stimulation. Total numbers of trials and larvae are indicated in each panel.

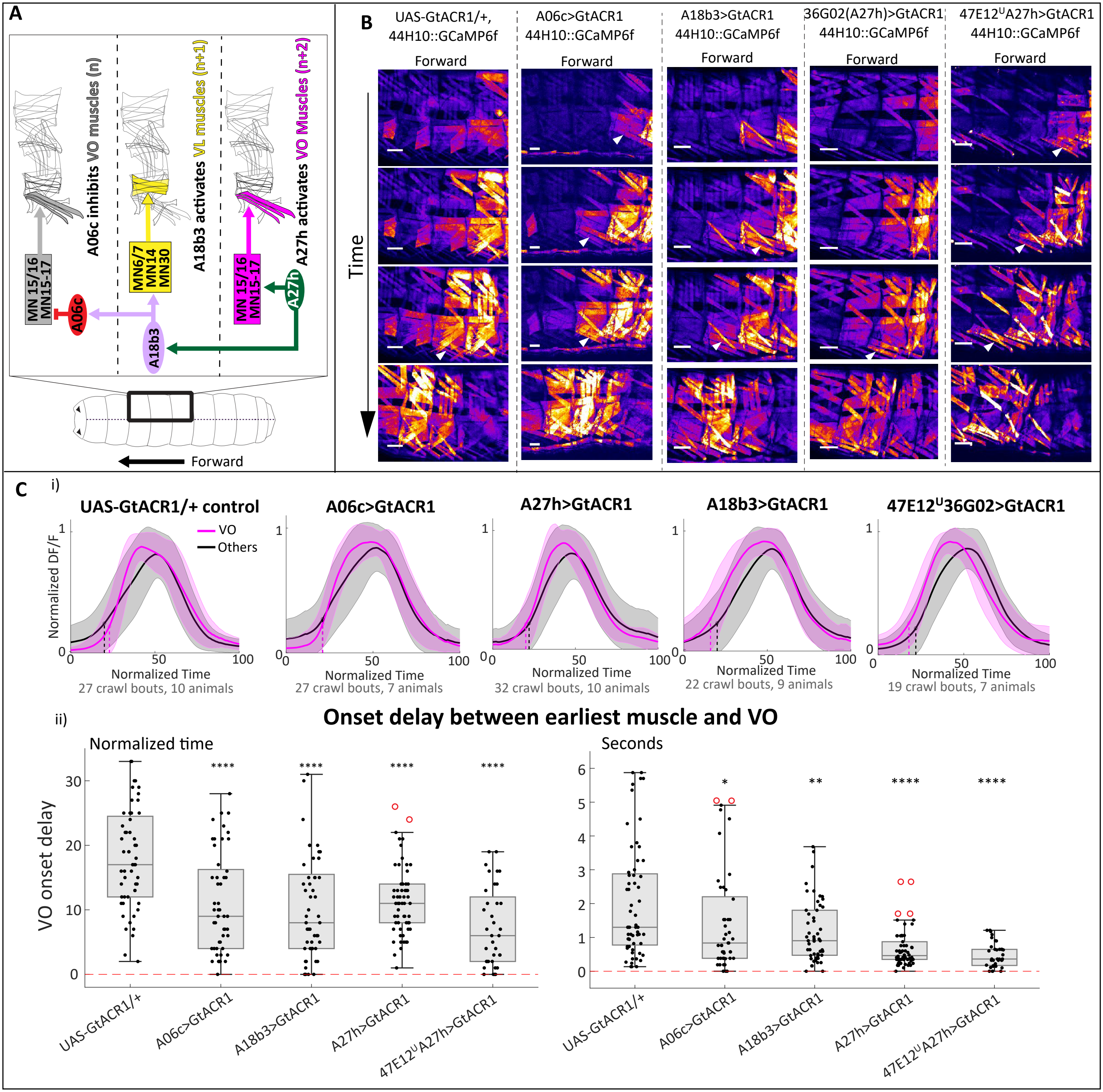
A
A06c inhibits VO muscles (n)
A18b3 activates VL muscles (n+1)
A27h activates VO Muscles (n+2)
MN 15/16
MN15-17
MN6/7
MN14
MN30
A06c
A18b3
A27h
Forward
B
UAS-GtACR1/+,
44H10::GCaMP6f
A06c>GtACR1
44H10::GCaMP6f
A18b3>GtACR1
44H10::GCaMP6f
36G02(A27h)>GtACR1
44H10::GCaMP6f
47E12UA27h>GtACR1
44H10::GCaMP6f
Time
C
i)
UAS-GtACR1/+ control
A06c>GtACR1
A27h>GtACR1
A18b3>GtACR1
47E12U36G02>GtACR1
VO
Others
Normalized DF/F
Normalized Time
27 crawl bouts, 10 animals
27 crawl bouts, 7 animals
32 crawl bouts, 10 animals
22 crawl bouts, 9 animals
19 crawl bouts, 7 animals
ii)
Onset delay between earliest muscle and VO
Normalized time
Seconds
VO onset delay
UAS-GtACR1/+
A06c>GtACR1
A18b3>GtACR1
A27h>GtACR1
47E12UA27h>GtACR1

**Figure S14. Disrupting the A27h–A18b3–A06c motif leads to premature VO activation during forward locomotion.**

**(A)** A06c–A18b3–A27h tri-segmental wiring diagram.

**(B)** Neuronal silencing was performed by expressing *UAS-GtACR1* with *A06c split-Gal4*, *A18b3 split-Gal4*, *36G02*-Gal4 *(A27h),* or the *47E12∪36G02* dual-Gal4 combination. Muscle activity was monitored using *44H10-GCaMP6f* during forward crawling in control animals (*UAS-GtACR1/+; 44H10-GCaMP6f/+*) and each PMN-silencing condition. Silencing A06c, A27h, A18b3, or both A18b3 and A27h led to premature VO activation, indicated by white arrowheads. Scale bars: 50 μm.

**(C)** Quantification of VO activation timing relative to other muscles in the same segment. **(i)** Line plots comparing VO muscles (magenta traces) with the remaining muscles in the same segment (black traces). In control animals, VO activity occurs later than the other muscles. In *A06c*, *A18b3*, *36G02*, or *47E12∪36G02>UAS-GtACR1* larvae, VO muscles activate earlier, indicated by left-shifted VO traces. **(ii)** Quantification of the phase delay between VO onset and the earliest muscle onset in the same segment. VO onset delay is reduced in *A06c*, *A18b3*, *36G02*, and *47E12∪36G02>UAS-GtACR1* larvae. Onset time was defined as the time point at which muscle activity reached 20% of its peak-normalized increase before peaking. Kruskal–Wallis test with Dunn's post hoc test comparing each experimental group with control. Asterisks denote adjusted $P$ values: *, $P < 0.05$; **, $P < 0.01$; ***, $P < 0.001$; ****, $P < 0.0001$.

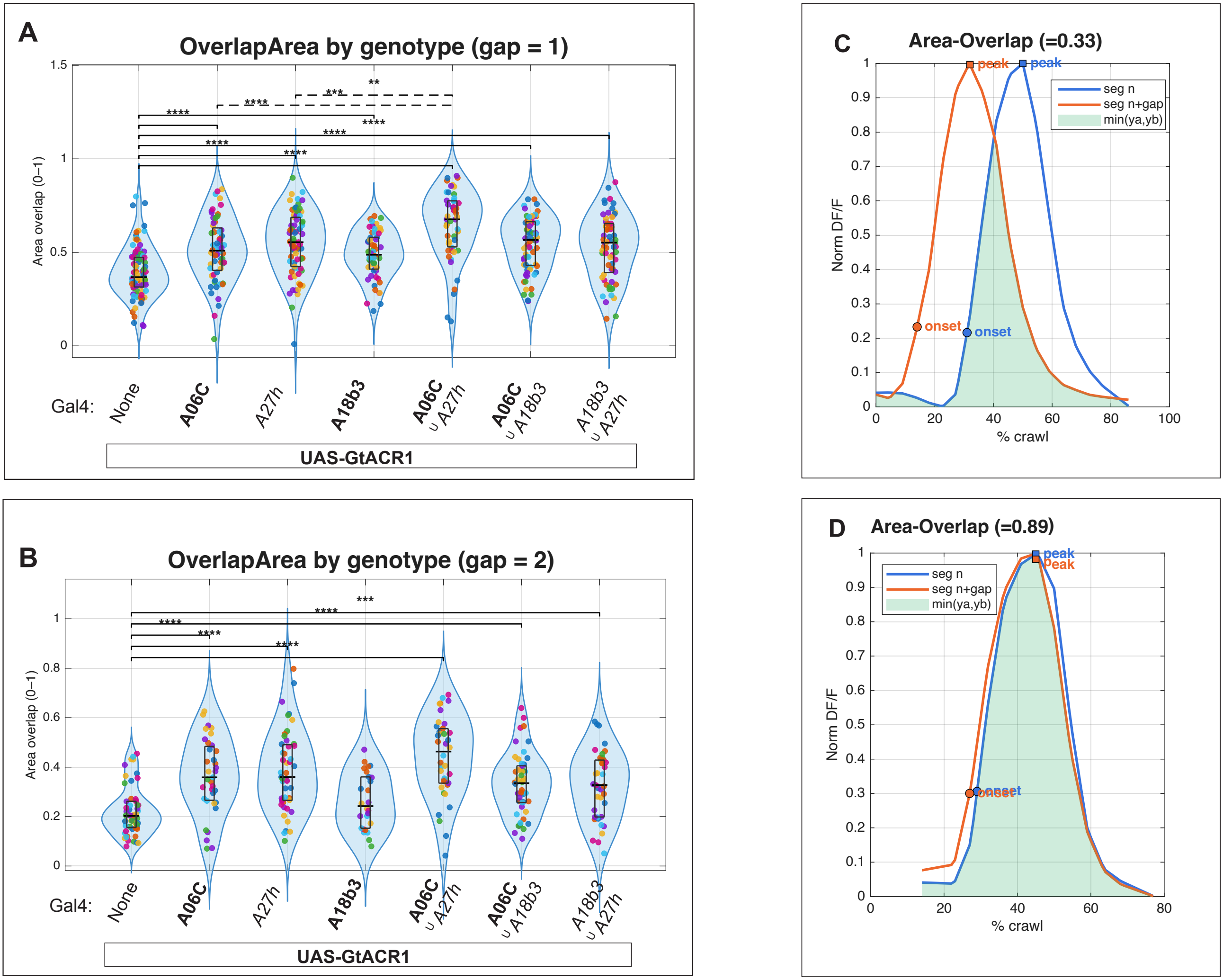
A
OverlapArea by genotype (gap = 1)
Area overlap (0–1)
Gal4:
None
A06C
A27h
A18b3
A06C ∪ A27h
A06C ∪ A18b3
A18b3 ∪ A27h
UAS-GtACR1
B
OverlapArea by genotype (gap = 2)
Area overlap (0–1)
Gal4:
None
A06C
A27h
A18b3
A06C ∪ A27h
A06C ∪ A18b3
A18b3 ∪ A27h
UAS-GtACR1
C
Area-Overlap (=0.33)
Norm DF/F
% crawl
peak
onset
seg n
seg n+gap
min(ya,yb)
D
Area-Overlap (=0.89)
Norm DF/F
% crawl
peak
onset
seg n
seg n+gap
min(ya,yb)

**Figure S15. Disrupting the A27h–A18b3–A06c premotor motif increases intersegmental overlap of VO muscle activity.**

Intersegmental coordination between VO muscles was quantified for adjacent segment pairs (gap = 1) and every-other segment pairs (gap = 2) using an amplitude-weighted overlap metric computed from paired calcium activity traces. These analyses complement the timing-based coordination and FWHM-overlap defects shown in **Figure 9** by directly assessing the extent to which intersegmental VO muscle activity temporally and quantitatively overlaps.

**(A,B)** Integrated area overlap quantifies intersegmental coordination as the integral of the point-by-point minimum of two normalized calcium traces, thereby incorporating both activity duration and amplitude, for gap = 1 **(A)** and gap = 2 **(B)**. Each dot represents one forward crawl bout–level measurement for a given segment pair, pooled across larvae. The number of larvae and analyzed crawl bout–segment pairs ranged from N = 5–15 larvae and n = 12–96 segment-pair measurements per genotype. Group differences were first assessed using Kruskal–Wallis tests; planned pairwise comparisons were performed using two-sided Wilcoxon rank-sum tests with Holm–Bonferroni correction. Solid black horizontal brackets indicate statistically significant comparisons between control and the corresponding PMN-silenced group, whereas dashed black brackets indicate statistically significant comparisons between the indicated genotype pairs. Asterisks denote adjusted $P$ values: *, $P < 0.05$; **, $P < 0.01$; ***, $P < 0.001$; ****, $P < 0.0001$. Only statistically significant comparisons are shown. PMN labels in italics indicate conventional Gal4 drivers (47E12-Gal4 for *A18b3* and 36G02-Gal4 for *A27h*), whereas boldface PMN labels in the figure indicate split-Gal4 lines.

**(C,D)** Representative examples of VO muscle calcium activity traces illustrating temporally segregated activity with low overlap **(C)** and highly synchronous activity with extensive overlap **(D)** between adjacent segments. Green shaded regions indicate the overlapping activity quantified by the area-overlap metric, defined as the point-wise minimum of the two traces. Together with the intersegmental timing and FWHM-overlap metrics quantified in **Figure 9**, these overlap-based analyses demonstrate that silencing components of the A27h–A18b3–A06c motif collapses normally staggered VO muscle activation into abnormally synchronous and overlapping bursts, revealing a complementary failure mode of intersegmental coordination during forward locomotion.

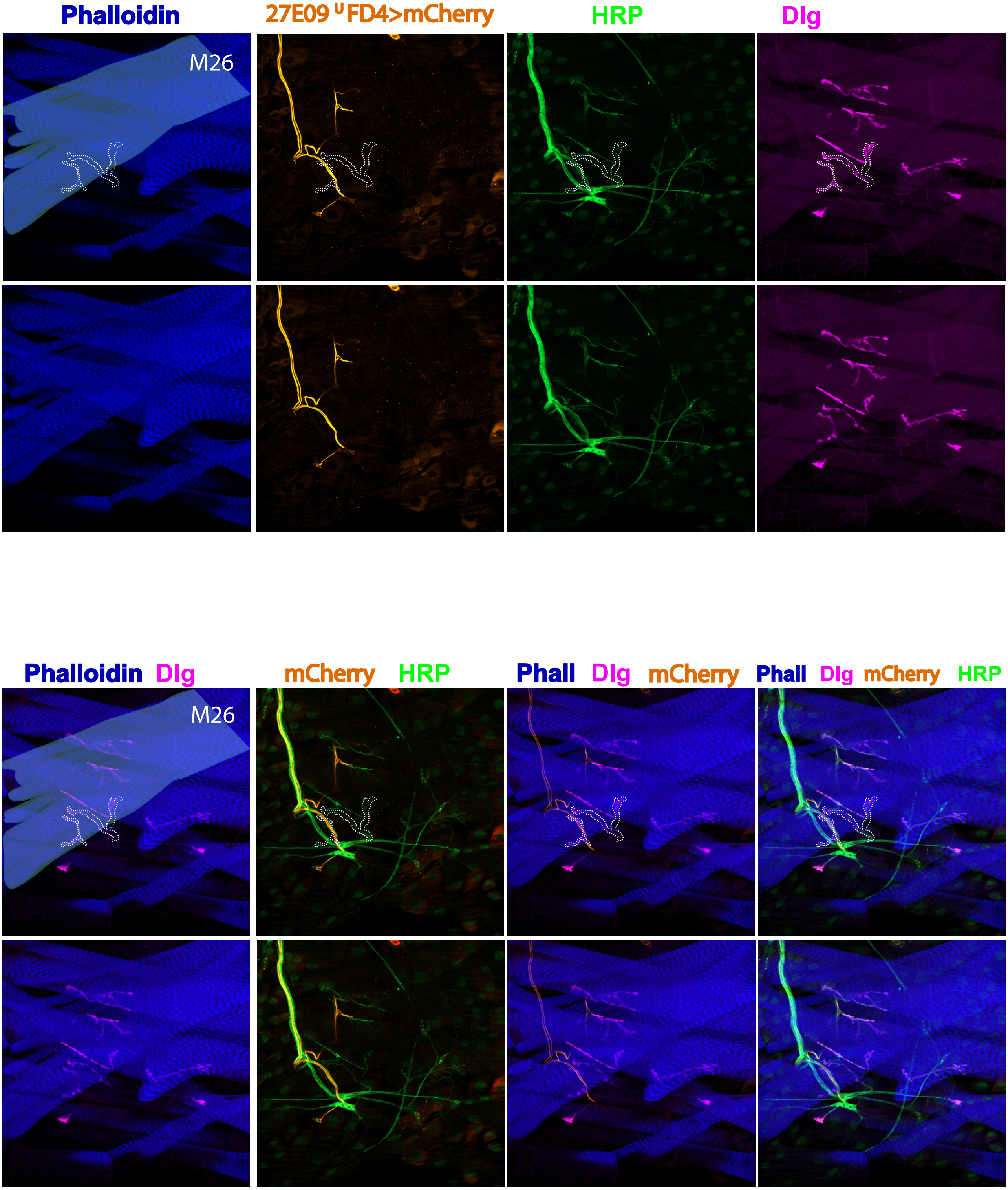
Phalloidin
27E09 U FD4>mCherry
HRP
Dlg
M26
Phalloidin Dlg
mCherry HRP
Phall Dlg mCherry
Phall Dlg mCherry HRP
M26

**Figure S16. Muscle 26 receives tonic motor innervation but not detectable phasic ISNb/d (RP5) input.** Confocal images of larval body-wall muscles showing the innervation of muscle 26 (M26). Muscle fibers are labeled with phalloidin (blue), and the boundary of M26 is indicated by cyan outlines in the leftmost panels. Motor neuron axons and presynaptic terminals are labeled with anti-HRP (green), and postsynaptic neuromuscular junctions (NMJs) are marked by Discs large (Dlg; magenta). Tonic motor neuron axons form clear presynaptic terminals on M26, accompanied by Dlg-positive postsynaptic specializations; these NMJ sites are highlighted by white dashed outlines. In contrast, the phasic ISNb/d motor neuron (RP5), visualized with *27E09(Is)-Gal4>UAS-mCherry* (orange), does not form presynaptic terminals or Dlg-positive postsynaptic sites on M26. Multiple channel combinations and replicate panels, including views without dashed outlines, are shown for comparison and clarity.

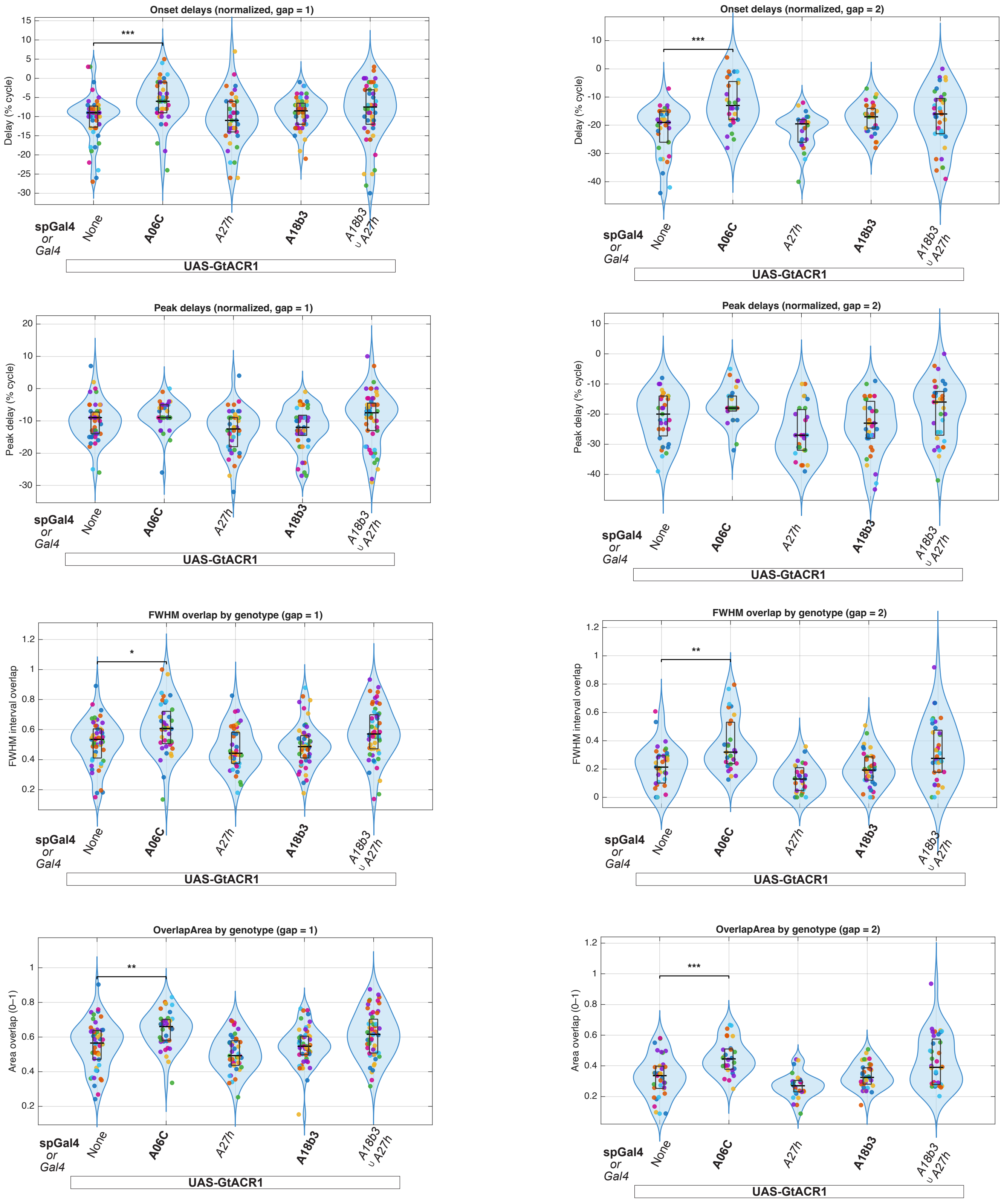

Onset delays (normalized, gap = 1)
Delay (% cycle)
***
Onset delays (normalized, gap = 2)
Delay (% cycle)
***
Peak delays (normalized, gap = 1)
Peak delay (% cycle)
Peak delays (normalized, gap = 2)
Peak delay (% cycle)
FWHM overlap by genotype (gap = 1)
FWHM interval overlap
*
FWHM overlap by genotype (gap = 2)
FWHM interval overlap
**
OverlapArea by genotype (gap = 1)
Area overlap (0–1)
**
OverlapArea by genotype (gap = 2)
Area overlap (0–1)
***
spGal4
or
Gal4
None
A06C
A27h
A18b3
A18b3 ∪ A27h
UAS-GtACR1

**Figure S17. A06c contributes to intersegmental VO coordination during backward locomotion.** VO15/16/17 calcium activity was measured during backward crawling after silencing A06c, A27h, A18b3, or A18b3∪A27h with *UAS-GtACR1*. Intersegmental coordination was quantified between adjacent segments, gap = 1, and between every other segment, gap = 2. **(A–D)** VO coordination between adjacent segments, gap = 1. **(A)** Onset delay. **(B)** Peak delay. **(C)** FWHM interval overlap. **(D)** Integrated area overlap. **(E–H)** VO coordination between every other segment, gap = 2. **(E)** Onset delay. **(F)** Peak delay. **(G)** FWHM interval overlap. **(H)** Integrated area overlap. A06c silencing reduced intersegmental onset delays and increased FWHM and area overlap during backward crawling, whereas A27h, A18b3, and A18b3∪A27h silencing had little effect on these measures. Each dot represents one crawl-bout/segment-pair measurement pooled across larvae. N = 4–6 larvae per genotype. Statistics: Kruskal–Wallis tests followed by planned Wilcoxon rank-sum comparisons with Holm–Bonferroni correction. Only significant comparisons are shown. The union symbol (∪) denotes combined PMN silencing.

**Video S1. Larval protopodia show differential heel-toe progression during forward and backward peristaltic crawling.** Video shows wild-type larval protopodia during crawling (anterior is left). During forward crawling, each protopodium enters swing phase in a heel-to-toe manner and returns to stance by the heel making contact with the surface followed by the toe. In backward crawling, this sequence is reversed. This resembles the heel-toe progression of one foot in human forward and backward walking.

**Video S2. The VO muscles are activated earlier in backward than in forward locomotion.** Muscle calcium imaging in an intact wild-type (*44H10::GCaMP6f*) animal performing forward and then backward locomotion. White arrowheads point at VO muscles as they are being activated in each segment. Compared to other muscles in the same segment, VO muscles are activated late in forward locomotion but early in backward locomotion. Anterior is left and dorsal is up.

**Video S3. VO muscles have different phase relationships with protopodia during forward and backward locomotion.** VO muscle activity and protopodium movement in wild-type larvae during forward and backward crawling were imaged simultaneously using calcium and bright-field imaging. During forward crawling, VO contraction is mostly synchronous with the swing phase of the protopodium in the next posterior segment. During backward crawling, VO contraction is mostly synchronous with the swing phase of the protopodium in the same segment. Anterior is left.

**Video S4. Mechanical model of VO/VL timing and protopodium anchoring during forward and backward crawling.**

Annotated video illustrating how behavior-specific VO/VL timing may convert centerward muscle contraction into direction-specific protopodium movement. **P n** denotes the protopodium associated with segment **n**. During forward crawling, VO/VL contraction in the active segment is proposed to fold and swing the posterior protopodium, while early VL activity in the next anterior segment opposes force transmission and helps maintain the anterior protopodium in stance. During backward crawling, this relationship is reversed: VO/VL contraction in the active segment is proposed to fold and swing the anterior protopodium, while early VO/VL activity in the next posterior segment opposes force transmission and helps maintain the posterior protopodium in stance. Red arrows indicate VO/VL contraction in the active segment. Purple arrows indicate early neighboring VL or VO/VL activity. White braces indicate increased muscle tone and resistance to stretch in the neighboring segment.

**Video S5. Robotic simulations of forward and backward protopodium gait generation.**
Physics-based robotic simulations corresponding to **Figure 3P–P‴ and Figure 3Q–Q‴**. Forward and backward locomotor simulations are shown separately. For each direction, four conditions are shown: control simulation driven by in vivo-derived VO/VL muscle dynamics, in silico mechanical silencing of VO contraction, in silico mechanical silencing of VL contraction, and simulation driven by muscle dynamics predicted by the kinematic model. In forward simulations, control in vivo-derived dynamics generated effective forward displacement, VO silencing markedly reduced displacement, VL silencing preserved displacement but produced skidding, and model-predicted dynamics generated forward displacement comparable to the VL-silenced condition. In backward simulations, control in vivo-derived dynamics generated backward displacement, whereas in silico silencing of either VO or VL reduced displacement. Simulations driven by model-predicted dynamics produced still lower backward displacement, likely reflecting features not fully captured by the current model, including passive tissue deformation, detailed substrate grip, and the mechanical effects of neighboring body-wall muscles. The robotic model is intended as a test of mechanical sufficiency rather than a complete quantitative reconstruction of larval locomotion.

**Video S6. Activation of VO MNs causes ventral C-bending in intact larvae.**
The dorsal trachea and ventral protopodia were labeled to distinguish dorsal and ventral bending. Activation of dorsal MNs using *CQ-Gal4* with *ChAT-Gal80* caused larvae to curve toward the dorsal side. Activation of VO-MN15–17-containing driver combinations, including *FD4-Gal4* with *ChAT-Gal80* and the *7-1∪FD4-Gal4* union driver, caused larvae to contract and curve toward the ventral side. A similar ventral-bending response was observed following activation of both VO and VL MNs using *vGlut-AD∩Nkx6-DBD split-Gal4*. Activation of VL/DO MNs using *HB9-Gal4* primarily caused body shortening.

**Video S7. VO muscle activity is sufficient to drive protopodium folding.**
MN-Gal4 lines targeting different sets of tonic Ib MNs were optogenetically activated with *UAS-Chrimson-mCherry* while muscle activity (*44H10-GCaMP6f*) and protopodium movement (bright

field) were recorded. The 488-nm laser used for GCaMP imaging also activated Chrimson. Activation of VO-MN15–17 using the *7-1∪FD4-Gal4* union driver, either alone or together with VL MN activation using *vGlut-AD∩Nkx6-DBD split-Gal4*, induced protopodium folding. By contrast, activation of VL/DO MNs alone using *HB9-Gal4* produced weaker protopodium folding. Short colored lines mark protopodium width.

**Video S8. VO and VL motor output are required for efficient forward crawling.**
Silencing MN inputs to VO, VL, or both VO and VL muscles produces distinct forward-crawling defects. In control animals, forward crawling is powered by propagating muscle contractions, with protopodia serving as ground-interaction points. Optogenetic silencing of VO-MN15–17 together with phasic Is MNs (*FD4∪7-1∪27E09>UAS-GtACR1*) caused slow propagation, short strides, protopodium folding defects, and occasional backward sliding. Silencing VL MNs together with phasic Is MNs (*HB9∪27E09>UAS-GtACR1*) caused slow propagation and short strides. Silencing tonic VO and VL MNs together with phasic Is MNs (*vGlut-AD∩Nkx6-DBD∪27E09>UAS-GtACR1*) caused slow crawling, short strides, protopodium folding defects, and occasional backward sliding.

**Video S9. VO muscles are necessary for normal protopodium movement.**
Video clips show dual-channel muscle calcium and brightfield protopodium imaging in control larvae and MN GtACR1 optogenetic-silencing conditions, with diagrams indicating the target muscles of each MN-Gal4 line. In *HB9>UAS-GtACR1* larvae **(VL-MN silencing)**, VO muscles retained GCaMP activity, and protopodia showed relatively normal folding with reduced forward displacement. In *FD4∪7-1∪27E09>UAS-GtACR1* larvae **(VO plus phasic Is MN silencing)**, VO muscle activity was reduced, and protopodia showed shorter forward displacement. In *vGlut-AD∩Nkx6-DBD>UAS-GtACR1* larvae **(VO and VL MN silencing)**, VO muscle activity was strongly reduced or absent, and protopodium folding and forward displacement were largely eliminated. Silencing phasic Is MNs (*27E09-Gal4*) in combination with tonic MN-Gal4 lines intensified the effects of tonic MN silencing.

**Video S10. A06c has a bi-modal firing pattern, while A18b3 and A27h are forward-dedicated.** Neuronal activity was recorded by dual-color calcium imaging in isolated larval CNS preparations performing fictive backward and then forward locomotion. Anterior is left. The activity of reference MNs expressing GCaMP6m (*CQ-LexA>GCaMP6m*), PMNs (*A06c split-Gal4, A18b3 split-Gal4 or 36G02-Gal4>jRCaMP1b*) expressing jRCaMP1b, and a merged image are shown. White arrowheads point at MNs and A06c PMN as they are actively firing in a single segment. During forward locomotion, A06c PMN fires twice, once before and once after MNs in its corresponding segment. During backward locomotion, A06c PMN fires only once after MNs in its corresponding segment. During forward locomotion, A18b3 PMN fires mostly in synchrony with MNs in the same segment. During backward locomotion, A18b3 PMN has no activity.

During forward locomotion, A27h PMN fires later than MNs in the same segment. During backward locomotion, A27h PMN has no activity. In addition to A27h, *36G02-Gal4* line hits another neuron known as A03g and M neuron (Kohsaka et al. 2019 Nat Comm., Zeng et al. 2021 Curr Biol.). The data from this animal has been previously published in another paper published by the same author (Zarin et al. 2019 eLife).

**Video S11. Disrupting the function of A06c-A18b3-A27h microcircuit leads to premature VO muscle activation in forward locomotion.** Video shows muscle calcium imaging in a control (*UAS-GtACR1/+*) animal performing forward locomotion and in animals in which A06c, A18b3, A27h, or both A18b3 and A27h were silenced using *A06c split-Gal4, A18b3 split-Gal4, 36G02-Gal4, or 47E12$^{\cup}$36G02-Gal4 driven GtACR1*. White arrowheads point at VO muscles as they are being activated in each segment. Anterior is left and dorsal is up. In any of the PMN loss-of-function animals, the VO muscles are activated abnormally early compared to other muscles in the same segment during forward locomotion (see Figure 9 and Figure S14).

**Video S12. Disruption of the A27h–A18b3–A06c premotor motif alters VO muscle coordination during forward crawling.**
Representative muscle calcium imaging showing VO muscle activity during forward locomotion in control larvae and following optogenetic silencing of individual or paired premotor neurons, illustrating disrupted intersegmental and intrasegmental coordination (see Figure 9).

**Video S13. Disrupting the A06c–A18b3–A27h circuit causes premature VO activity, reduced protopodium folding, shortened stride length, and defective heel-to-toe progression during forward crawling.**
In control larvae (*44H10-GCaMP6f; UAS-GtACR1/+*), VO activity progresses sequentially, with VO contraction occurring one segment at a time. Protopodia remain anchored as posterior segments contract and show normal heel-to-toe progression during folding. In larvae in which one or two PMNs from the A06c–A18b3–A27h circuit are silenced with GtACR1, VO muscles in

three consecutive segments frequently show synchronized or strongly overlapping activity, suggesting premature VO recruitment. In these larvae, protopodia are frequently dragged backward by posterior segments, consistent with weakened anchoring, and show reduced folding amplitude, shortened stride length, and abnormal heel/toe progression, including toe-to-heel folding or near-synchronous toe and heel folding. Color-coded shaded regions mark protopodia in different segments, and dashed lines of the corresponding color indicate the initial protopodium positions. Arrowheads mark active VO muscles. Black arrows point to the toe, defined as the anterior-most denticle band, or the heel, defined as the posterior-most denticle band, to indicate abnormal heel/toe progression during folding.

**Video S14. Disrupting the A27h–A18b3–A06c premotor motif reduces forward peristalsis efficiency.**

Video shows a control animal (*UAS-GtACR1/+*, left) and an *A18b3(47E12) ∪ A27h>UAS-GtACR1* animal (right) crawling forward in straight tracks in an agarose gel. The two animals have similar peristalsis frequency, but the *A18b3(47E12) ∪ A27h>UAS-GtACR1* animal moves a shorter distance during each peristaltic cycle than the control. Each grid square is 5 mm.

**Table S1. PMN connectivity to VO motor neurons across abdominal segments A1–A3.**
This Excel file identifies premotor neuron (PMN) inputs to tonic VO motor neurons (MNs) in abdominal segments A1, A2, and A3, with one worksheet per segment. Rows correspond to presynaptic PMNs, and columns correspond to individual VO MNs, including MN15/16 and MN15–17 in each hemisegment. Values indicate the percentage of total synaptic input received by each VO MN that is contributed by the PMN in the corresponding row. The final column reports the mean input across the VO MNs shown in that worksheet. This table shows that A06c and A27h are among the largest and most consistent PMN inputs to VO MNs across segments.

**Table S2. Output specificity of major pre-VO PMNs across the motor neuron pool.**
This table shows synaptic input from major pre-VO PMNs to all annotated MNs in the analyzed segment. Rows correspond to PMNs, and columns correspond to individual MNs. Values indicate the percentage of total synaptic input received by each MN that is contributed by the PMN in the corresponding row. This table is intended to show whether PMNs that contact VO MNs are VO-biased or also provide substantial input to other motor outputs. A06c input is strongly enriched on VO MNs, whereas A27h and other PMNs have broader output patterns across the MN pool.

**Table S3. Synapse number and synaptic fraction for A06c, A27h, and A18b3 outputs to MNs across A1–A3.**
This Excel file lists MNs receiving synaptic input from A06c, A27h, or A18b3 across abdominal segments A1–A3. Separate worksheets are provided for each PMN and segment. For each worksheet, the upper block reports synapse number, and the lower block reports synaptic fraction, calculated as the number of synapses from the indicated PMN divided by the total synaptic input received by the postsynaptic MN. This table clarifies that A06c and A27h provide strong inputs to VO MNs, whereas A18b3 contacts VL MNs and other MN classes.

**Table S4. Tri-segmental connectivity among A06c, A18b3, and A27h PMNs.**
This table reports PMN-to-PMN synaptic connectivity among A06c, A18b3, and A27h across abdominal segments A1–A3. Rows correspond to presynaptic PMNs, and columns correspond to postsynaptic PMNs. Values indicate the percentage of total synaptic input received by each postsynaptic PMN that is contributed by the presynaptic PMN in the corresponding row, calculated within each hemisegment. This table supports the proposed tri-segmental motif in which A27h and A18b3 provide segmentally organized input to A06c, linking excitatory premotor activity to the timing of A06c-mediated inhibition.

**Table S5. PMN inputs to MN26 compared with VO motor neurons.**
This Excel file contains two worksheets comparing PMN inputs to MN26 and VO MNs. Sheet 1 reports PMN inputs to MN26, and sheet 2 reports PMN inputs to VO MNs. Rows correspond to PMNs, and columns correspond to individual MN26 or VO MNs. Values indicate the percentage of total synaptic input received by each MN that is contributed by the PMN in the corresponding row. This comparison supports the use of M26 as a reference muscle in timing analyses: A06c and A18b3 provide no direct synaptic input to MN26, and A27h contributes only small input to MN26, whereas A06c and A27h provide stronger inputs to VO MNs.

| Driver line | Target MNs |
|---|---|
| NB7-1-Gal4 | Tonic MNs:<br>**MN15-17 (VO MN)**, MN9, MN10, MN2, MN3, MN4 |
| FD4-Gal4 | Tonic MNs:<br>**MN15-17 (VO-MN)**, MN9, MN10, MN2, MN3, MN4 |
| vGlut$^{AD}$∩Nkx6$^{DBD}$ | Tonic MNs:<br>**MN15-17 (VO MN)**, **MN15/16 (VO MN)**, MN12, MN13, MN6/7, MN14, MN30, MN28, MN11, MN19, MN20 |
| HB9-Gal4 | Tonic MNs: MN12, MN13, MN6/7, MN14, MN30, MN28, MN11, MN19, MN20 |
| CQ-Gal4 | Tonic MNs: MN9, MN10, MN2, MN3, MN4 |
| 27E09-Gal4 | Phasic MNs: MNISN (RP2), MNISNb/d (RP5) |

**Table S6. Motor neurons targeted by Gal4 drivers used in this study.**
This table summarizes the larval motor neurons (MNs) labeled or manipulated by each Gal4 or split-Gal4 driver used in the study. For each driver, the corresponding MN identities were determined based on anatomical characterization and/or prior published descriptions.
